# Beyond Linear Paths: Generalized Interpolation for Flow-Based Generative Modeling

**DOI:** 10.64898/2026.11.16.767201

**Authors:** B E Nagesh, Jagannath Mondal

## Abstract

Flow-based generative models learn to transform random noise into molecular conformations by following a prescribed interpolation path between the two. Straight-line (linear) interpolation is the usual choice, but it is only one of many possible paths. We compare four paths within a single affine framework, ***x***_*t*_ = *α*_*t*_***x*** + *β*_*t*_***z***: linear, trigonometric, spherical and a new operator-based path in which each degree of freedom follows its own schedule. Linear interpolation is a limiting case of both the operator-based and spherical paths. The operator-based path bends inside it (contracted), whereas the trigonometric and spherical paths bend outside it (expanded). We train the same Transformer velocity field with each path on a three-well potential, the mini-protein Trp-cage and the intrinsically disordered protein *α*-synuclein. All four paths reproduce the reference distributions with comparable accuracy, but they get there differently. The trigonometric and spherical paths resolve conformational basins early and approach the reference smoothly. The operator-based path delays part of the transport, so some features stall or briefly diverge before converging sharply. An exact relation between the rate of change of the KL divergence and the alignment of the learned velocity with the score difference, together with closed-form moments of each path, explains these differences. The choice of interpolation path therefore controls how a flow model converges, not only where it ends.

**FOR TABLE OF CONTENTS ONLY:** 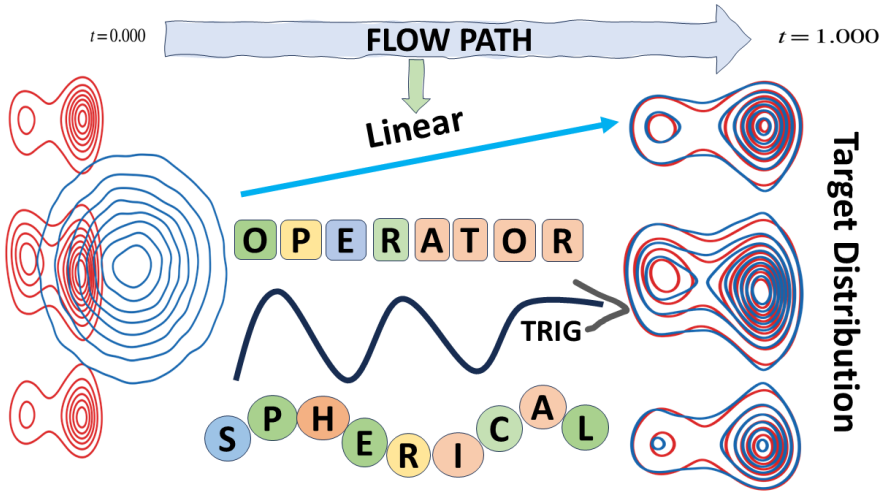

## I. INTRODUCTION

Molecular dynamics (MD) simulations sample conformational space by integrating Newton’s equations, but rugged landscapes with many metastable minima make converged sampling expensive [1–3]. Deep generative models offer a complementary route: once trained on a limited set of configurations, they draw new, statistically consistent samples at negligible cost [4, 5]. Denoising diffusion and flow-based models have been used to enhance conformational sampling [6–11]. However, how these models transport probability mass, that is, when and how conformational states emerge along the generative trajectory, has received far less attention than the quality of their final samples. In earlier work we examined these sampling dynamics for diffusion and rectified-flow models and their dependence on network architecture [12]. Here we ask how the *interpolation path*, the central design choice of a flow model, shapes those dynamics.

A flow model transports samples ***z*** ~ *p*_0_ (a standard Gaussian) to data ***x*** ~ *p*_1_ by integrating a learned, time-dependent velocity field ***v***_*θ*_(***x***, *t*) from *t* = 0 to *t* = 1 (Fig. 1) [13– 17]. In flow matching, the network is trained to reproduce the velocity of a prescribed interpolant between paired source and data samples. For the paths considered here, the interpolant takes the affine form

**FIG 1.**
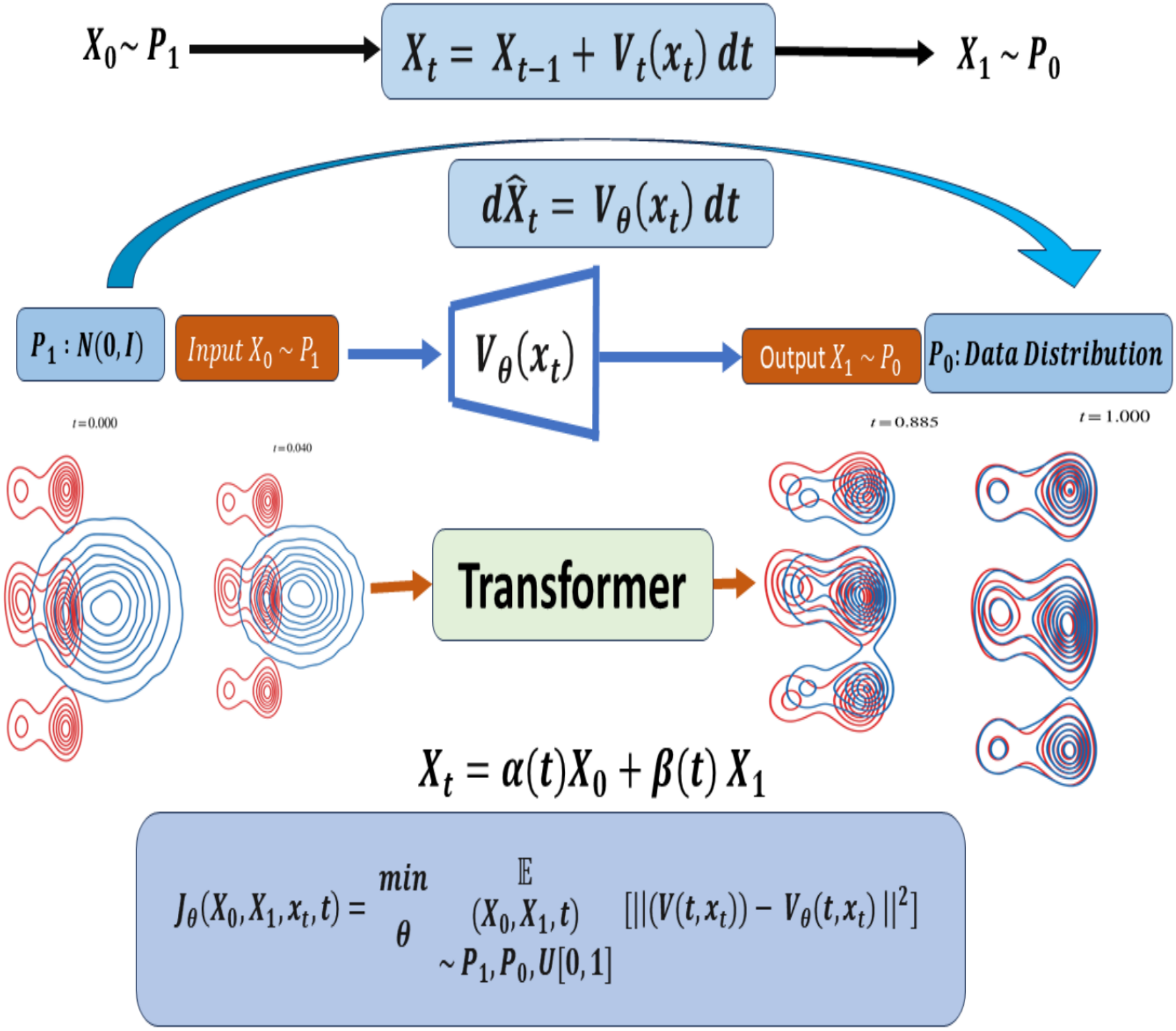
Flow-based generative model. A learned velocity field ***v***_*θ*_(***x***, *t*) transports samples from a Gaussian base distribution *p*_0_ to the data distribution *p*_1_ through an ordinary differential equation. During training, intermediate states are constructed with an interpolation scheme ***x***_*t*_ = *α*_*t*_***x*** + *β*_*t*_***z***, and a Transformer is trained to regress the corresponding conditional velocity with a mean-squared-error loss. At generation, the learned field is integrated from *t* = 0 to *t* = 1.

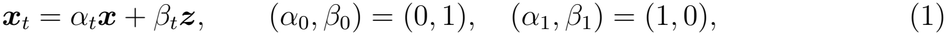

so a path is fully specified by its data coefficient *α*_*t*_ and noise coefficient *β*_*t*_. The common choice is linear interpolation, (*α*_*t*_, *β*_*t*_) = (*t*, 1 − *t*). Curved alternatives have been proposed in several contexts [18–23], but how they change the course of convergence has not been compared systematically, particularly for molecular data.

We study four paths, sketched in Fig. 2. The *linear* path moves each sample along a straight line at constant speed. The *operator-based* path, which we introduce, generalises it: each feature *i* follows its own schedule 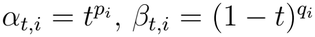, so different degrees of freedom are released at different times (Fig. 2A). The *spherical* path (SLERP) follows the arc between the two endpoint vectors, and the *trigonometric* path, 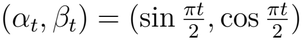, is the special case of an orthogonal arc (Fig. 2B). The four are therefore not independent alternatives. Linear interpolation is a limit of both the operator-based and the spherical constructions. The operator-based path bends inside it, and the spherical and trigonometric paths bend outside it (Sec. II C). The snapshots in Fig. 2 already show that the four paths pass through visibly different intermediate distributions before reaching the same endpoint. We train the same Transformer velocity field with each path on three systems of increasing dimension and complexity: a two-dimensional three-well potential, the folded mini-protein Trp-cage (38 backbone dihedrals) and the intrinsically disordered protein *α*-synuclein (60 dihedrals). Besides the final accuracy, we follow the KL divergence to the reference and the first two moments along the flow. We interpret them using two exact relations: a rate equation linking *dD*_KL_*/dt* to the velocity–score interaction, and the closed-form moments of each interpolant. All four paths reach comparable final accuracy. Their convergence, however, differs in a way that follows directly from where each path sits relative to the linear one.

**FIG 2.**
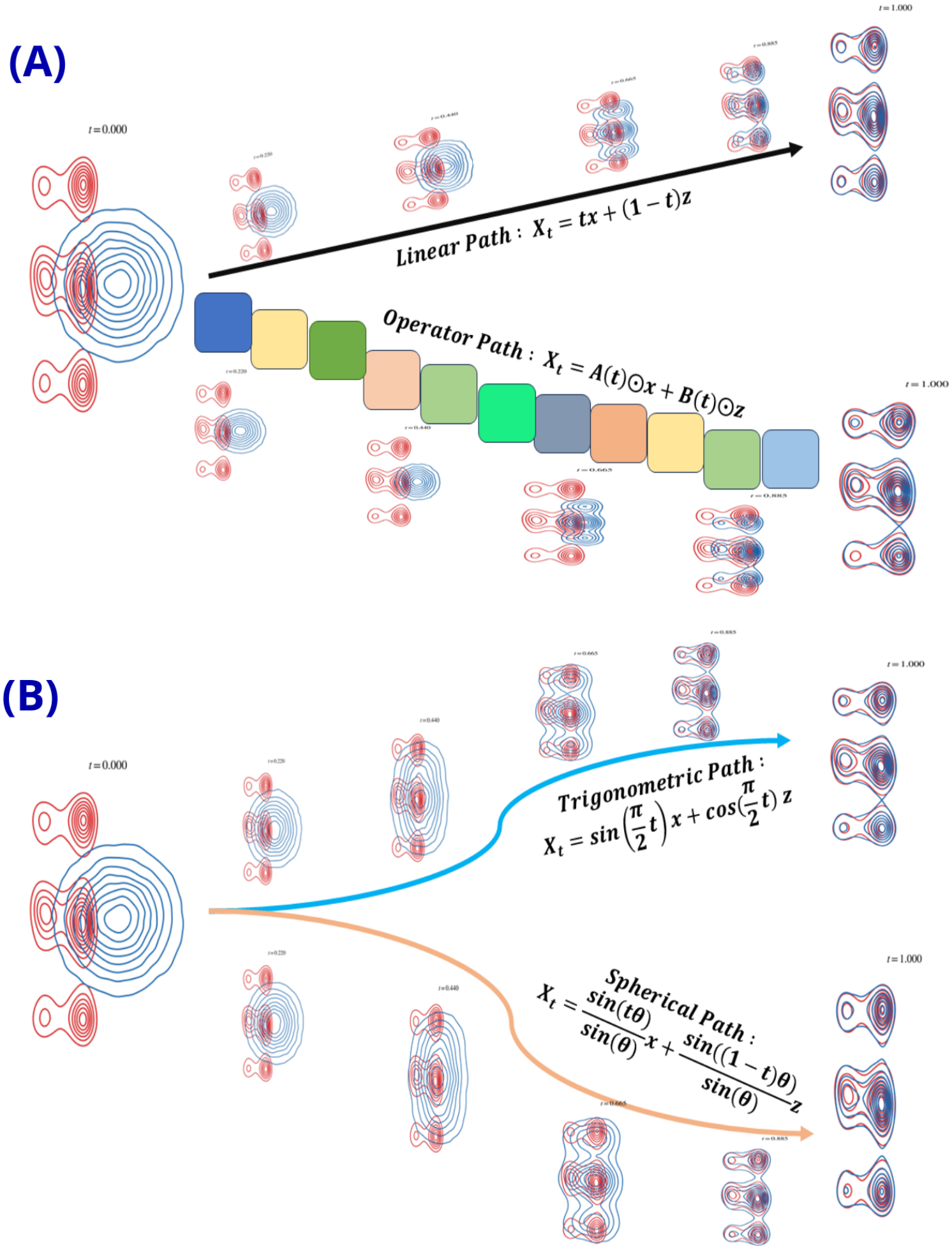
The four interpolation paths. (A) Linear and operator-based paths. The coloured blocks represent the feature-wise coefficient operators **A**(*t*) and **B**(*t*) of the operator-based path. (B) Trigonometric and spherical paths. For each path, the generated (*ψ*_11_, *ϕ*_11_) marginal of Trp-cage (blue) is overlaid on the reference (red) at intermediate flow times. All paths reach the same endpoint through different intermediate distributions.

## II. THEORY

### A. Conditional flow matching

Let ***z*** ~ *p*_0_ and ***x*** ~ *p*_1_, both in ℝ^*d*^. A conditional path ***x***_*t*_ = Ψ_*t*_(***x, z***) with Ψ_0_ = ***z*** and Ψ_1_ = ***x*** defines the conditional velocity ***u***_*t*_(***x, z***) = *∂*_*t*_Ψ_*t*_(***x, z***). Averaged over endpoint pairs, it induces a marginal density *p*_*t*_ and a marginal velocity ***u***_*t*_(***x***_*t*_) = *E*[***u***_*t*_(***x, z***) | ***x***_*t*_] that transports *p*_0_ to *p*_1_. Conditional flow matching trains a network ***v***_*θ*_ by regression onto the conditional velocity,

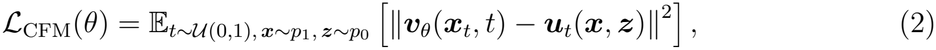

whose minimiser is the marginal velocity. Samples are generated by integrating

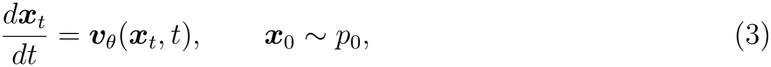

from *t* = 0 to *t* = 1. For the affine paths of Eq. (1), the conditional velocity is

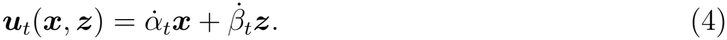

### B. Four interpolation paths

Table I lists the coefficients of the four paths, and Figs. S1 and S2 plot them.

#### Linear

***x***_*t*_ = *t* ***x*** + (1 − *t*)***z*** has the constant conditional velocity ***u***_*t*_ = ***x*** − ***z*** and serves as the reference path.

#### Trigonometric

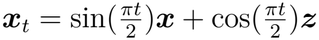, with

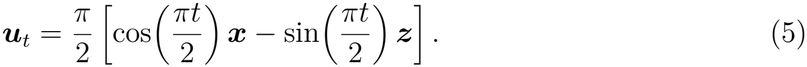

#### Operator-based

Each feature follows its own power-law schedule,

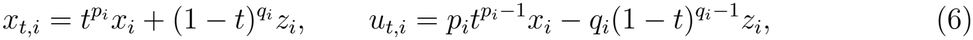

i.e. ***x***_*t*_ = ***α***_*t*_ ⊙ ***x*** + ***β***_*t*_ ⊙ ***z*** with element-wise coefficients. The exponents vary linearly across features,

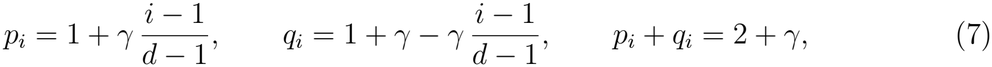

so the first feature has (*p, q*) = (1, 1 + *γ*) and the last has (1 + *γ*, 1). We use *γ* = 1, so the exponents range from 1 to 2. This is a prescribed schedule, not an optimised one.

**TABLE I.**
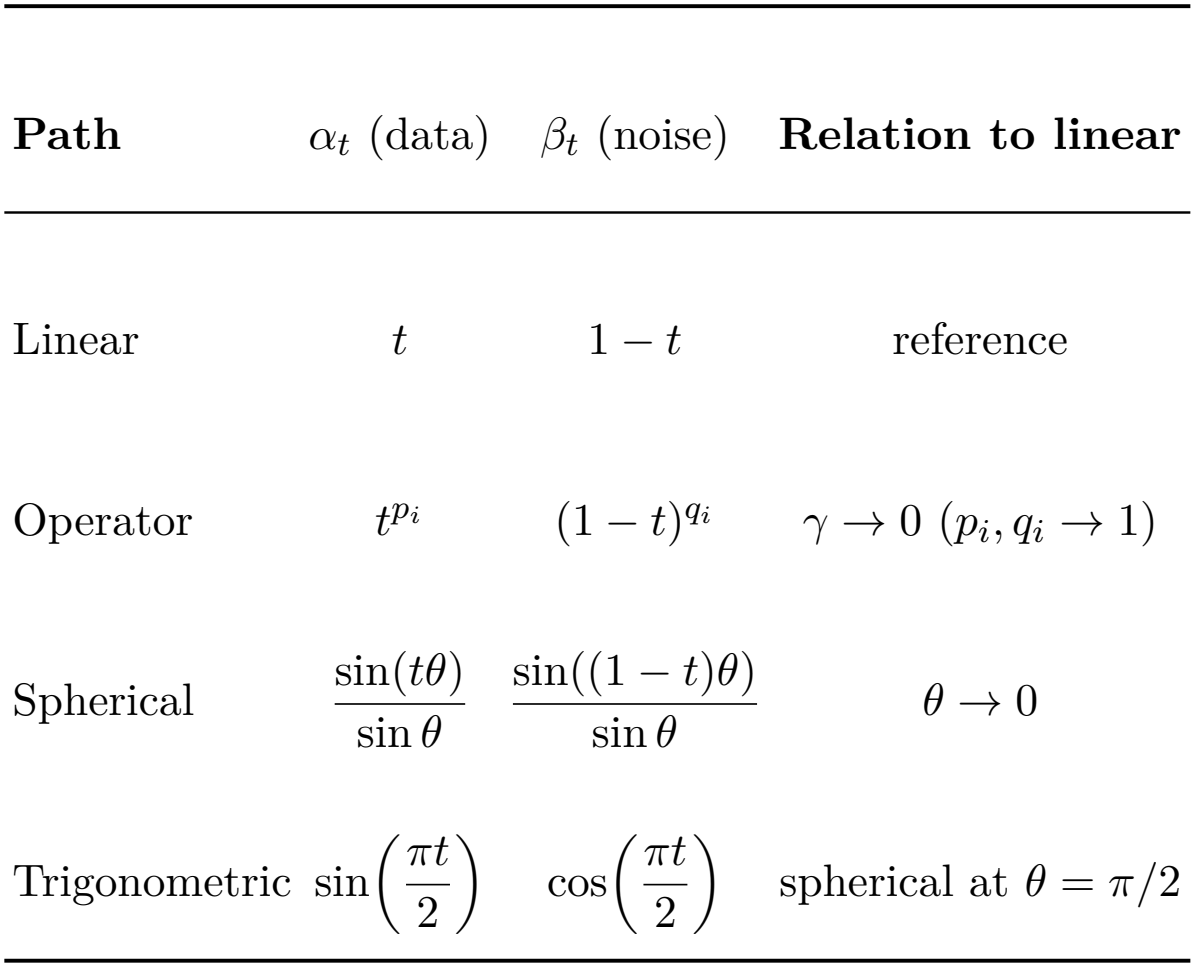
Interpolation coefficients of the four paths and their relation to the linear path.

| **Path** | α<sub>*t*</sub> (data) | β<sub>*t*</sub> (noise) | **Relation to linear** |
|---|---|---|---|
| Linear | *t* | 1 − *t* | reference |
| Operator | *t*<sup>*p*<sub>*i*</sub></sup> | (1 − *t*)<sup>*q*<sub>*i*</sub></sup> | γ → 0 (*p*<sub>*i*</sub>, *q*<sub>*i*</sub> → 1) |
| Spherical | $\frac{\sin(t\theta)}{\sin\theta}$ | $\frac{\sin((1-t)\theta)}{\sin\theta}$ | θ → 0 |
| Trigonometric | $\sin\left(\frac{\pi t}{2}\right)$ | $\cos\left(\frac{\pi t}{2}\right)$ | spherical at θ = π/2 |

#### Spherical

With 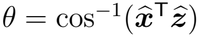 the angle between the normalised endpoints,

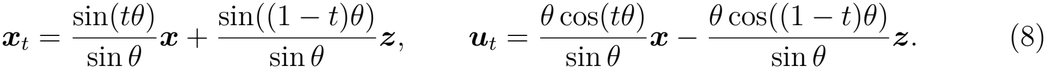

For equal-norm endpoints this follows the great-circle arc between them and preserves their norm.

### C. How the paths are related

#### Limits

Setting *γ* = 0 in Eq. (7) gives *p*_*i*_ = *q*_*i*_ = 1 and recovers the linear path. As *θ* → 0, sin(*tθ*)*/* sin *θ* → *t* and sin((1 − *t*)*θ*)*/* sin *θ* → 1 − *t*, so SLERP also reduces to the linear path (Fig. S3). For orthogonal, equal-norm endpoints, *θ* = *π/*2 and Eq. (8) becomes the trigonometric path.

#### Bracketing

The paths are also ordered relative to the linear one. Since *p*_*i*_, *q*_*i*_ ≥ 1 and sin is concave on [0, *π*], so that sin(*tθ*) ≥ *t* sin *θ* for *t* ∈ [0, 1], the following holds for every *t* ∈ [0, 1]:

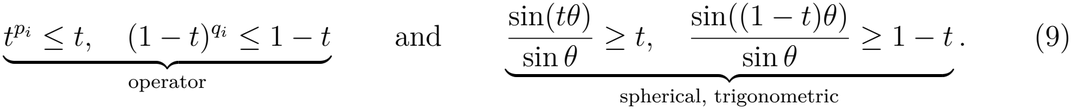

Consequently *α*_*t*_ + *β*_*t*_ ≤ 1 for the operator-based path, = 1 for the linear path and ≥ 1 for the spherical and trigonometric paths. The linear path is thus the boundary between a *contracted* family (operator-based, indexed by *γ*) and an *expanded* family (spherical, indexed by *θ*, with trigonometric at *θ* = *π/*2). The early slopes differ in the same way. On the trigonometric path, data enter at rate 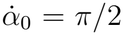 while the noise coefficient is still flat 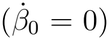. On the operator-based path, features with *p*_*i*_ *>* 1 have 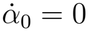: their data component is deferred to late flow times. The results below show that this single ordering accounts for most of the observed differences.

## III. SYSTEMS AND MODEL

### Systems

(i) A particle in a two-dimensional three-well potential, simulated in our group [24]; the reference contains 5 × 10^5^ samples. (ii) Trp-cage, represented by 19 backbone (*ψ, ϕ*) pairs (*d* = 38), **X**_Trp_ = [*ψ*_1_, *ϕ*_1_, …, *ψ*_19_, *ϕ*_19_], from a 100 *μ*s a99SB-disp trajectory generated by D. E. Shaw Research on Anton [25]. (iii) *α*-Synuclein, represented by 30 (*ψ, ϕ*) pairs (*d* = 60), from a 73 *μ*s a99SB-disp trajectory, also generated by D. E. Shaw Research [25, 26]. To create a data-limited regime, models are trained on a random subset of the reference data: 10% for the three-well potential and Trp-cage, and 50% for *α*-synuclein.

### Velocity network and training

Following our earlier finding that flow models learn molecular velocity fields best with a Transformer [12], all four paths use the same time-conditioned Transformer [27], *f*_*θ*_ : ℝ^*d*^ × [0, 1] → ℝ^*d*^. It has hidden dimension 256, six layers with eight attention heads and dropout 0.2. Models are trained for 5 × 10^4^ epochs with batch size 4096 and AdamW (learning rate 5 × 10^−5^, weight decay 10^−4^) [28, 29] in PyTorch [30]. Accuracy is measured by the KL divergence [31] between generated and reference (marginal) distributions, and convergence by following the same quantity, and the moments, along the flow.

## IV. RESULTS

### A. Three-well potential

The training subset (Fig. 3a) samples the barrier regions of the reference landscape (Fig. 3b) only sparsely. All four paths recover the full three-basin topology, including the barrier regions (Fig. 3c–f). Their final KL divergences lie between 0.013 and 0.017 (Fig. 3g).

**FIG 3.**
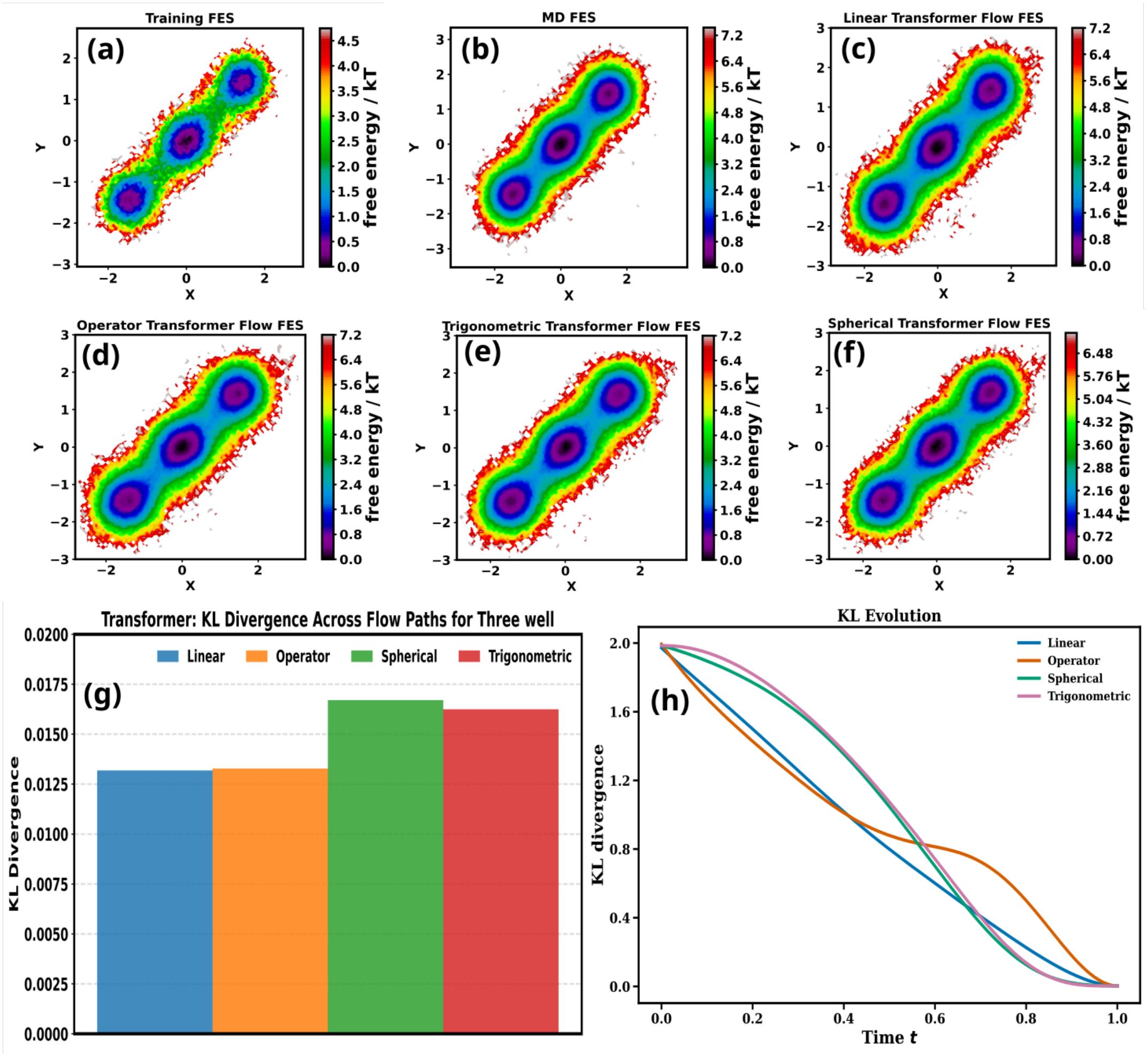
Three-well potential. (a) Free-energy surface (FES) of the 10% training subset and (b) of the full reference. (c)–(f) FES generated with the linear, operator-based, trigonometric and spherical paths. (g) Final KL divergence to the reference. (h) KL divergence along the flow.

The linear and operator-based paths are marginally lower, but the differences are negligible on this scale.

The KL trajectories along the flow (Fig. 3h) are, in contrast, clearly different. The linear path decreases almost uniformly. The trigonometric and spherical paths are S-shaped: slow at first, fastest near *t* ≈ 0.6, then levelling off. The operator-based path drops fastest initially, stalls at *D* ≈ 0.8 around *t* ≈ 0.55–0.7, and then falls sharply.

#### Rate of KL change

To see where these shapes come from, we write the generated density *p*_*t*_ under Eq. (3) as obeying the continuity equation *∂*_*t*_*p*_*t*_+∇·(*p*_*t*_***v***_*θ*_) = 0. Differentiating *D*(*t*) = *D*_KL_(*p*_*t*_∥*p*_ref_) = *p*_*t*_ log(*p*_*t*_*/p*_ref_) *d****x***, substituting the continuity equation and integrating by parts (with vanishing boundary terms) gives

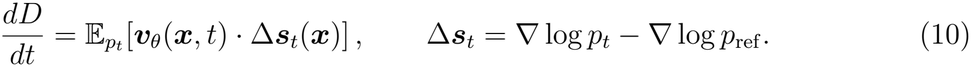

KL therefore decreases when the transport velocity is anti-aligned with the score difference, increases when it is aligned, and plateaus when the two are nearly orthogonal on average. Nothing requires *D*(*t*) to be monotonic. Equation (10) involves the *learned* field, so it reflects both the path and the network’s approximation of it. Because the dot product is a sum over coordinates, the rate decomposes into per-coordinate contributions, *dD/dt* = Σ_*k*_ *A*_*k*_(*t*) with 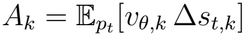.

The interaction evaluated directly (Fig. 4a) matches the finite-difference derivative of the KL curves (Fig. 4b), which validates Eq. (10) numerically. The interaction stays negative throughout for every path, so all four paths decrease KL monotonically here. Its time profile reproduces each KL shape. The linear path has a nearly constant rate. The trigonometric and spherical paths start close to zero and reach their most negative rate at *t* ≈ 0.6, which gives the S-shape. The operator-based path is most negative at *t* = 0, nearly vanishes at *t* ≈ 0.6 (the plateau) and has a second minimum at *t* ≈ 0.85 (the late drop).

**FIG 4.**
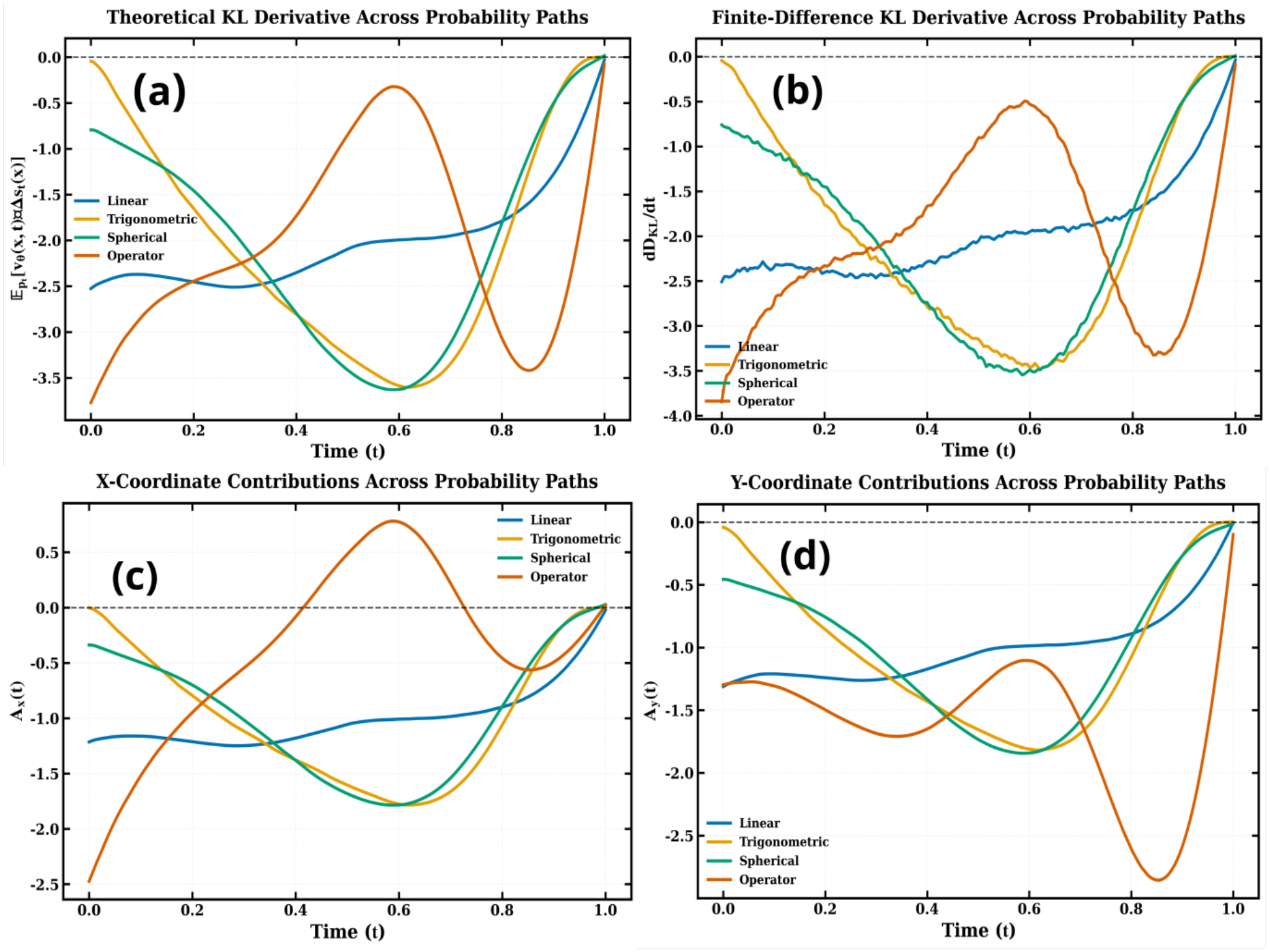
Velocity–score interaction for the three-well potential. (a) 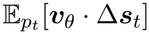 evaluated along the generated trajectories. (b) Finite-difference estimate of *dD/dt* from Fig. 3h. (c,d) Contributions *A*_*x*_(*t*) and *A*_*y*_(*t*) of the two coordinates.

The coordinate decomposition shows why the operator-based path behaves this way (Fig. 4c,d). With *d* = 2 and *γ* = 1, Eq. (7) gives (*p, q*) = (1, 2) for *x* and (2, 1) for *y*. The *x* coordinate thus loses its noise quickly while its data component enters linearly, whereas *y* receives its data component only late (*α*_*t*_ = *t*^2^). Accordingly, *A*_*x*_ dominates the early decrease, and *A*_*y*_ produces the late drop at *t* ≈ 0.85. In between, *A*_*x*_ turns positive, so the *x* coordinate briefly works against convergence, and the net rate approaches zero. The plateau is therefore a direct consequence of the feature-dependent schedule. For the linear, trigonometric and spherical paths, the two coordinates contribute with similar time profiles.

### B. Trp-cage

For Trp-cage, all four paths reproduce the populated regions of the representative (*ψ*_1_, *ϕ*_1_) marginal (Fig. 5a–f) and of other pairs (Figs. S4–S7). Every marginal KL divergence across the 19 pairs is below 0.08, and no path is systematically better (Fig. 5g).

**FIG 5.**
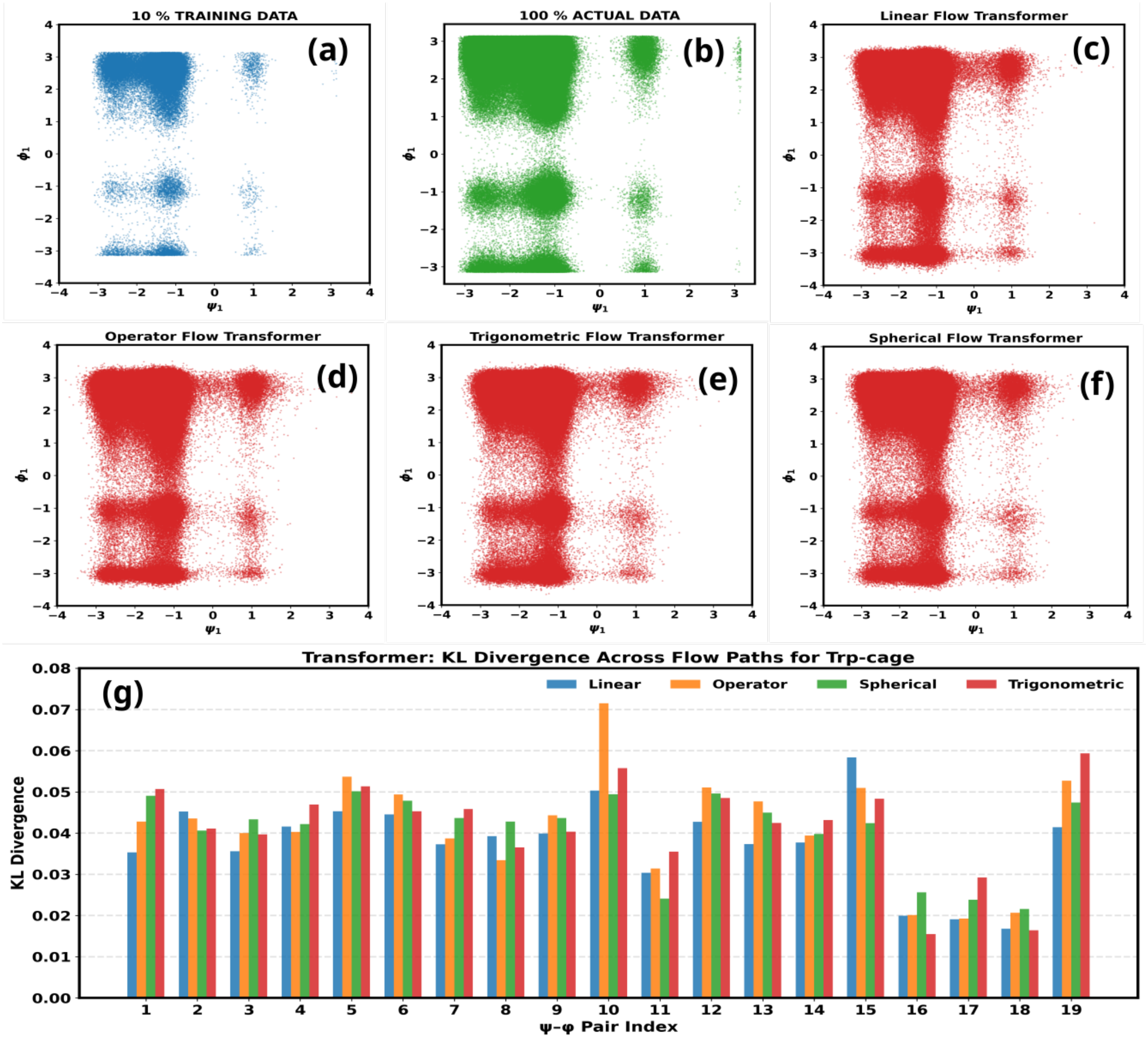
Trp-cage marginal distributions. (a) 10% training subset and (b) full reference for the (*ψ*_1_, *ϕ*_1_) pair. (c)–(f) Samples generated with the linear, operator-based, trigonometric and spherical paths. (g) Marginal KL divergence to the reference for all 19 (*ψ, ϕ*) pairs.

The marginal KL trajectories (Fig. 6) show that convergence is not uniform across degrees of freedom. On the trigonometric and spherical paths, nearly all marginals decrease smoothly and monotonically. On the linear path, most marginals do the same, but a few remain on a plateau before dropping. Delayed marginals are most pronounced for the operator-based path, where several KL curves rise transiently before falling. The pair-averaged KL is given in Fig. S8.

**FIG 6.**
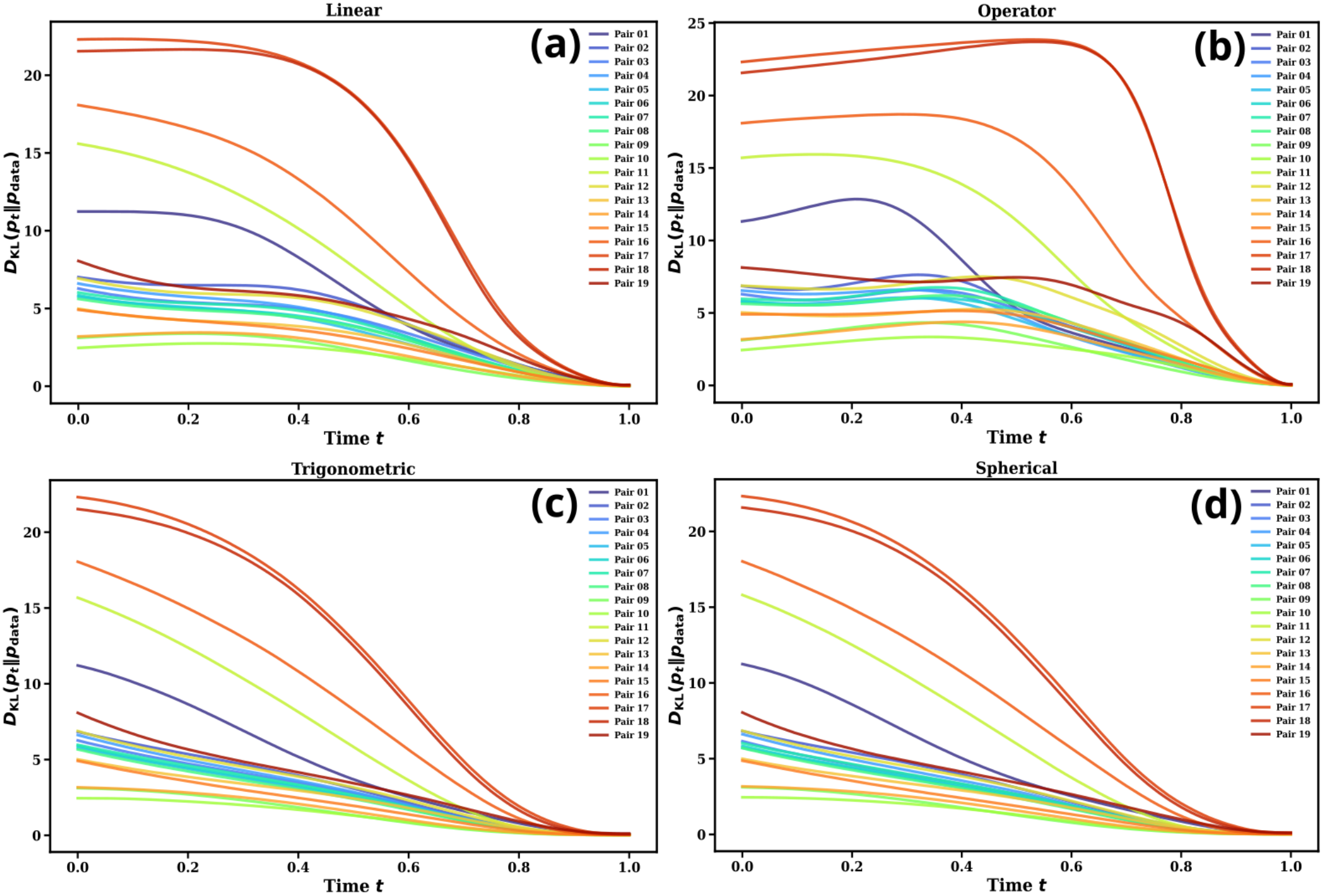
Trp-cage: marginal KL divergence along the flow for the 19 (*ψ, ϕ*) pairs with the (a) linear, (b) operator-based, (c) trigonometric and (d) spherical paths.

Applying Eq. (10) to each marginal accounts for these features (Fig. 7). The finite-difference check and the *ψ* and *ϕ* contributions are given in Figs. S9–S11. Two features of the interaction follow the ordering of Eq. (9). The first is the *timing* of the strongest negative interaction. It is reached at *t* ≈ 0.6 for the trigonometric and spherical paths, *t* ≈ 0.67 for the linear path and *t* ≈ 0.78 for the operator-based path, which is the order in which each path brings in its data component. The second is the *shape*. On the expanded paths the minima are broad and shallow (about −40). On the linear path they are deeper (about −65). On the operator-based path they are narrow and deep (about −125), preceded by an interval in which the interaction is close to zero or slightly positive. Plateaus and transient rises therefore appear where the interaction vanishes or changes sign, and sharp drops follow where it is concentrated. A path that defers transport has to complete it faster later.

**FIG 7.**
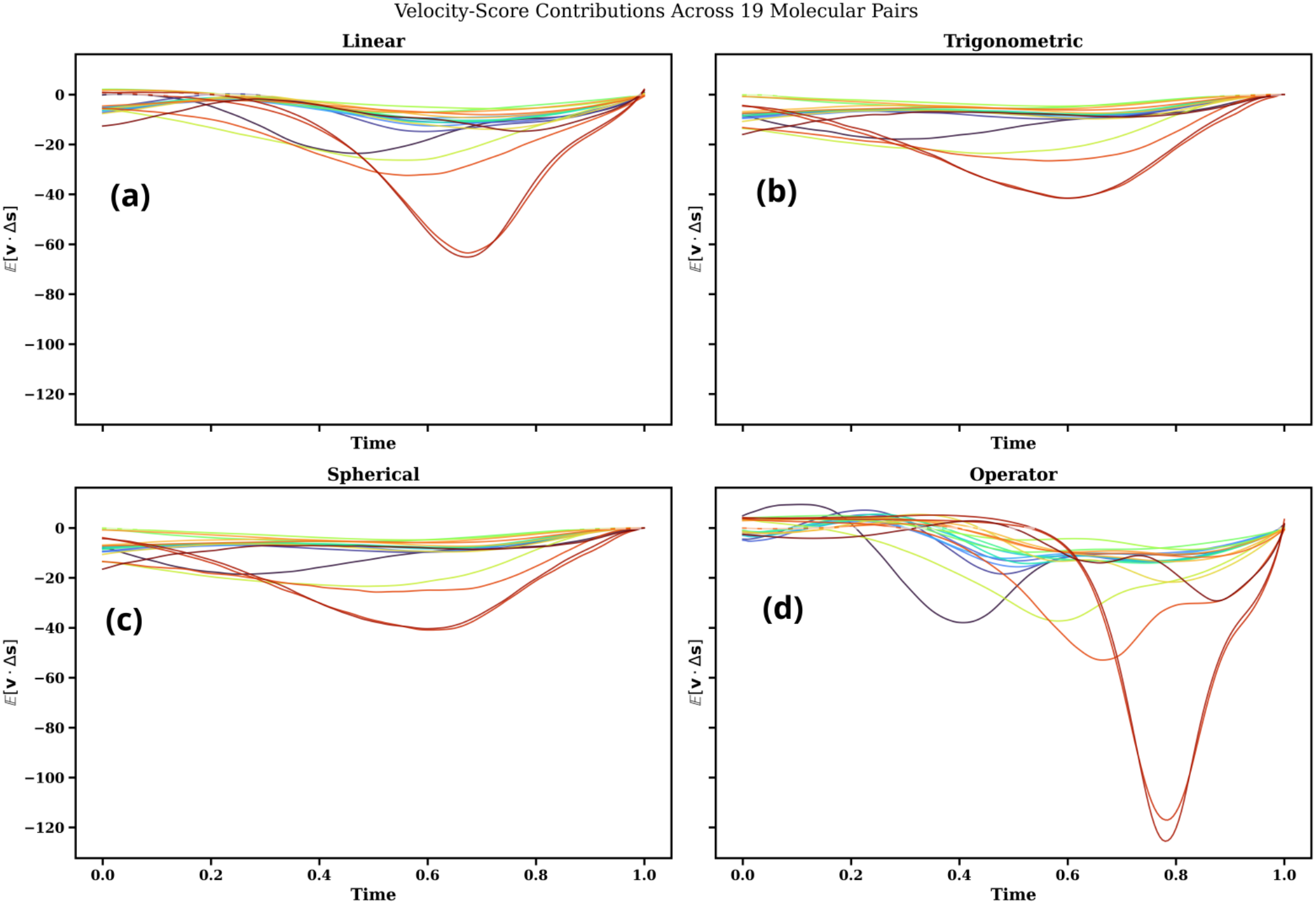
Trp-cage: velocity–score interaction. 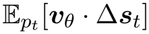 for the 19 (*ψ, ϕ*) pairs with the (a) linear, (b) trigonometric, (c) spherical and (d) operator-based paths.

#### Moments

For an exact flow, the generated marginal *p*_*t*_ equals that of the interpolant. With ***z*** ~ N (0, *I*) independent of ***x***, the mean and variance of feature *i* along any affine path with fixed coefficients follow in closed form:

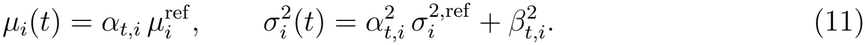

The mean trajectory is therefore the data coefficient itself, scaled by the reference mean. By Eq. (9), it lies below the straight line for the operator-based path and above it for the trigonometric path. The variance dips below its initial value when both coefficients are well below one, and the dip is deepest for the operator-based path. For the spherical path, *θ* varies between sample pairs, so Eq. (11) holds only approximately, but the same inequality applies pair by pair. We compare these predictions with the means and variances estimated from the generated samples,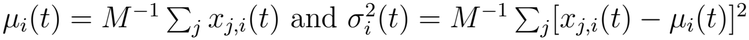.

The generated moments follow Eq. (11) closely. The linear path gives straight-line means (Fig. 8a). The operator-based path gives contracted, convex means (Fig. 8c), and its variances dip lower than on the linear path, because both of its coefficients lie below the linear ones (Fig. 8b,d). The trigonometric and spherical paths give expanded, concave means that approach the reference early (Fig. 9). All paths end at the reference mean and variance (squares). The remaining pairs behave the same way (Figs. S12–S15 and S16–S19).

**FIG 8.**
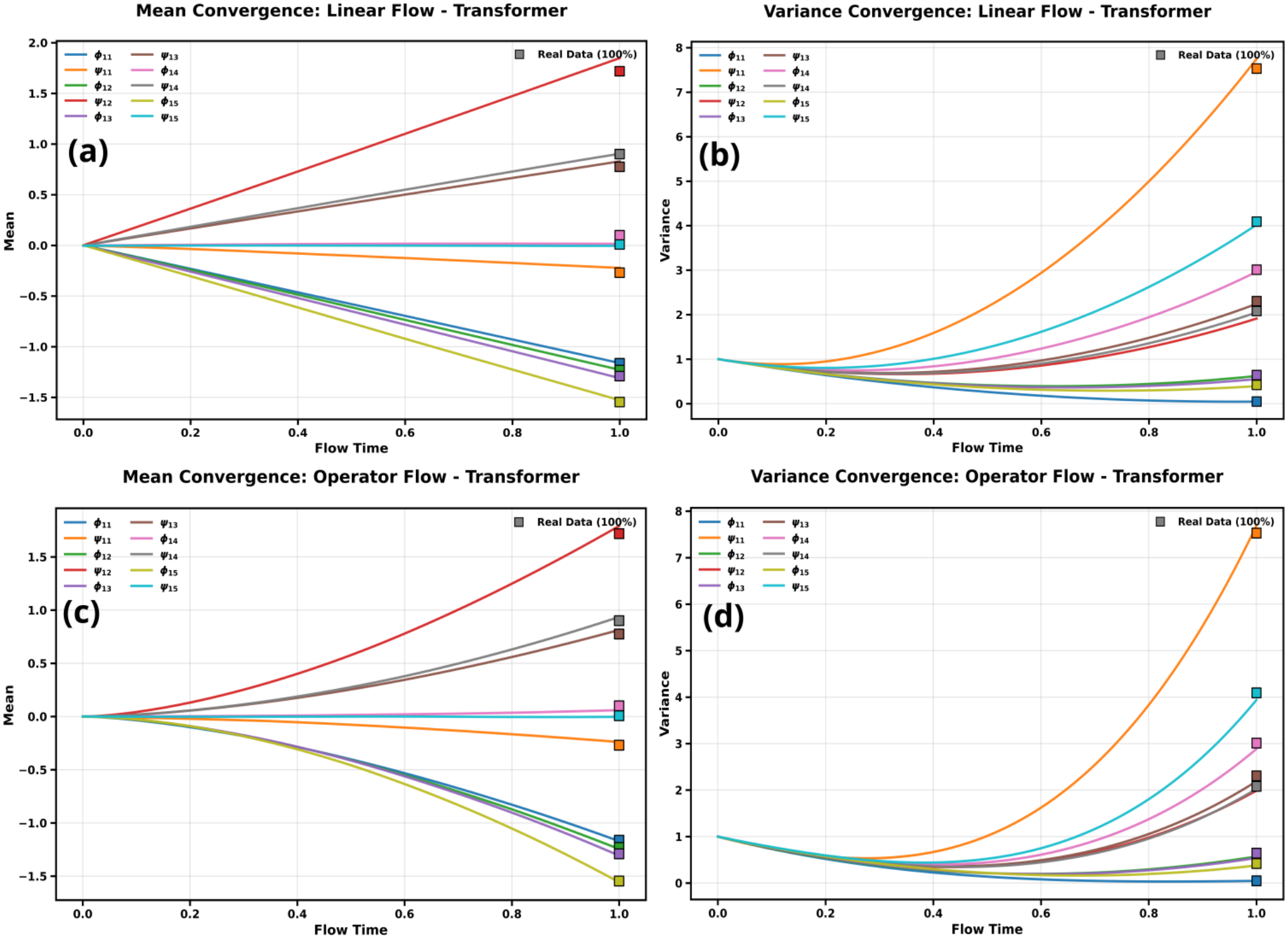
Trp-cage moments, linear and operator-based paths. (pairs 11–15). (a) Mean and (b) variance along the flow for the linear path; (c,d) the same for the operator-based path. Squares mark reference values.

**FIG 9.**
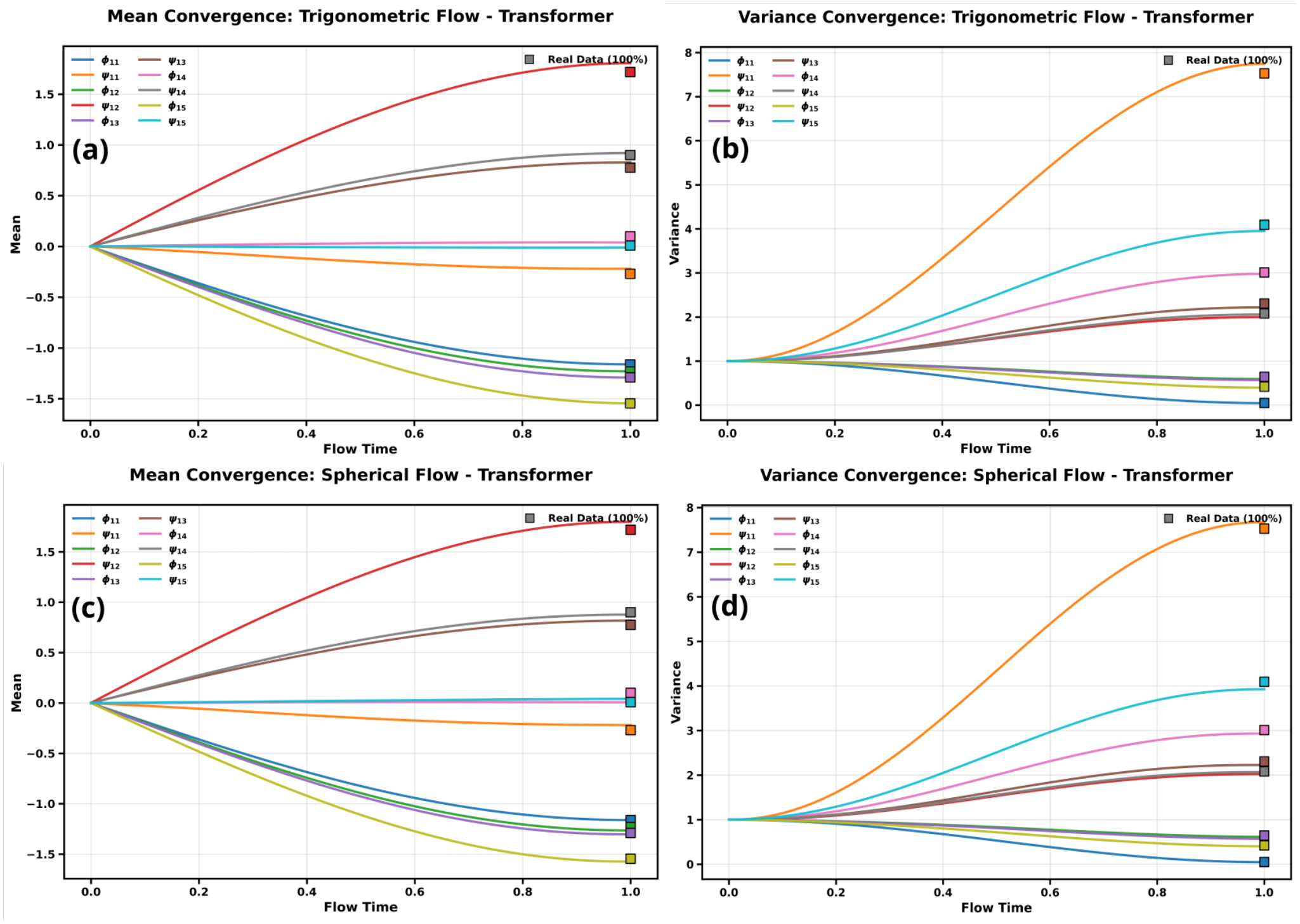
Trp-cage moments, trigonometric and spherical paths. (pairs 11–15). (a) Mean and (b) variance along the flow for the trigonometric path; (c,d) the same for the spherical path. Squares mark reference values.

The density snapshots of the (*ψ*_11_, *ϕ*_11_) marginal show the same ordering directly in the densities (Figs. S20–S22). By *t* ≈ 0.3, two basins are already resolved on the trigonometric and spherical paths, the linear path shows the first signs of structure, and the operator-based path shows none (Fig. S20). By *t* ≈ 0.6, the expanded paths have essentially recovered the reference basins, whereas the operator-based path is still furthest from them (Fig. S21). All four paths agree by *t* = 1 (Fig. S22).

#### C. α-Synuclein

For the 60-dimensional disordered ensemble of *α*-synuclein, all four paths again reproduce the reference marginals (Figs. S23–S27), with marginal KL divergences below 0.16 for all 30 pairs (Fig. S23g). The values are higher than for Trp-cage, consistent with a broader, more heterogeneous ensemble. The convergence behaviour seen for Trp-cage carries over. Marginals converge non-uniformly; delayed relaxation is most evident for the linear and operator-based paths, and on the operator-based path many marginal KL curves rise transiently before converging (Figs. S28 and S29). The velocity–score interaction again agrees with the finite-difference KL derivative (Figs. S30–S33). The moments follow the same contracted and expanded trajectories (Figs. S34–S39 and S40–S45).

## V. DISCUSSION

Across a two-dimensional potential, a folded mini-protein and an intrinsically disor-dered protein, the four interpolation paths reach comparable final accuracy with the same Transformer velocity field. The path determines *how* the model gets there, not *whether* it does.

The four paths fit a single picture (Table II). All four are affine interpolants, and linear interpolation is their common limit. Relative to it, the operator-based family bends inward (*α*_*t*_ + *β*_*t*_ ≤ 1, with data entry deferred feature by feature), and the spherical family, which includes the trigonometric path, bends outward (*α*_*t*_ + *β*_*t*_ ≥ 1, with data entering early while noise is retained). The observed dynamics follow from this ordering through two exact relations. Equation (11) turns the coefficients directly into mean and variance trajectories, which are contracted, straight or expanded. Equation (10) turns the timing of transport into the shape of the KL curve. Expanded paths spread the velocity–score interaction broadly and early, so basins appear early and KL decreases smoothly. The linear path spreads it roughly uniformly. The operator-based path concentrates it late and, for features with large *p*_*i*_, in a short interval, which produces plateaus or transient rises followed by sharp drops. In the three-well potential, this feature-by-feature release can be seen directly in the coordinate-resolved rates. The near-identity of the spherical and trigonometric results suggests that sampled endpoint pairs lie close to the orthogonal (*θ* ≈ *π/*2) case.

Two caveats apply. First, Eq. (10) involves the learned velocity field, so network approximation error also contributes to the observed rates. The close agreement with the finite-difference derivative and with Eq. (11) indicates that the Transformer reproduces the prescribed paths well, but the decomposition between path and architecture effects is not resolved here. Second, for the protein systems we analyse one- and two-dimensional marginals and the first two moments. Agreement at this level does not by itself establish agreement of the full joint distribution.

The practical message is that the interpolation path is a tunable control over the course of generation. Expanded paths resolve conformational basins early and smoothly. Contracted, feature-dependent paths let selected degrees of freedom be resolved before others, with *γ* setting the degree of staging. When comparing or designing flow models for molecular systems, the full trajectory of convergence, and not only the final distribution, should therefore be examined.

**TABLE II.** Summary: position of each path relative to the linear path and the resulting convergence behaviour.

| **Path** | α<sub>t</sub> + β<sub>t</sub> | **Data entry** | **Mean trajectory** | **KL evolution** |
|---|---|---|---|---|
| Operator (γ > 0) | ≤ 1 | late, feature-dependent | contracted | plateaus or transient rises, then sharp drops |
| Linear (γ = 0, θ → 0) | = 1 | uniform | straight | near-uniform; few delayed marginals |
| Spherical (θ > 0) | ≥ 1 | early | expanded | smooth, S-shaped, monotonic |
| Trigonometric (θ = π/2) | ≥ 1 | early | expanded | smooth, S-shaped, monotonic |

## SUPPORTING INFORMATION

Coefficient schedules and limiting cases (Figs. S1–S3); additional Trp-cage marginals, KL, velocity–score and moment analyses, and density snapshots (Figs. S4–S22); *α*-synuclein results (Figs. S23–S45).

## CODE AVAILABILITY

Code and scripts are available at https://github.com/nagesh123-geek/Generative-Flow-Path.

## ACKNOWLEDGEMENTS

We acknowledge support of the Department of Atomic Energy, Government of India, under Project Identification No. RTI 4007.

