## Supplemental Information for "Beyond Linear Paths: Generalized Interpolation for Flow-Based Generative Modeling"

Nagesh B E* and Jagannath Mondal†

*Tata Institute of Fundamental Research, 36/P,*

*Gopanpally Village, Serilingampally Mandal,*

*Ranga Reddy District, Hyderabad 500046, Telangana, India*

(Dated: November 17, 2026)

### S1. INTERPOLATION COEFFICIENTS AND LIMITING CASES

All four paths have the affine form $\mathbf{x}_t = \alpha_t \mathbf{x} + \beta_t \mathbf{z}$, with $\mathbf{x}$ a data sample and $\mathbf{z}$ a standard Gaussian sample. Below we plot the data coefficient $\alpha_t$ and the noise coefficient $\beta_t$ for each path, together with the limits in which the operator-based and spherical paths reduce to the linear one.

*

†

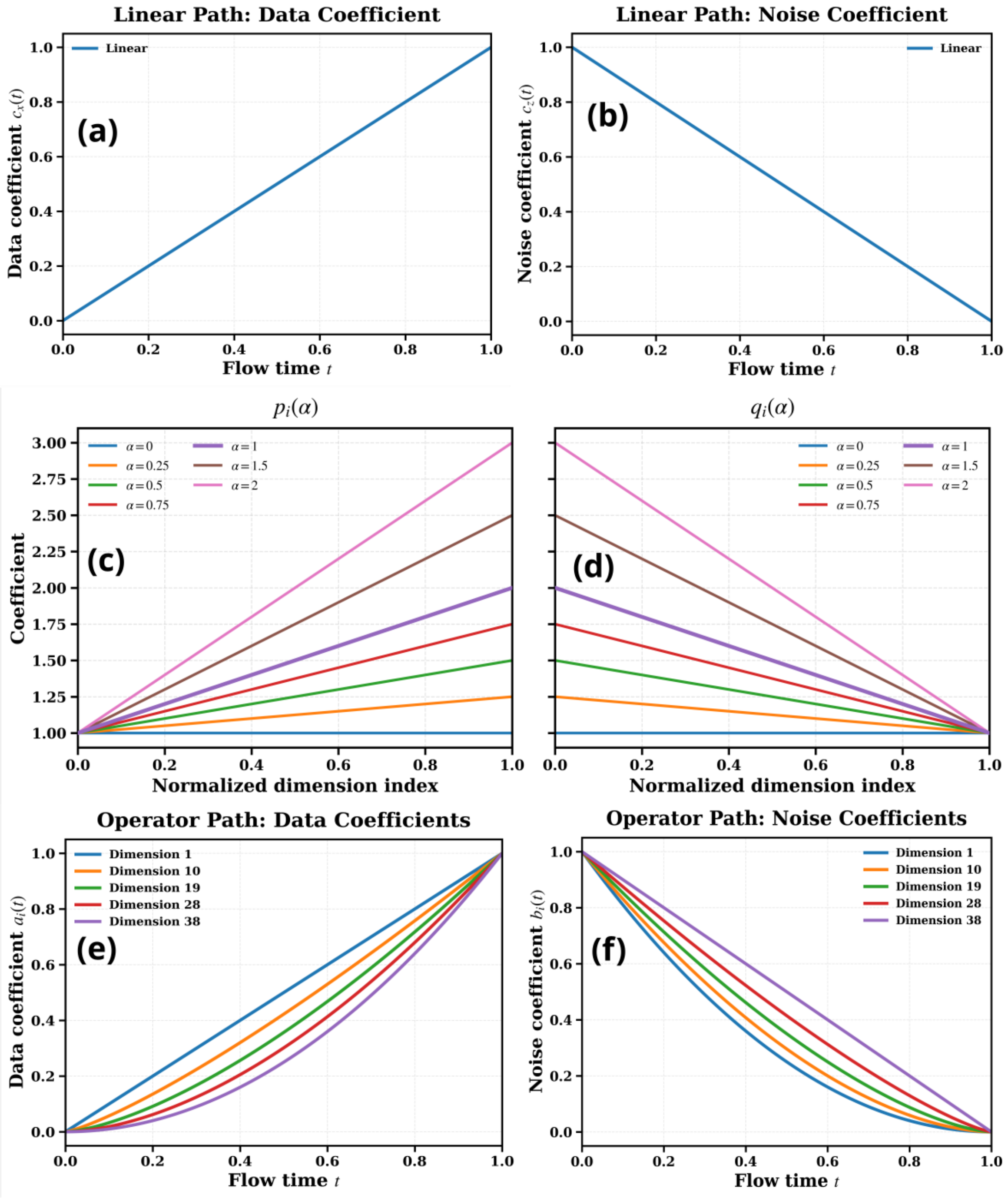


FIG. S1. **Coefficients of the linear and operator-based paths.** (a,b) Data and noise coefficients of the linear path. (c,d) Exponents $p_i$ and $q_i$ as a function of feature index $i$ for several values of the hyperparameter $\gamma$ (labelled $\alpha$ in the panels). (e,f) Data and noise coefficients of the operator-based path ($\gamma = 1$) for five selected features.

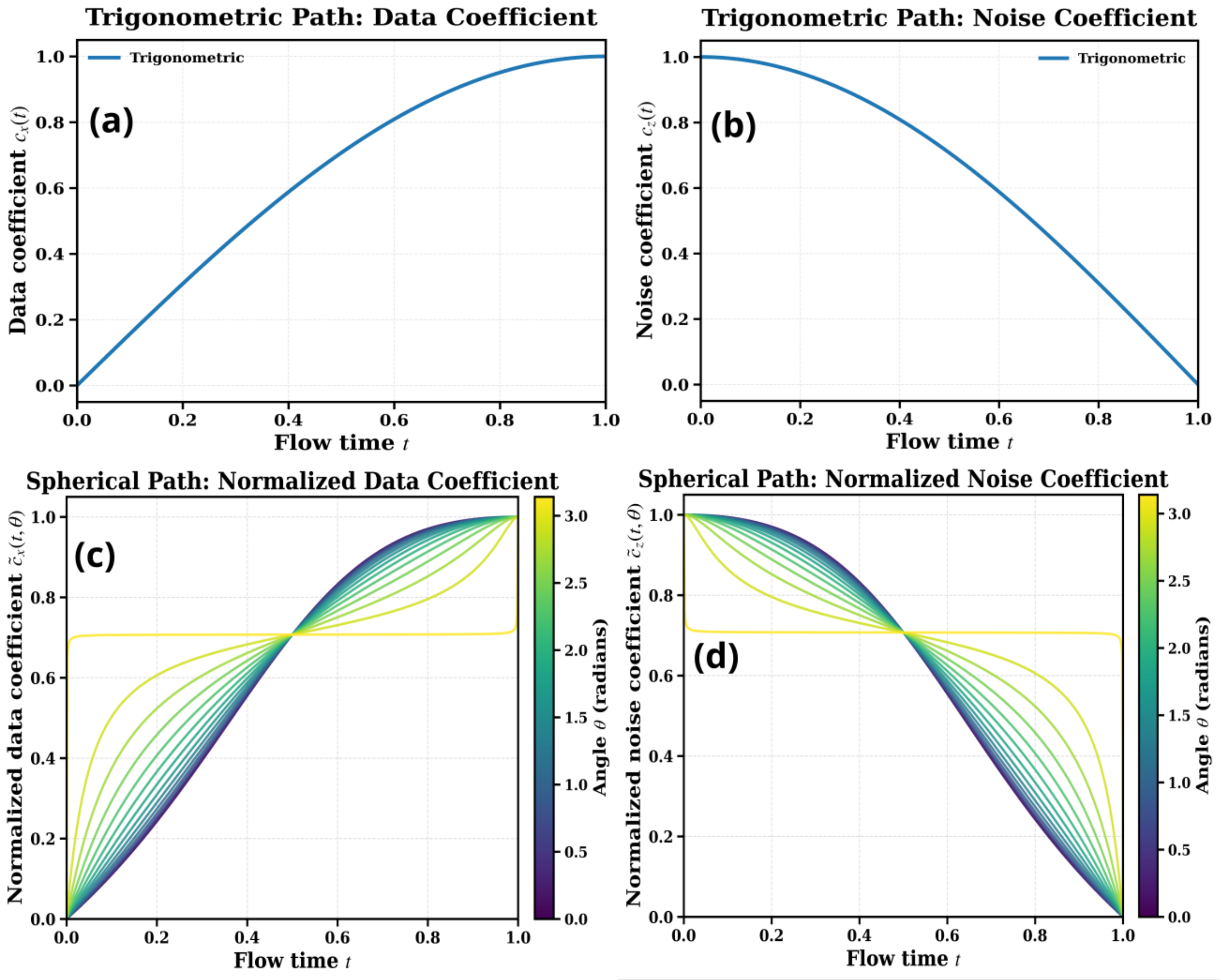


FIG. S2. **Coefficients of the trigonometric and spherical paths.** (a,b) Data and noise coefficients of the trigonometric path. (c,d) Data and noise coefficients of the spherical path for several endpoint angles $\theta$.

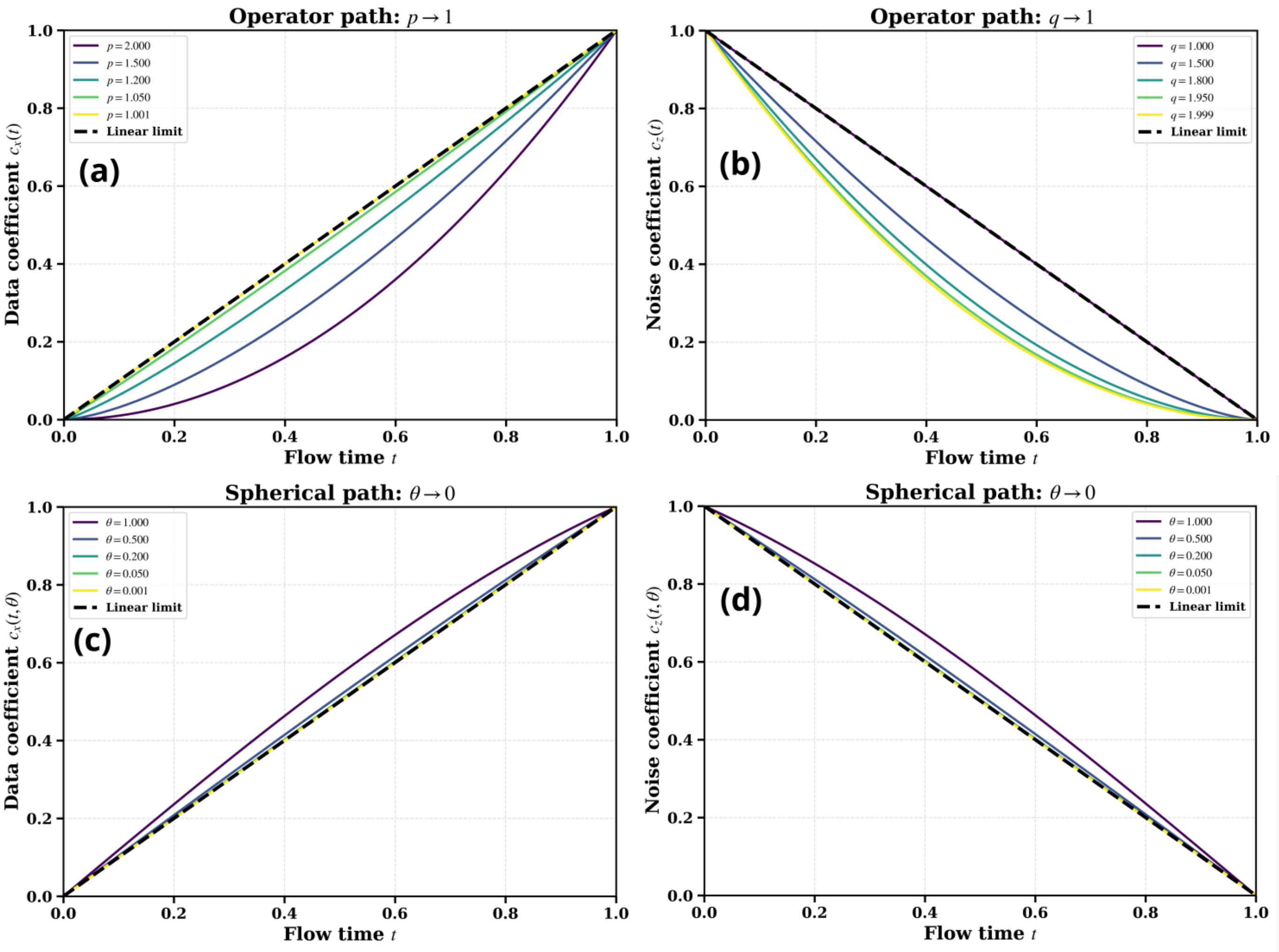


FIG. S3. **Linear interpolation as a limiting case.** (a,b) Operator-based path: as $\gamma \to 0$ the exponents $p_i, q_i \to 1$ and the coefficients approach $(t,\, 1-t)$. (c,d) Spherical path: as $\theta \to 0$ the coefficients approach $(t,\, 1-t)$.

### S2. TRP-CAGE

#### A. Additional marginal distributions

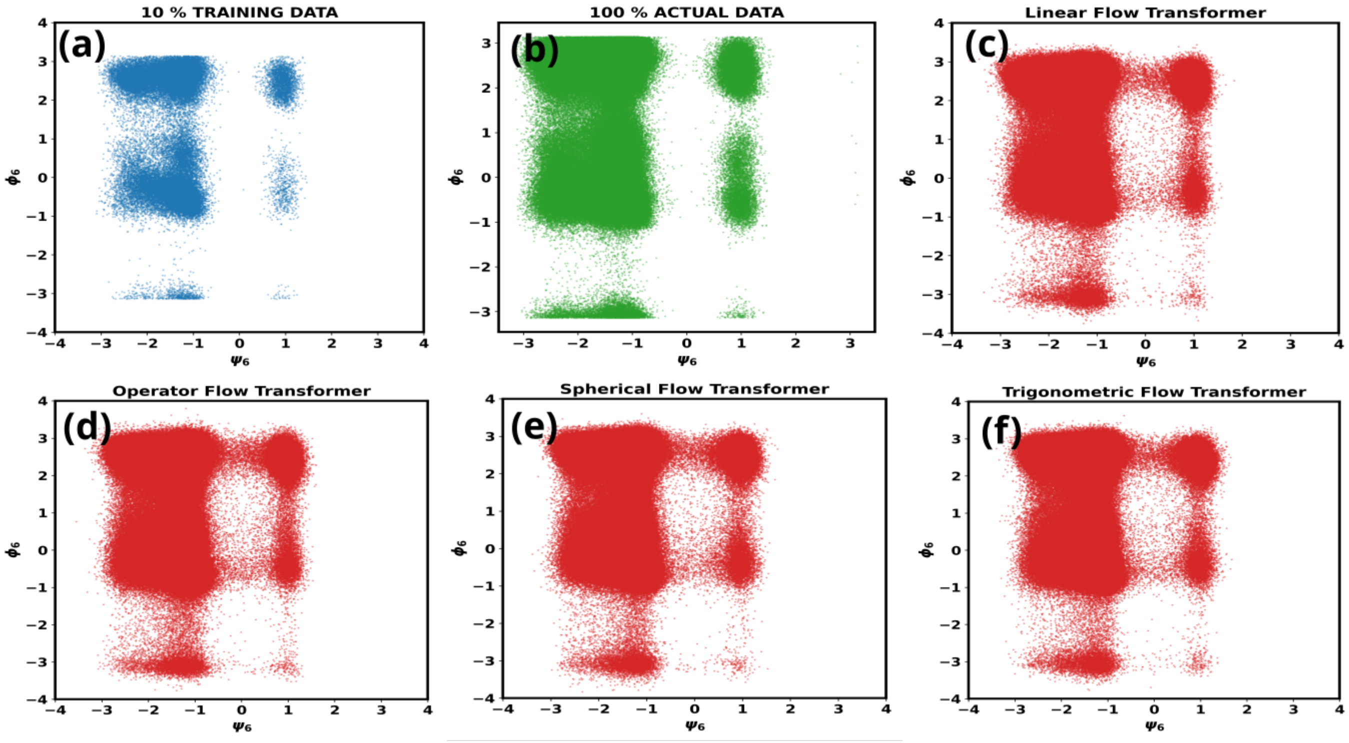


FIG. S4. **Trp-cage** $(\psi_6, \phi_6)$ **marginal distribution.** (a) Training subset; (b) full reference ensemble; (c)–(f) samples generated with the linear, operator-based, trigonometric and spherical paths.

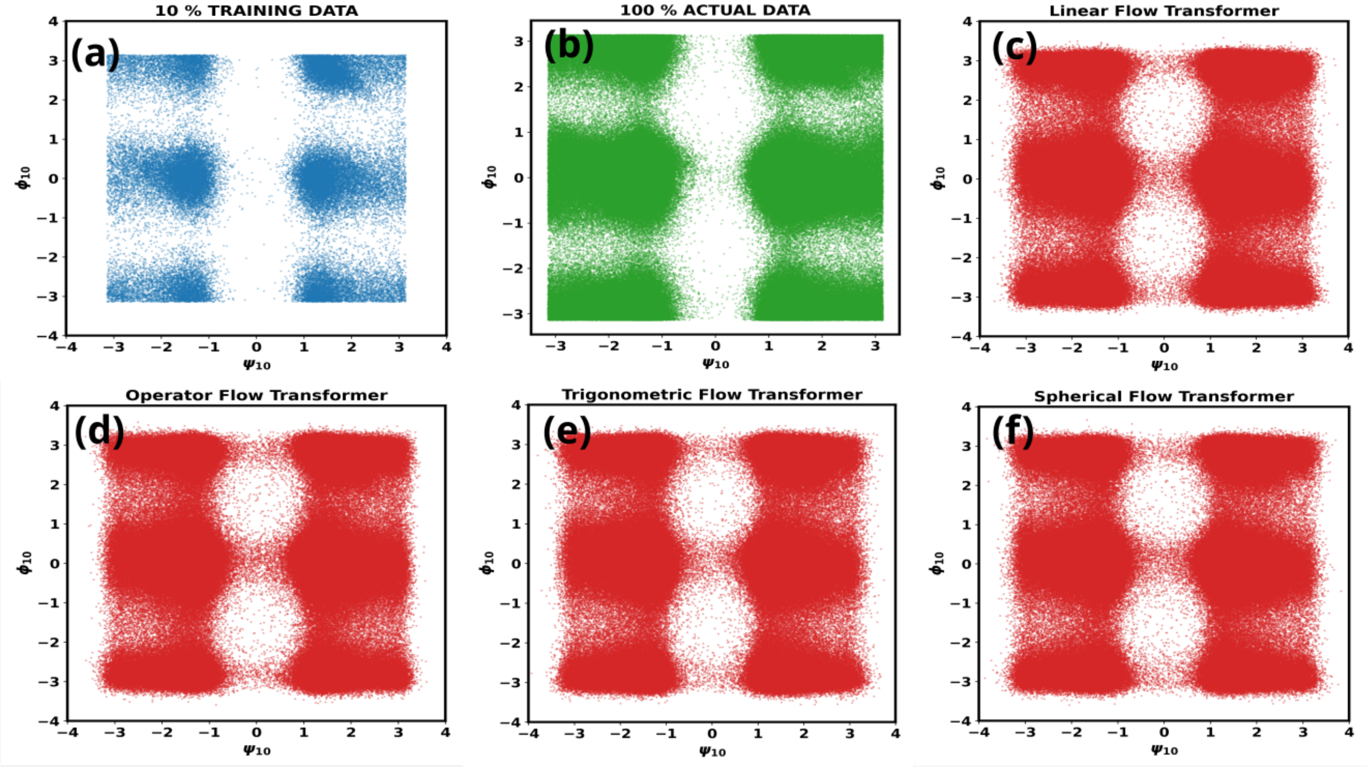


FIG. S5. **Trp-cage $(\psi_{10}, \phi_{10})$ marginal distribution.** Panels as in Fig. S4.

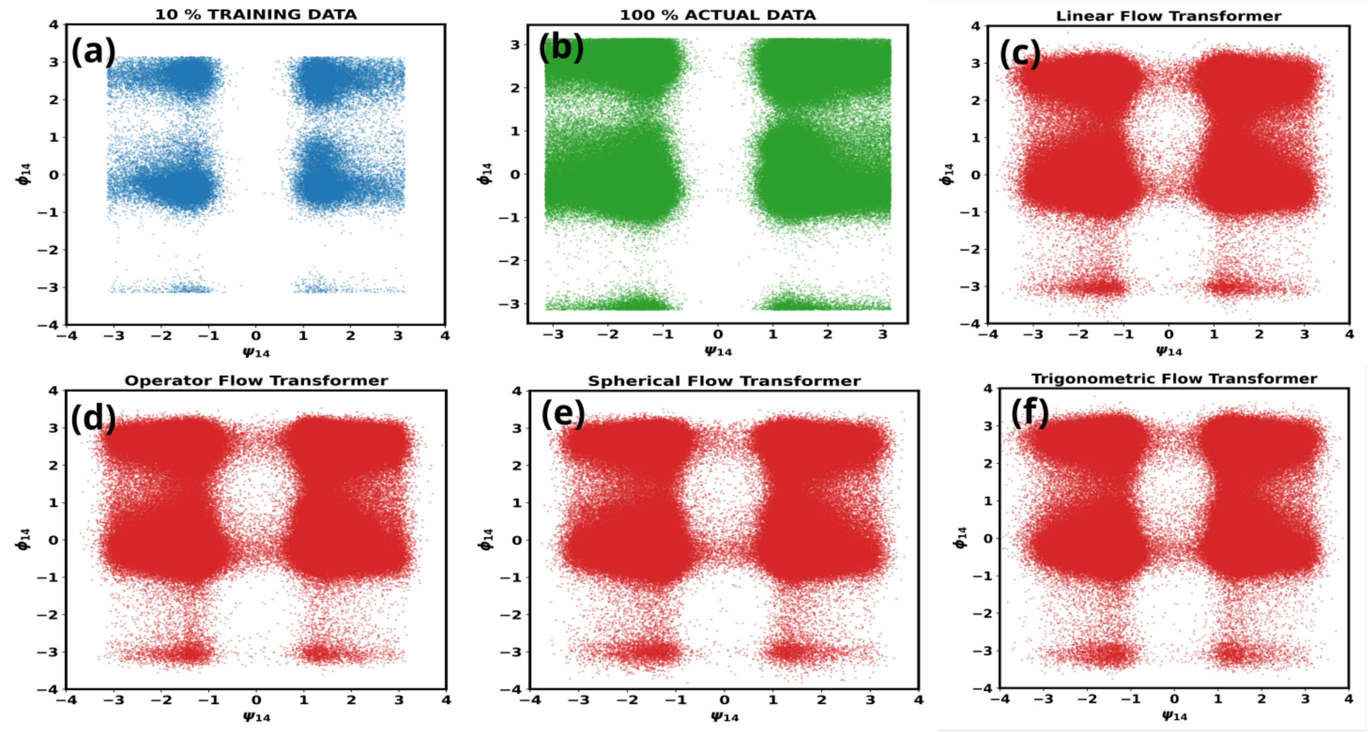


FIG. S6. **Trp-cage $(\psi_{14}, \phi_{14})$ marginal distribution.** Panels as in Fig. S4.

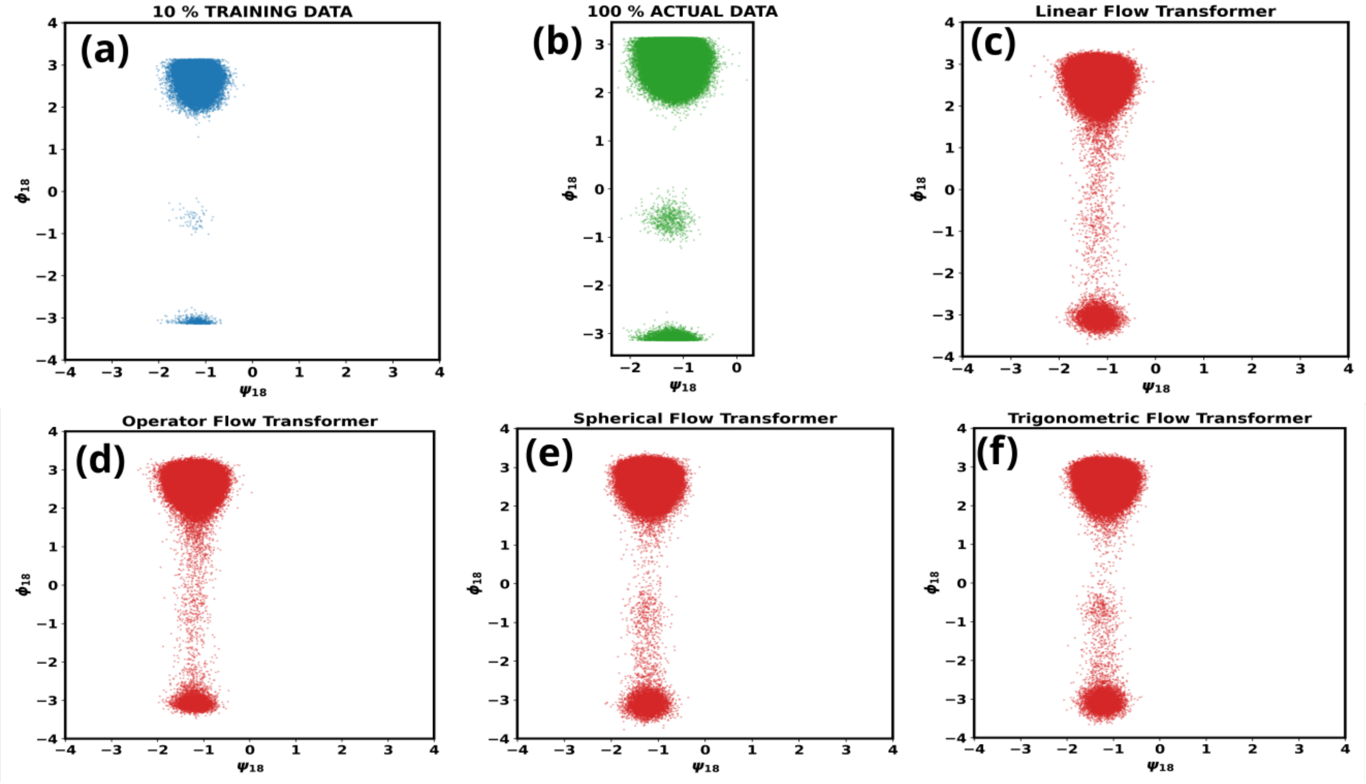


FIG. S7. **Trp-cage ($\psi_{18}, \phi_{18}$) marginal distribution.** Panels as in Fig. S4.

#### B. KL divergence and velocity–score interaction

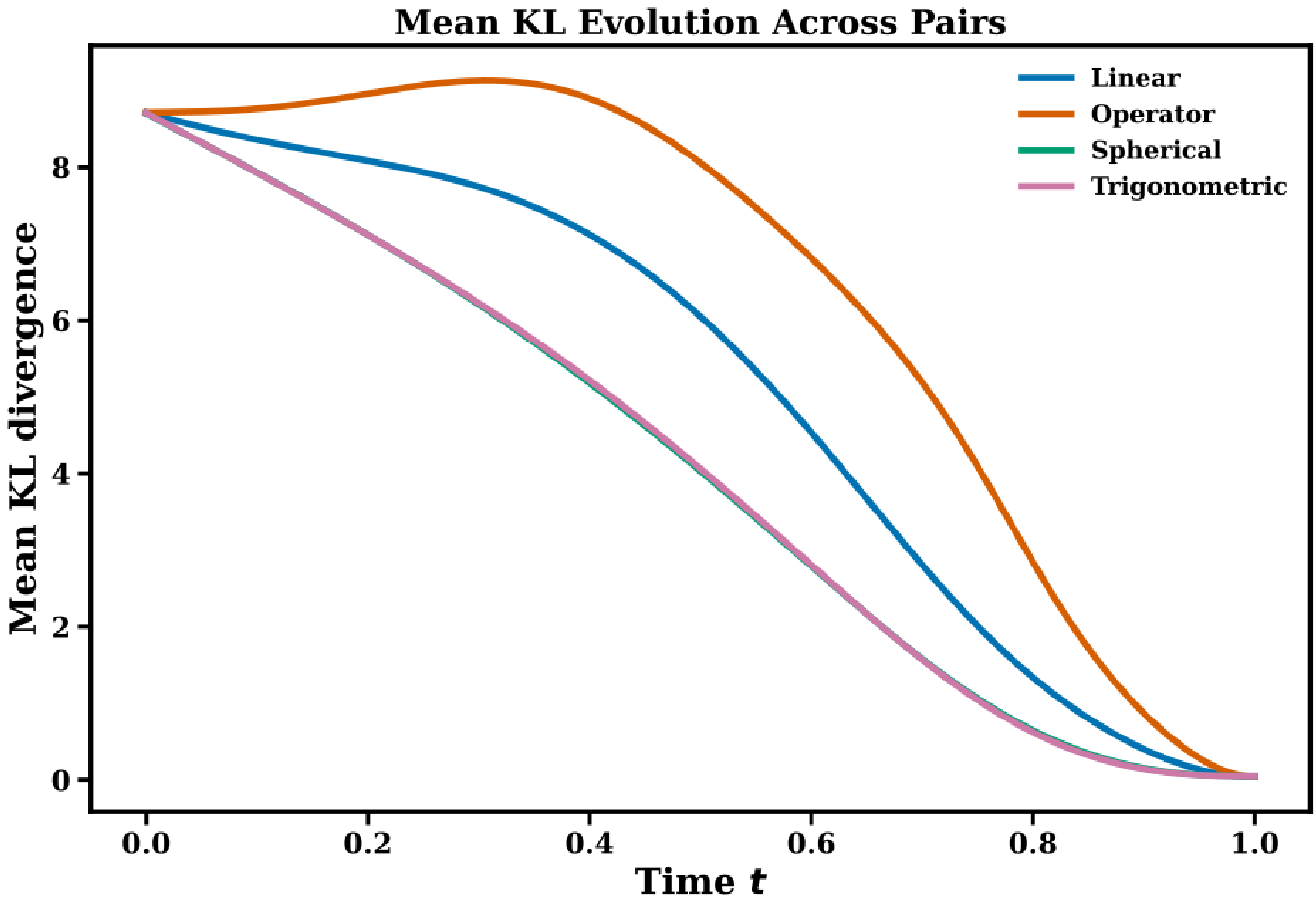


FIG. S8. **Trp-cage: marginal KL divergence averaged over the 19 $(\psi, \phi)$ pairs as a function of flow time.**

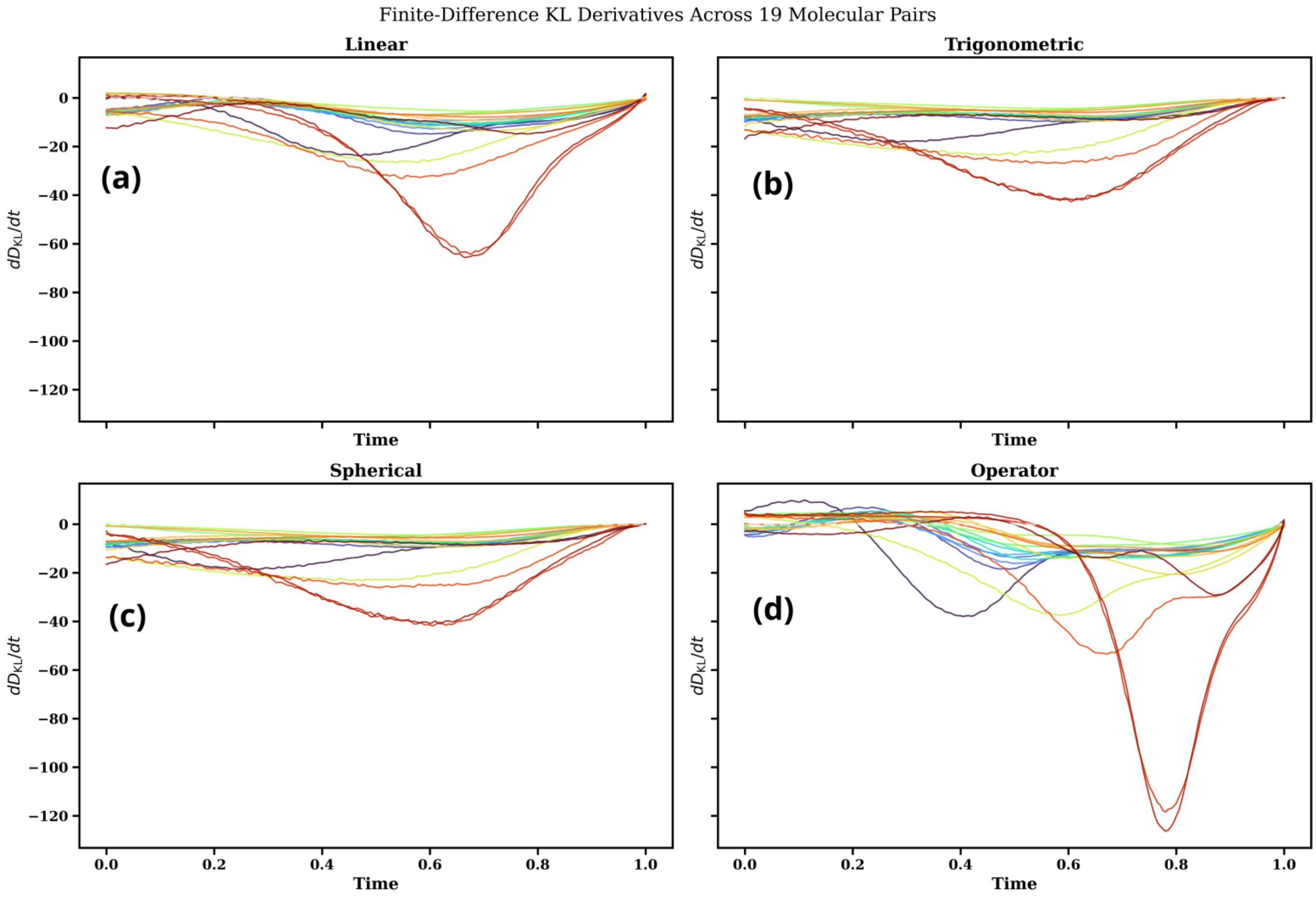


FIG. S9. **Trp-cage: finite-difference estimate of $dD/dt$ for the 19 $(\psi, \phi)$ pairs.** Compare with the velocity–score interaction in Fig. 7 of the main text.

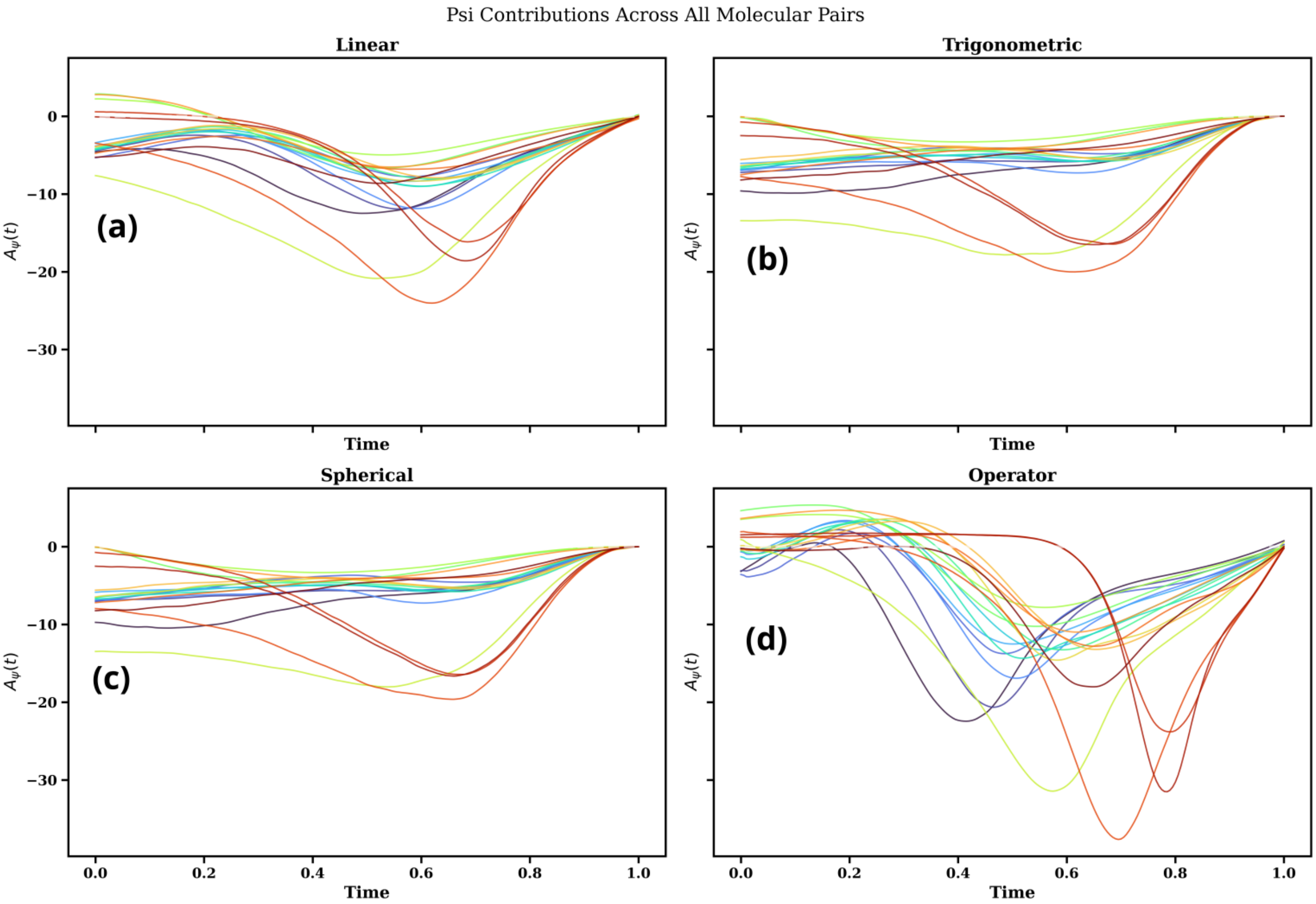


FIG. S10. **Trp-cage: $\psi$ contribution to the velocity–score interaction for the 19 $(\psi, \phi)$ pairs.**

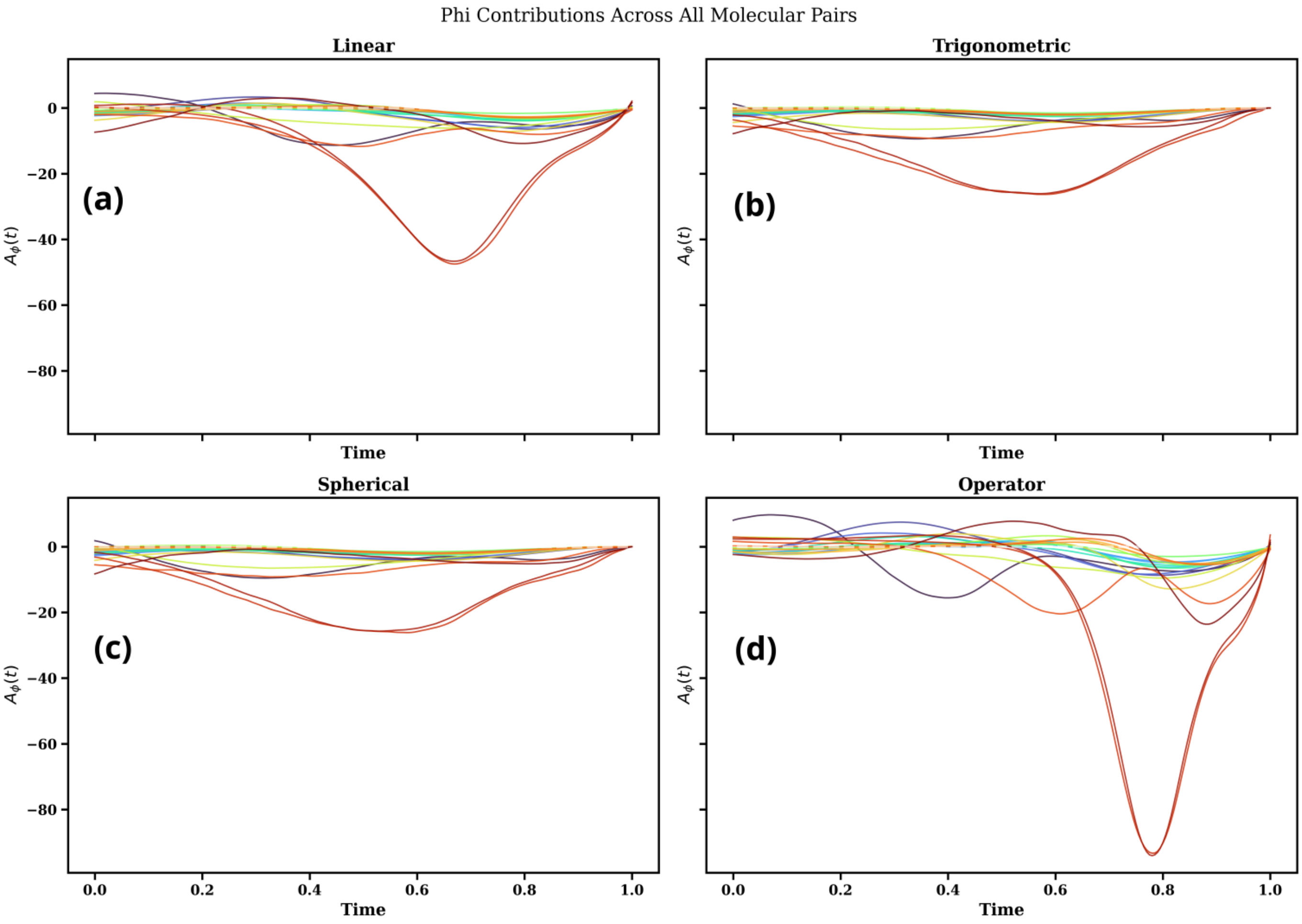


FIG. S11. **Trp-cage: $\phi$ contribution to the velocity–score interaction for the 19 $(\psi, \phi)$ pairs.**

### C. Moment evolution: linear and operator-based paths

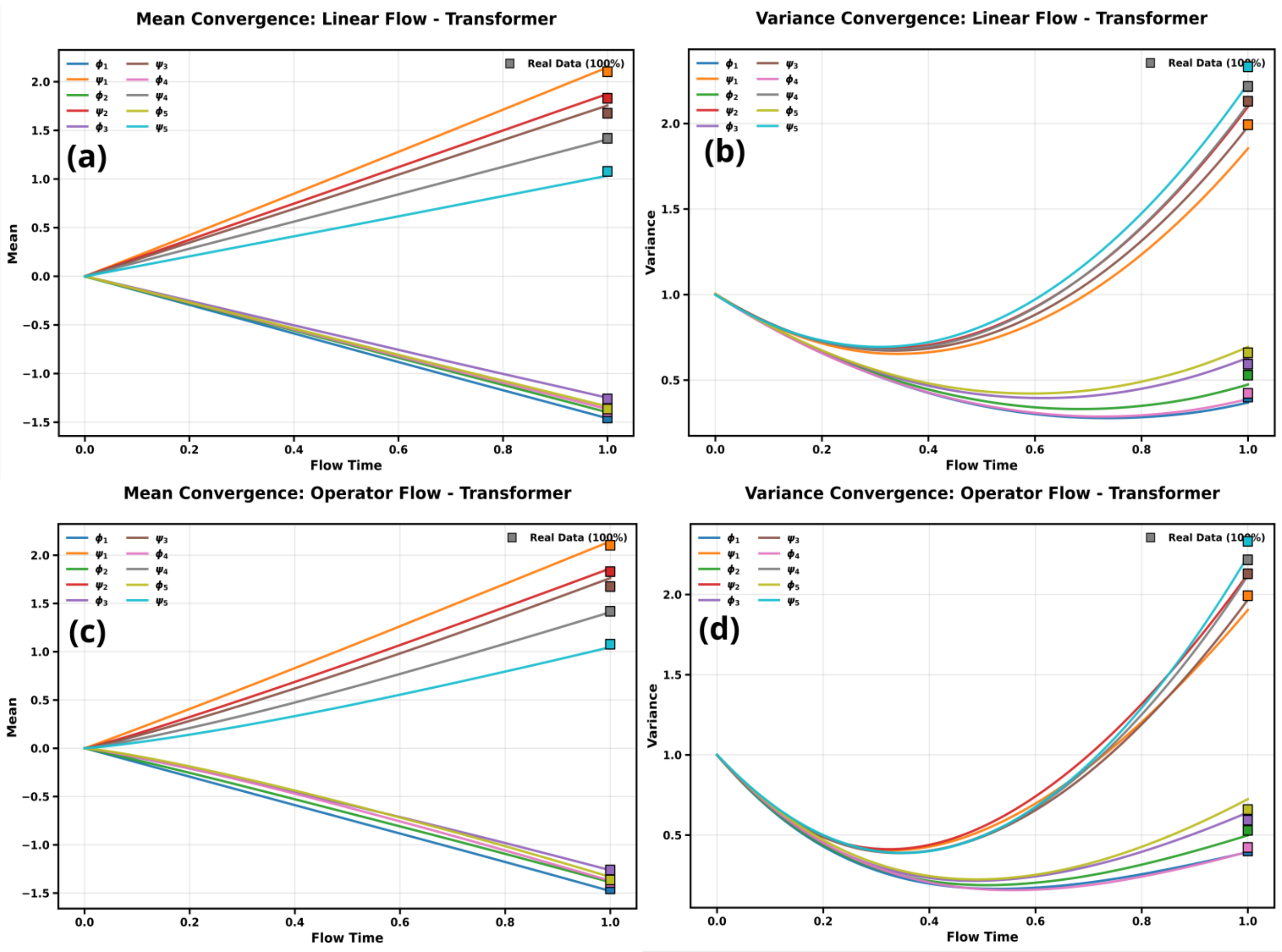


FIG. S12. **Trp-cage, pairs 1–5: mean and variance versus flow time for the linear (a,b) and operator-based (c,d) paths.** Squares mark the reference values.

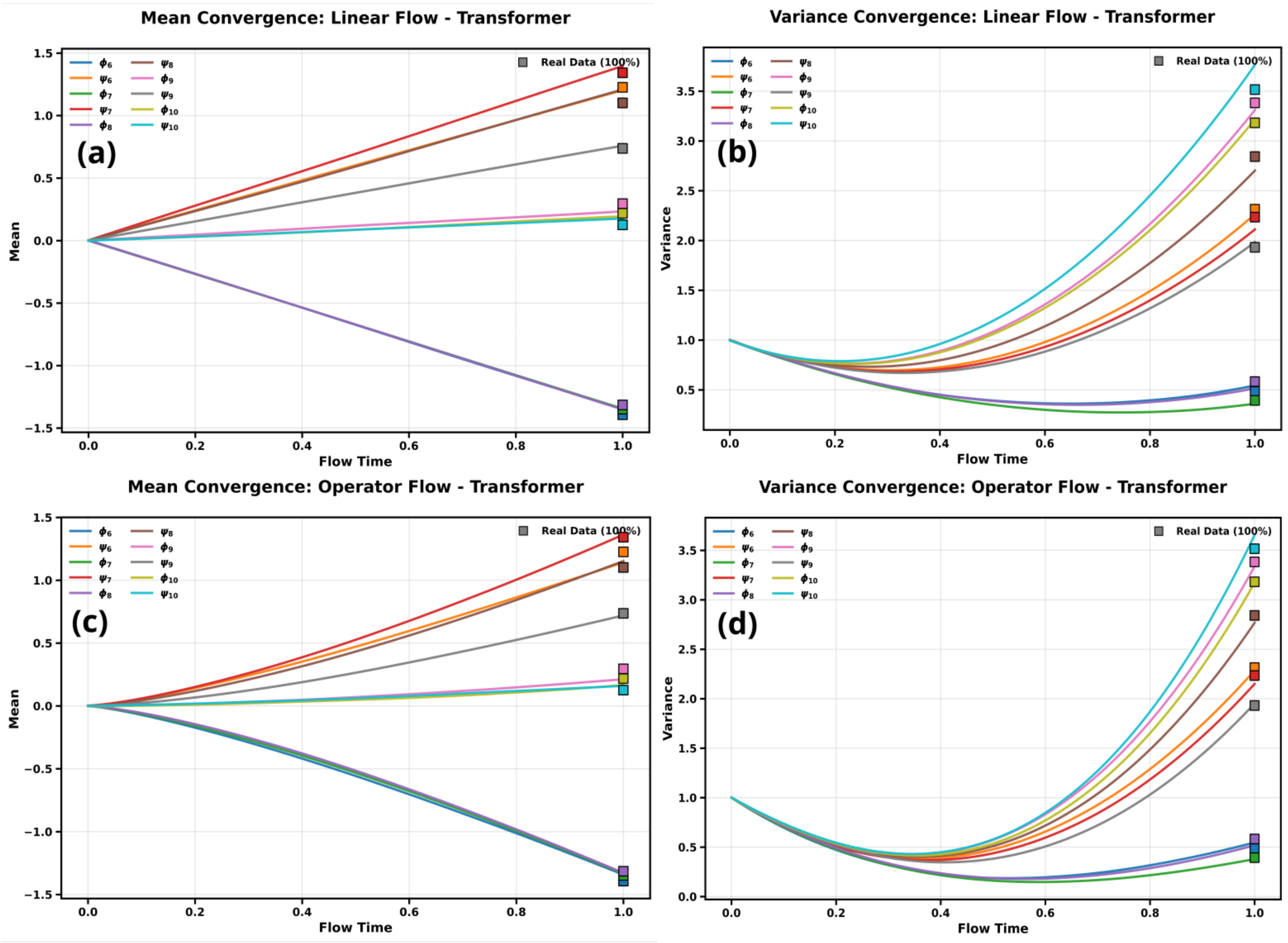


FIG. S13. **Trp-cage, pairs 6–10: mean and variance for the linear and operator-based paths.** Panels as in Fig. S12.

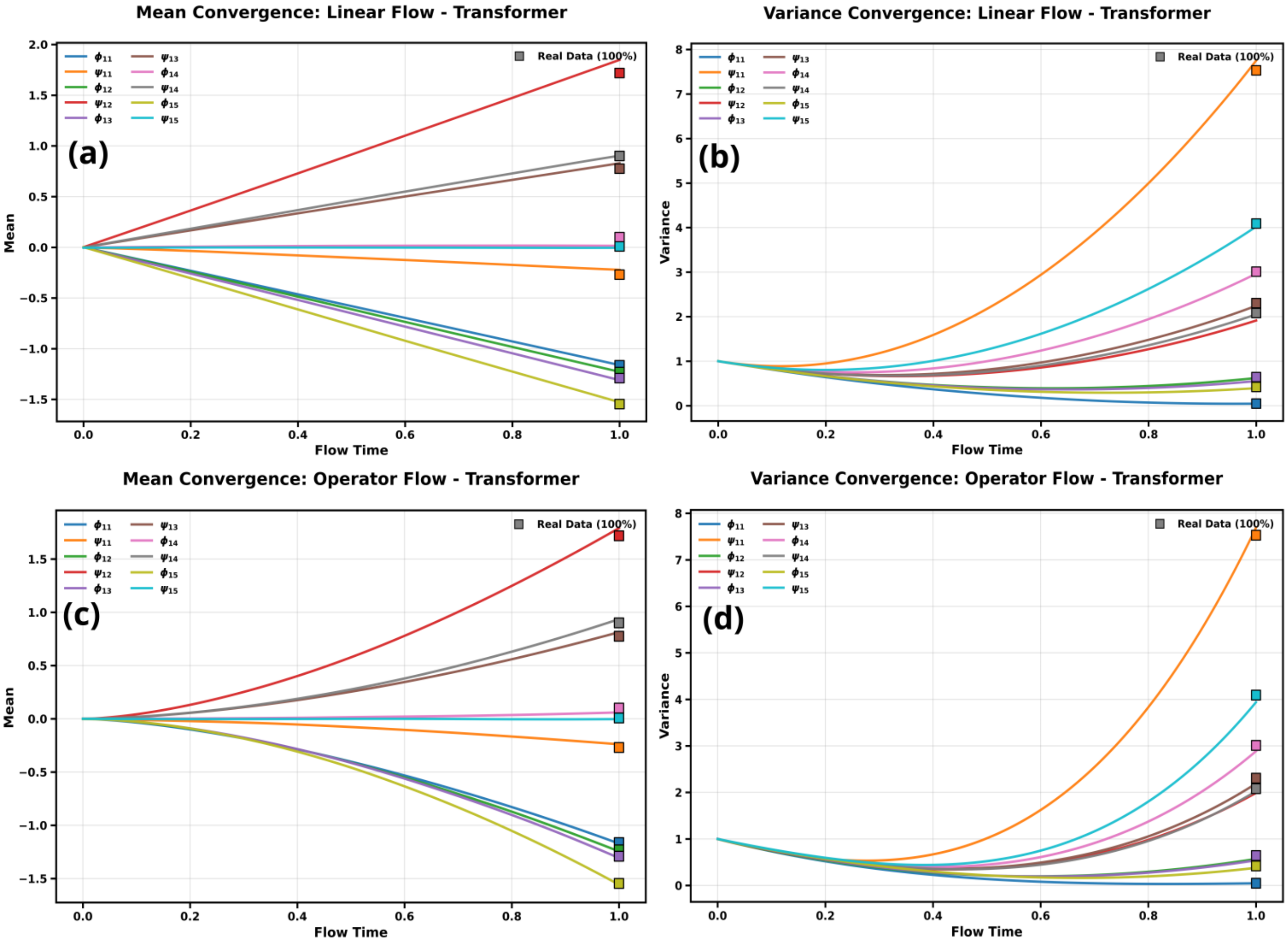


FIG. S14. **Trp-cage, pairs 11–15: mean and variance for the linear and operator-based paths.** Panels as in Fig. S12.

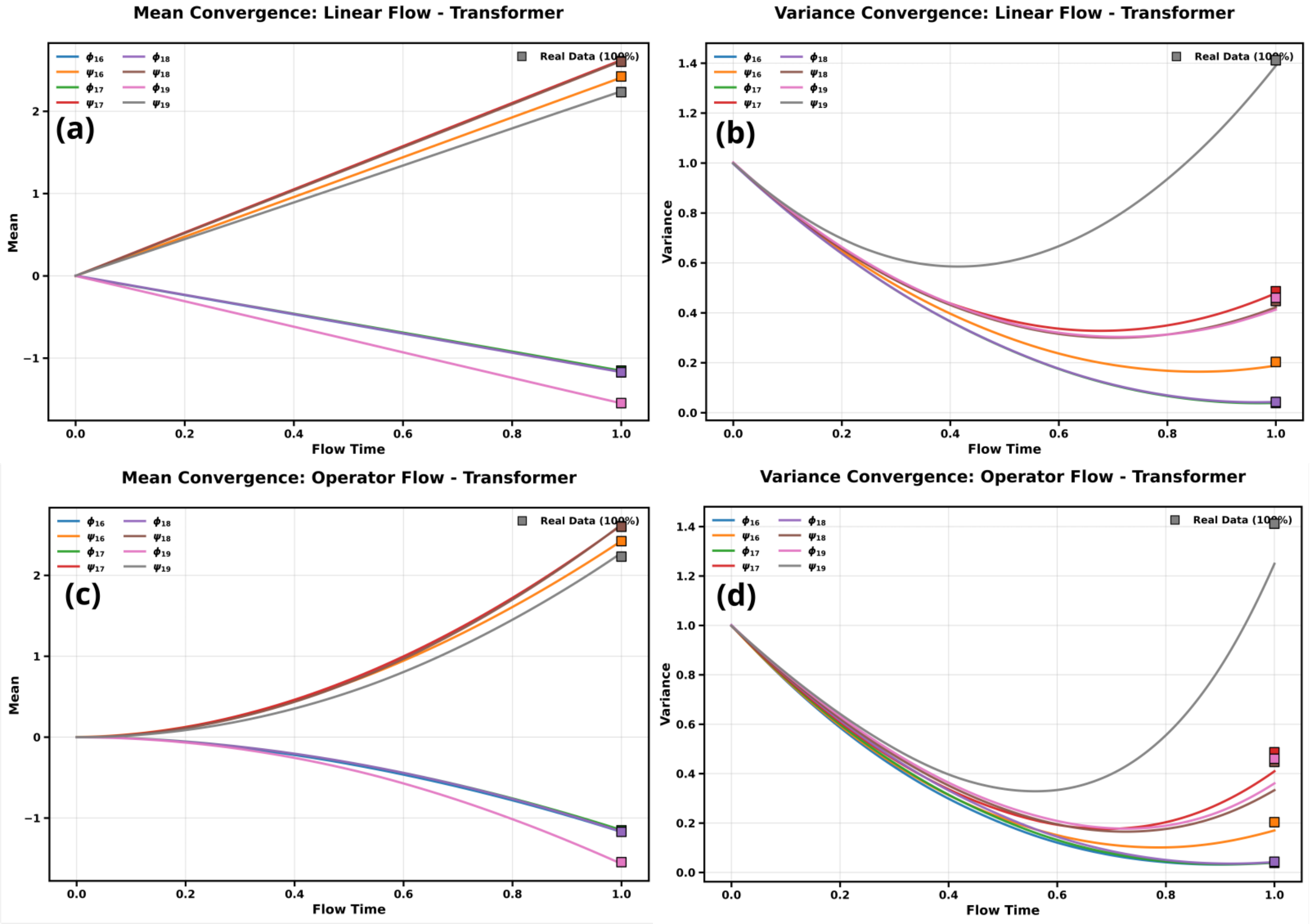


FIG. S15. **Trp-cage, pairs 16–19: mean and variance for the linear and operator-based paths.** Panels as in Fig. S12.

### D. Moment evolution: trigonometric and spherical paths

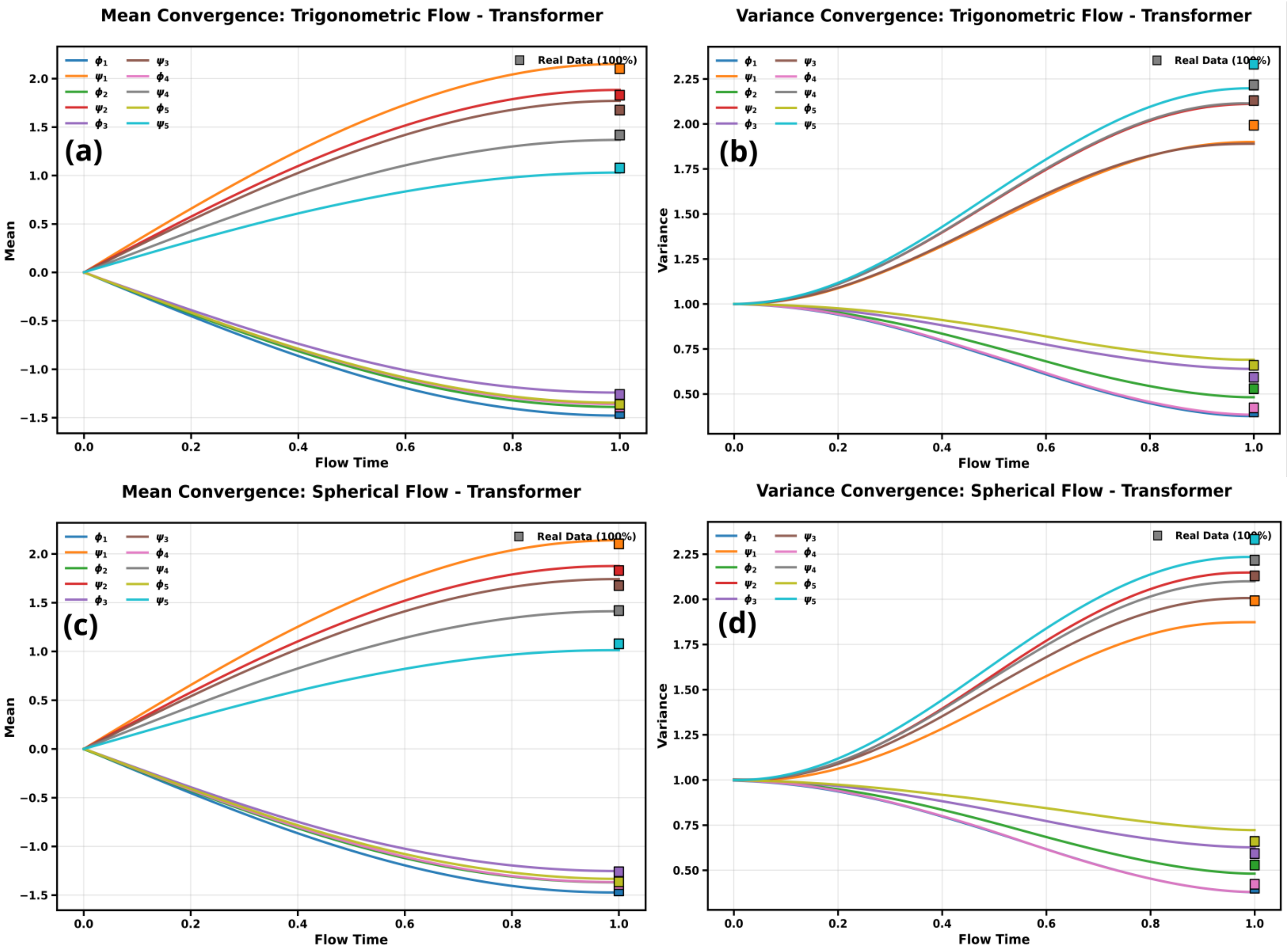


FIG. S16. **Trp-cage, pairs 1–5: mean and variance versus flow time for the trigonometric (a,b) and spherical (c,d) paths.** Squares mark the reference values.

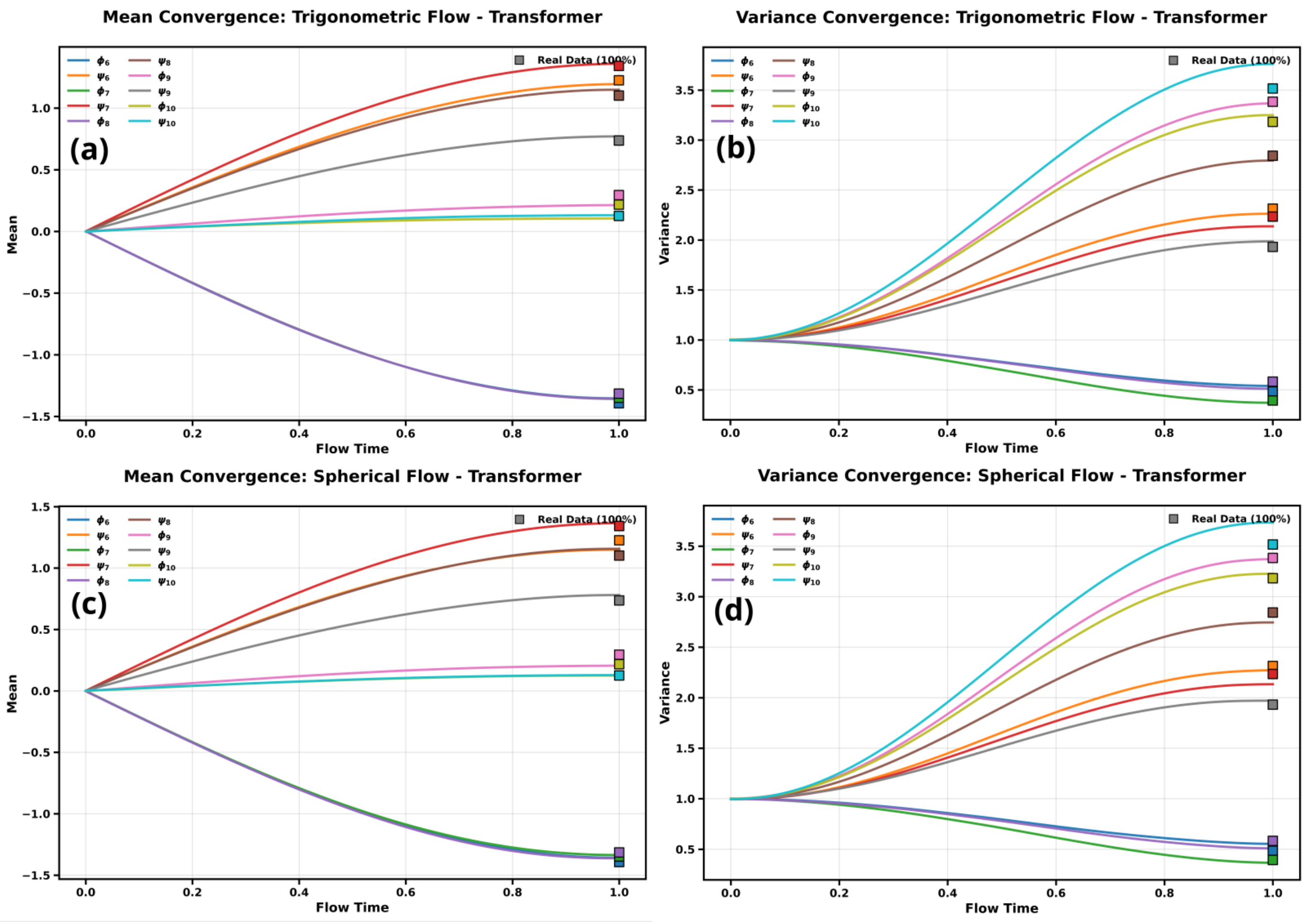


FIG. S17. **Trp-cage, pairs 6–10: mean and variance for the trigonometric and spherical paths.** Panels as in Fig. S16.

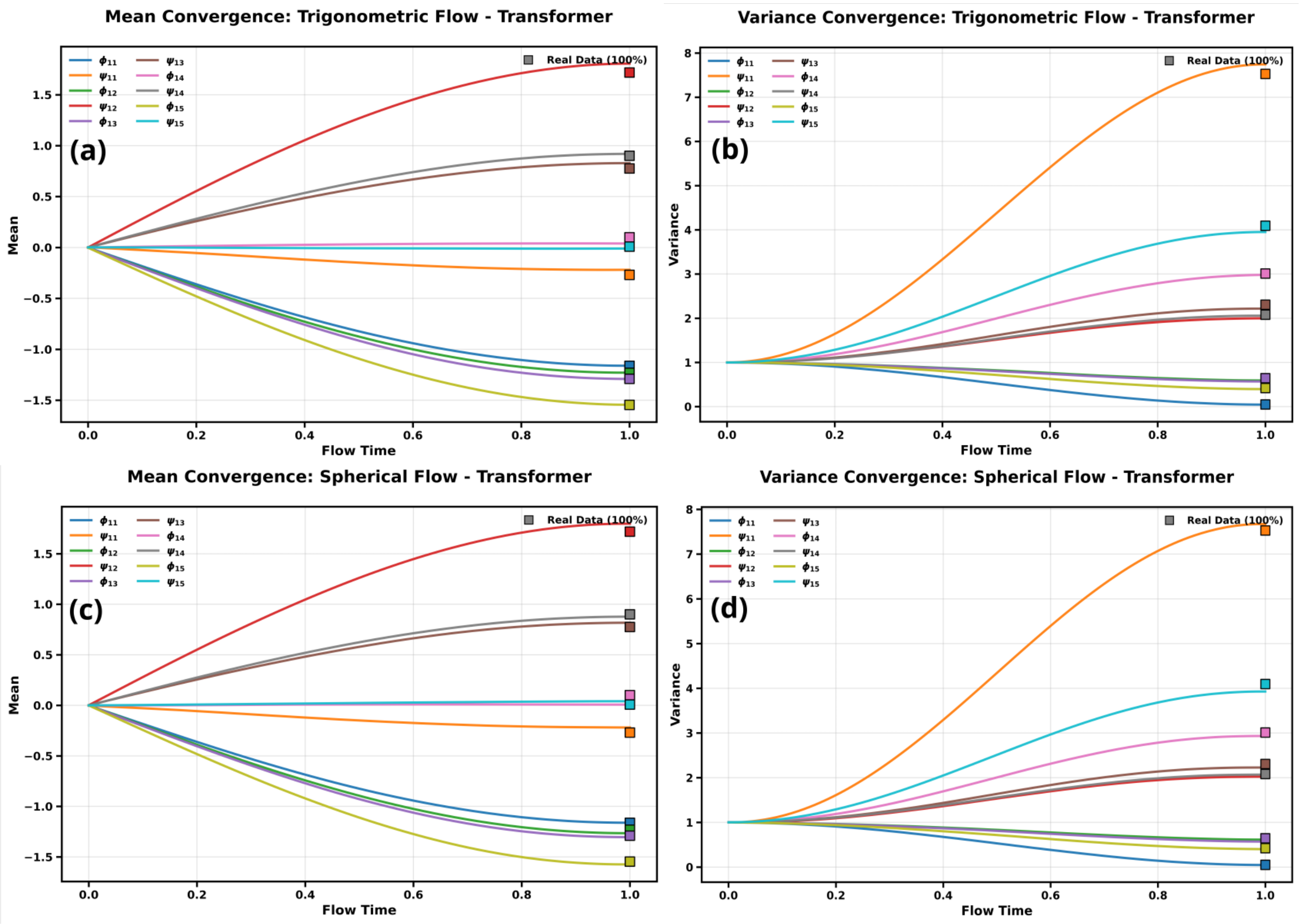


FIG. S18. **Trp-cage, pairs 11–15: mean and variance for the trigonometric and spherical paths.** Panels as in Fig. S16.

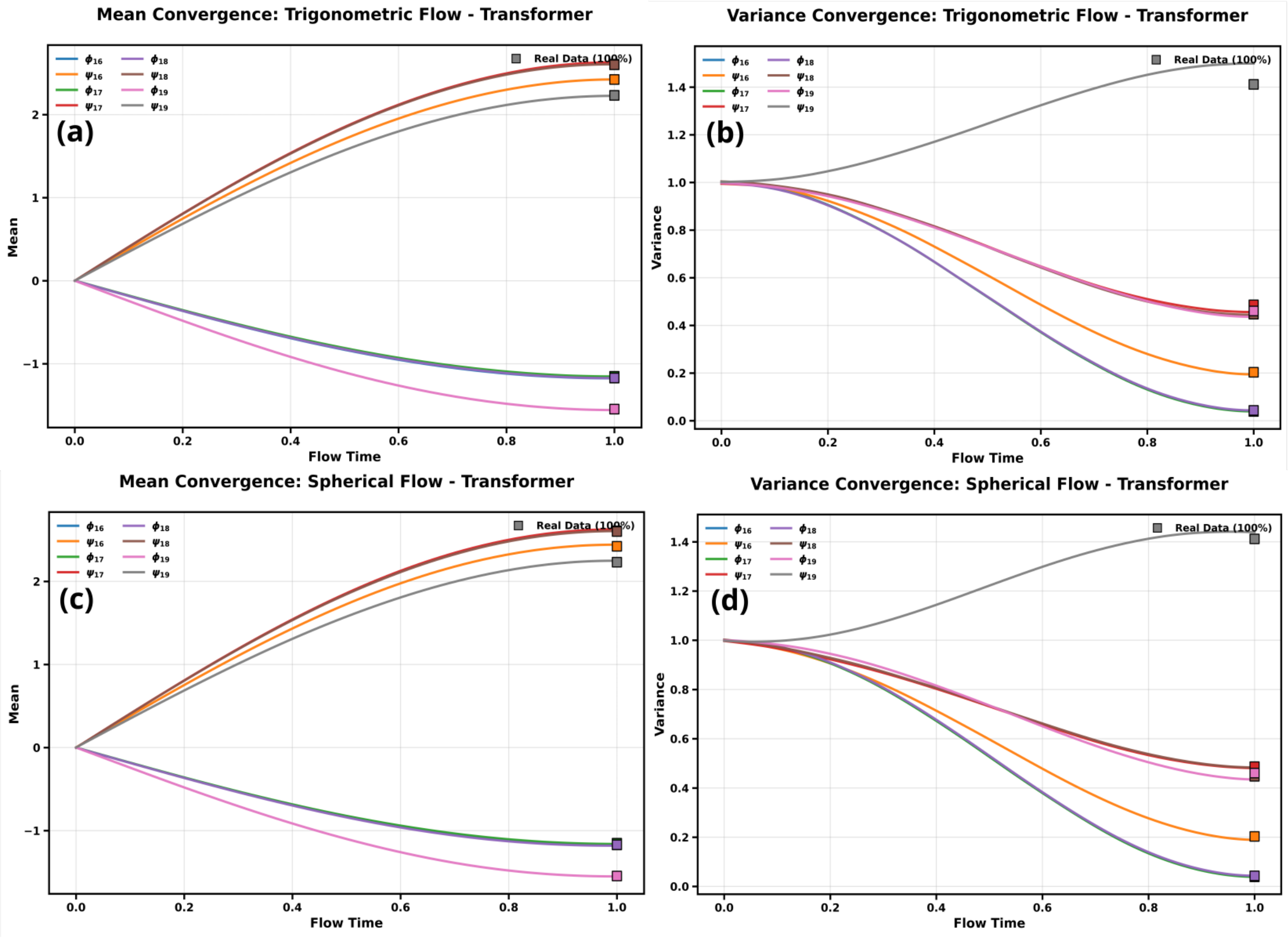


FIG. S19. **Trp-cage, pairs 16–19: mean and variance for the trigonometric and spherical paths.** Panels as in Fig. S16.

### E. Evolution of a marginal density over flow time

We follow the $(\psi_{11}, \phi_{11})$ marginal density along the flow and divide the trajectory into three phases.

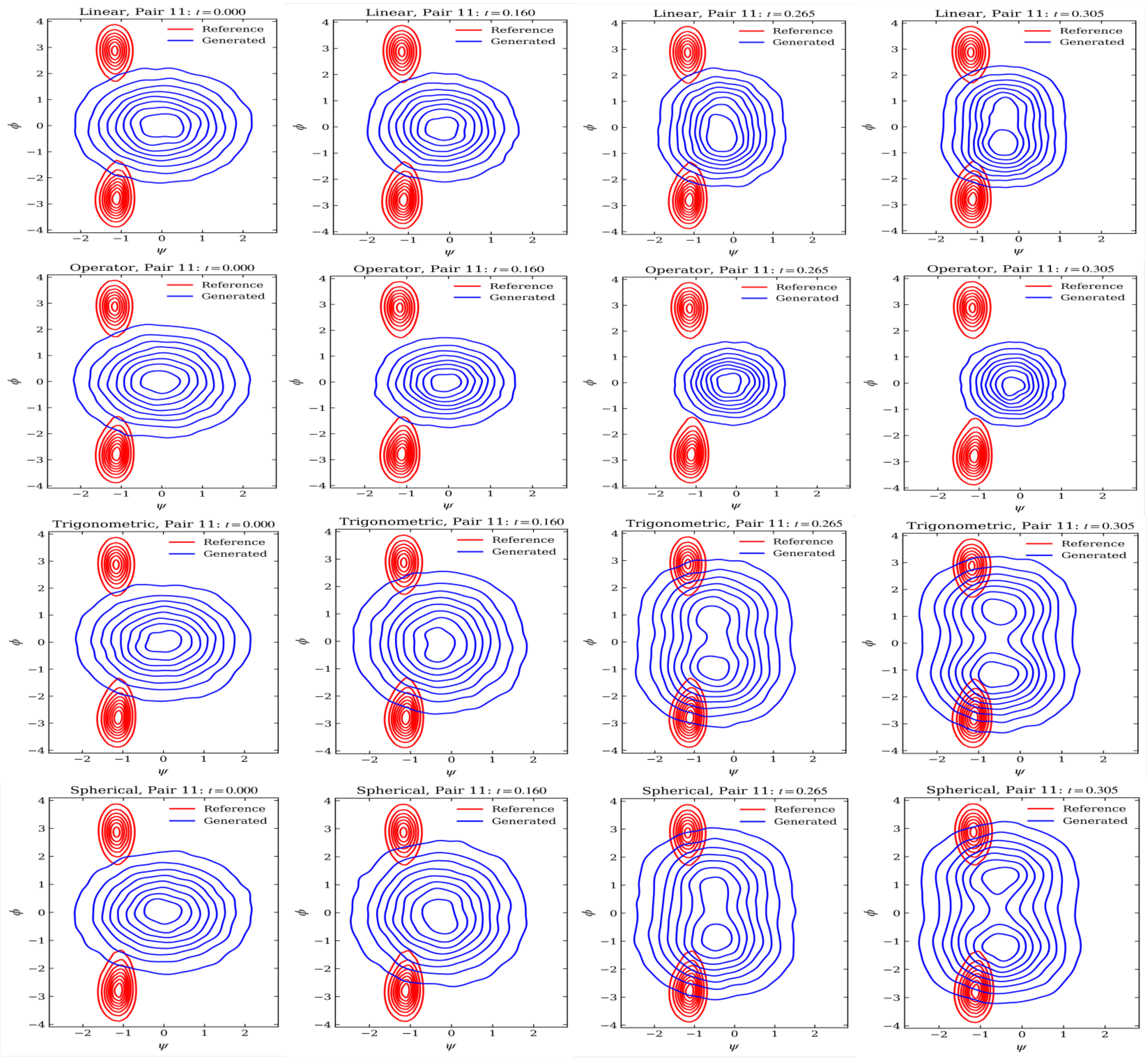


FIG. S20. **Phase I, $t \in [0, 0.31]$: emergence of basins in $(\psi_{11}, \phi_{11})$.** Rows, top to bottom: linear, operator-based, trigonometric and spherical paths (generated in blue, reference in red). The trigonometric and spherical paths already show two distinct basins; the linear path shows the first signs of structure; the operator-based path shows none.

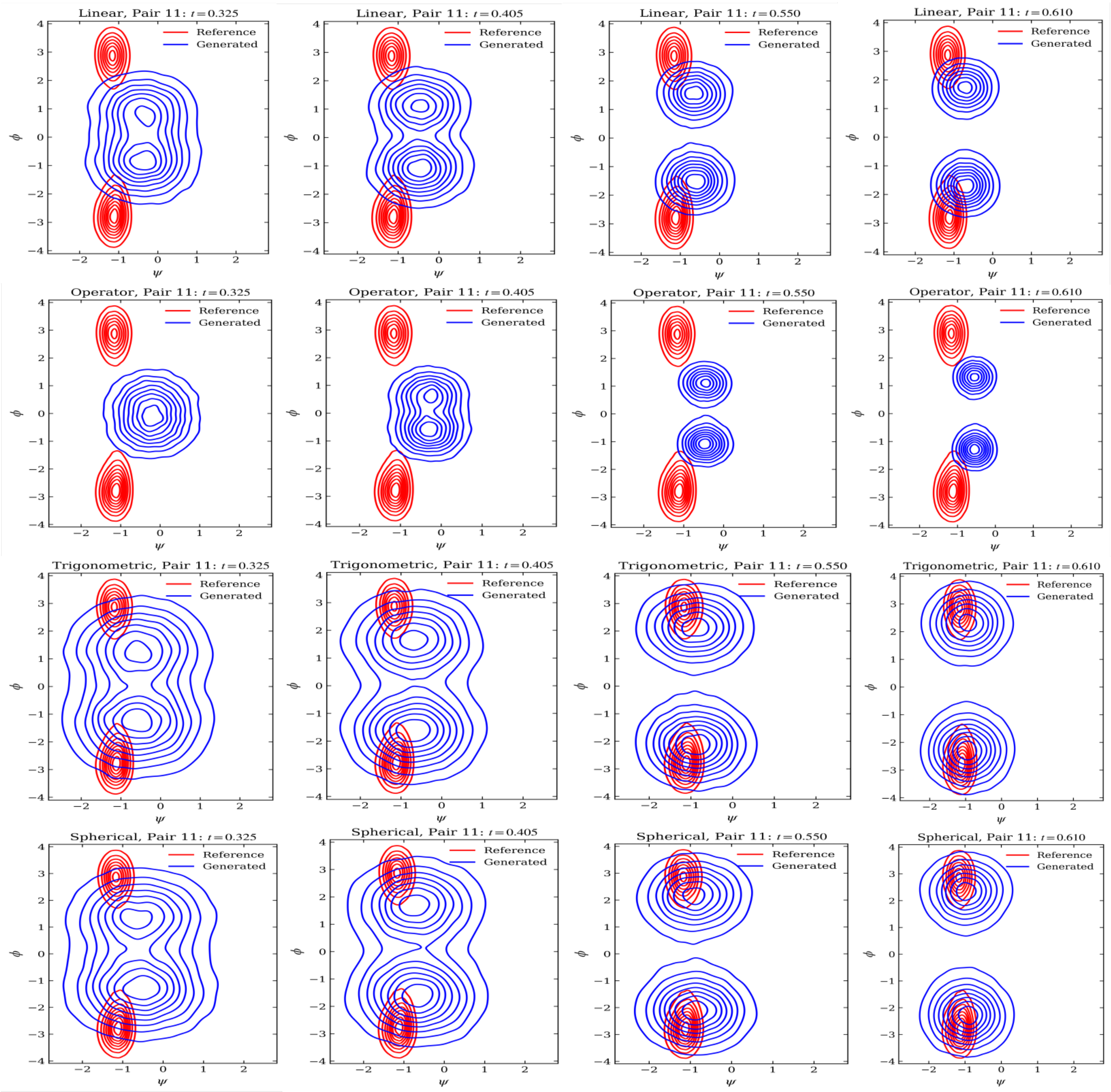


FIG. S21. **Phase II,** $t \in [0.325, 0.61]$**: basin formation.** Rows as in Fig. S20. Basins are visible for all four paths. The spherical and trigonometric paths have essentially recovered the reference basins, the linear path is approaching them, and the operator-based path remains furthest away.

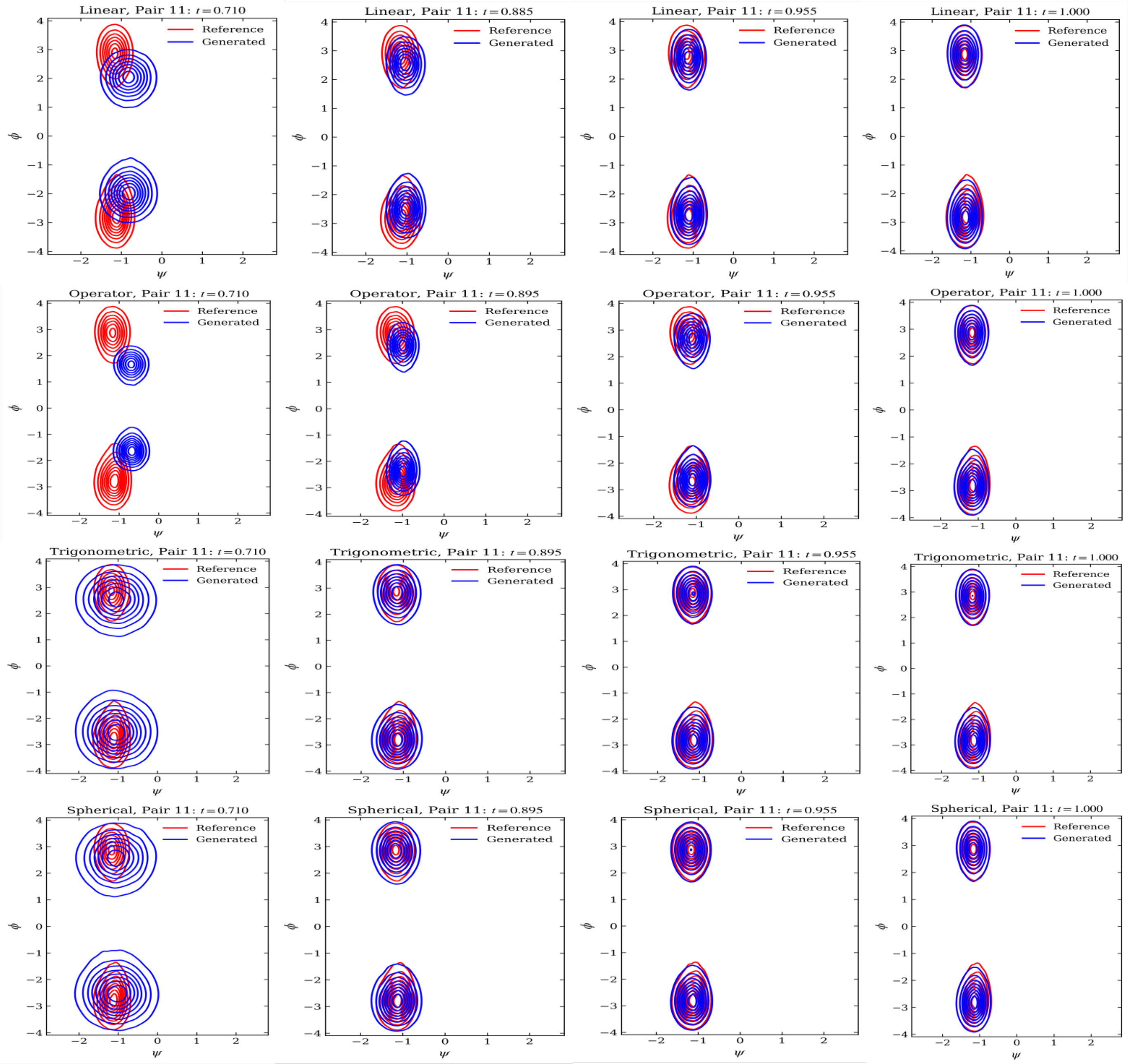


FIG. S22. **Phase III,** $t \in [0.71, 1]$**: completion of convergence.** Rows as in Fig. S20. All four paths reach the reference basins; the linear and operator-based paths do so last.

### S3. $\alpha$-SYNUCLEIN

#### A. Marginal distributions

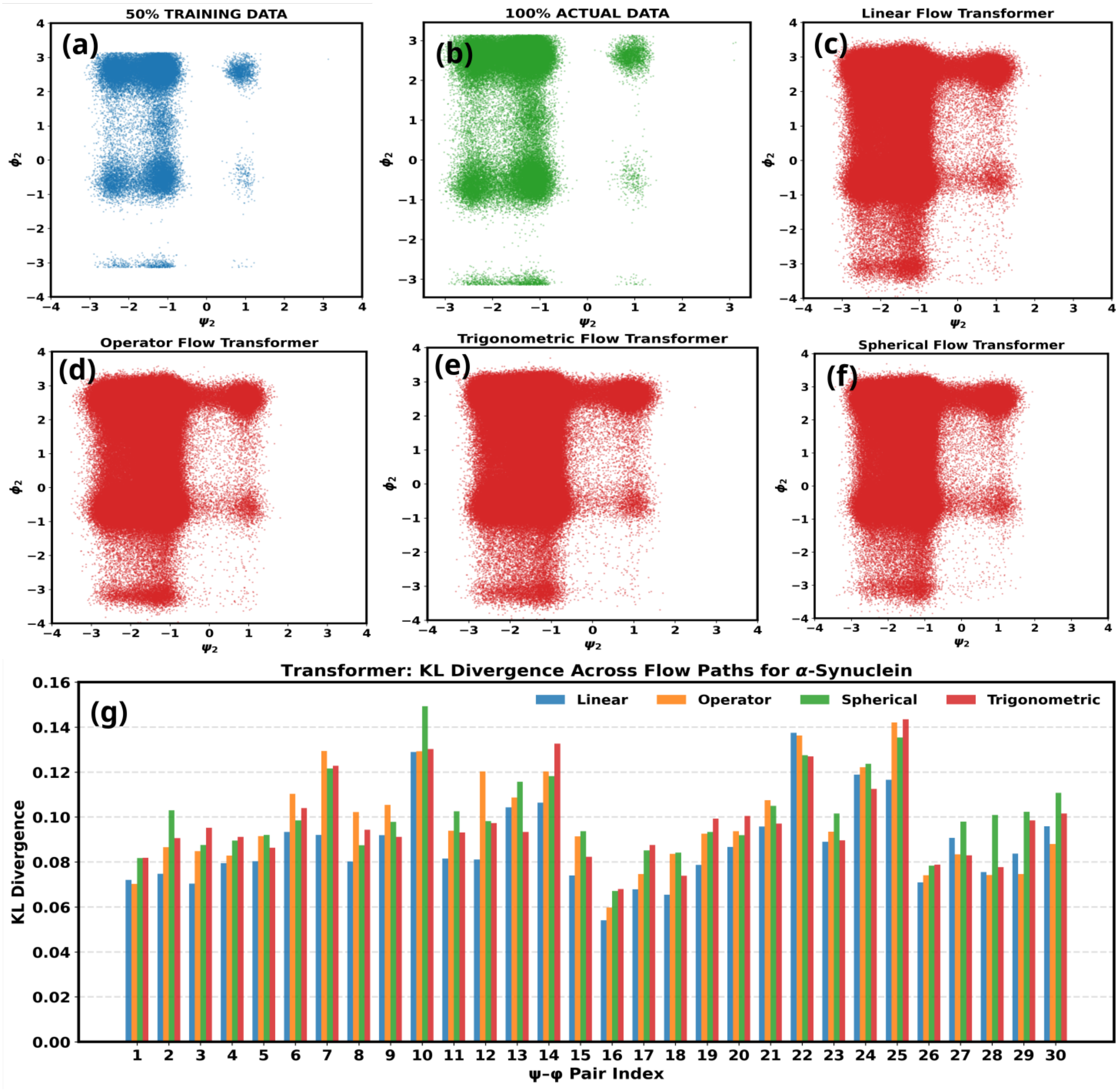


FIG. S23. **$\alpha$-Synuclein $(\psi_2, \phi_2)$ marginal distribution and marginal KL divergences.** (a) Training subset; (b) full reference ensemble; (c)–(f) samples generated with the linear, operator-based, trigonometric and spherical paths. (g) KL divergence between generated and reference marginals for all 30 $(\psi, \phi)$ pairs; all values lie below 0.16.

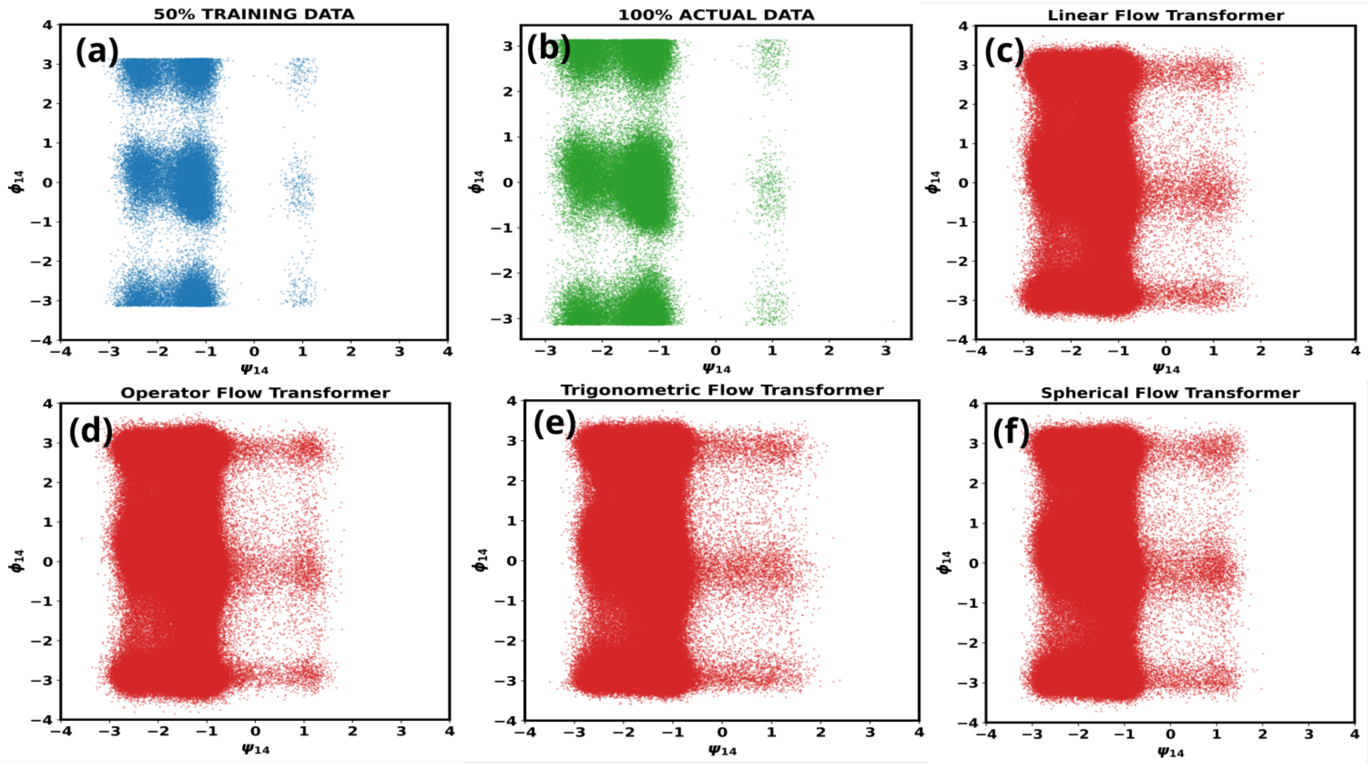


FIG. S24. $\alpha$**-Synuclein** $(\psi_{14}, \phi_{14})$ **marginal distribution.** Panels (a)–(f) as in Fig. S23.

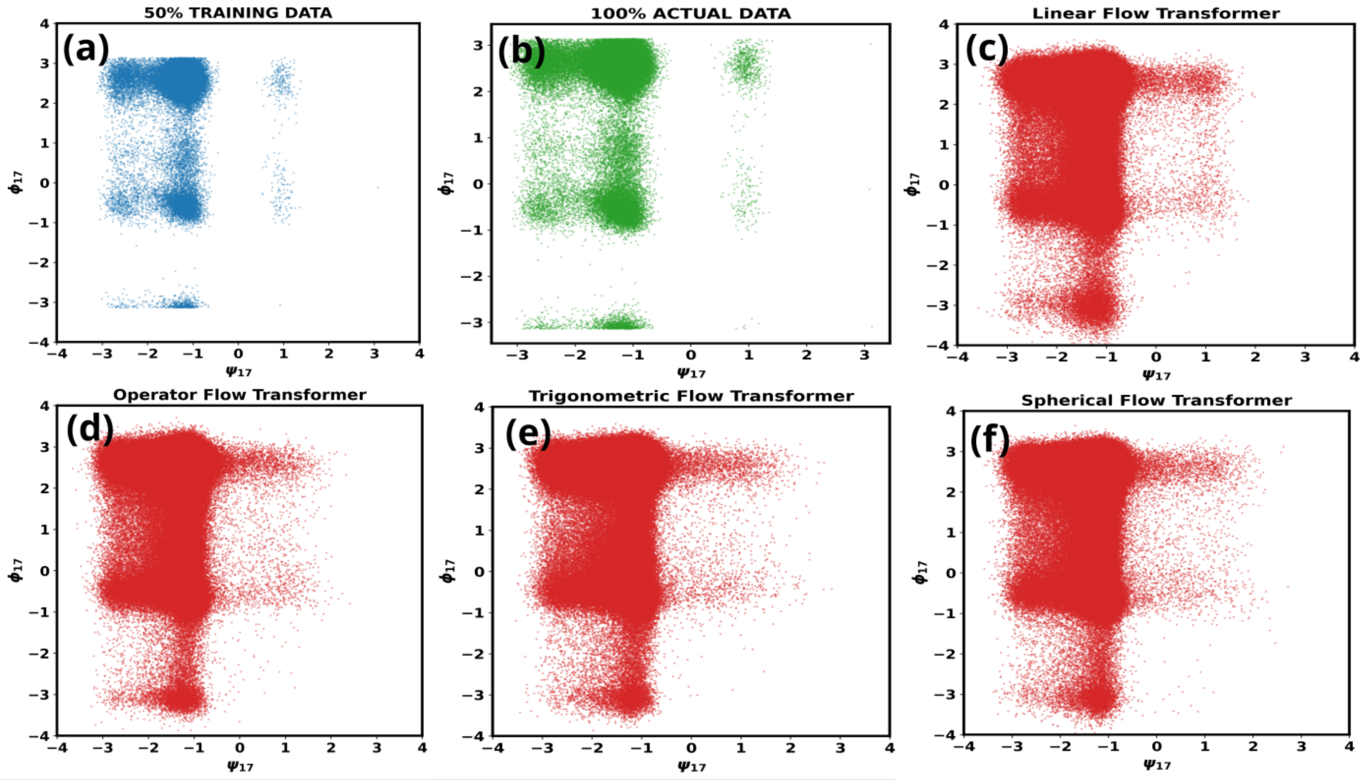


FIG. S25. $\alpha$**-Synuclein** $(\psi_{17}, \phi_{17})$ **marginal distribution.** Panels (a)–(f) as in Fig. S23.

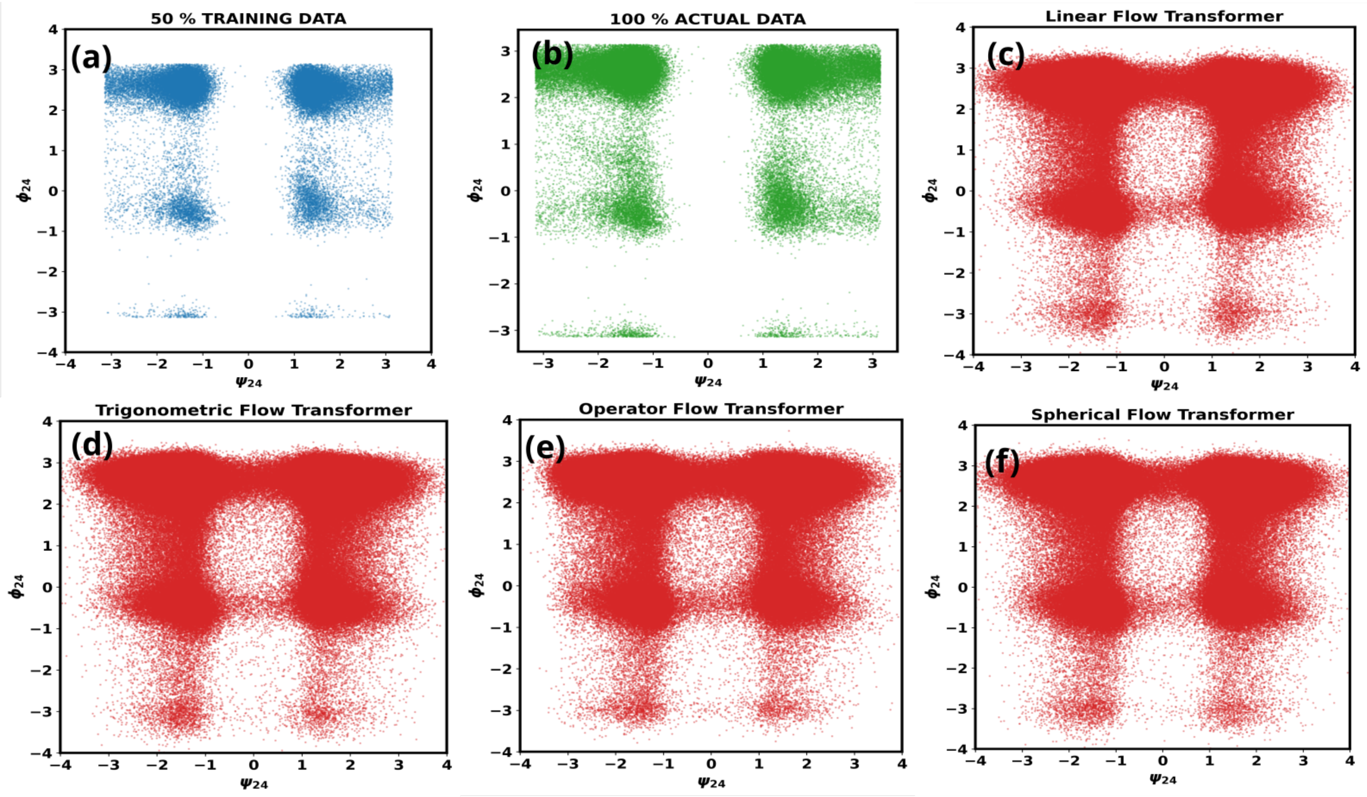


FIG. S26. $\alpha$**-Synuclein** $(\psi_{24}, \phi_{24})$ **marginal distribution.** Panels (a)–(f) as in Fig. S23.

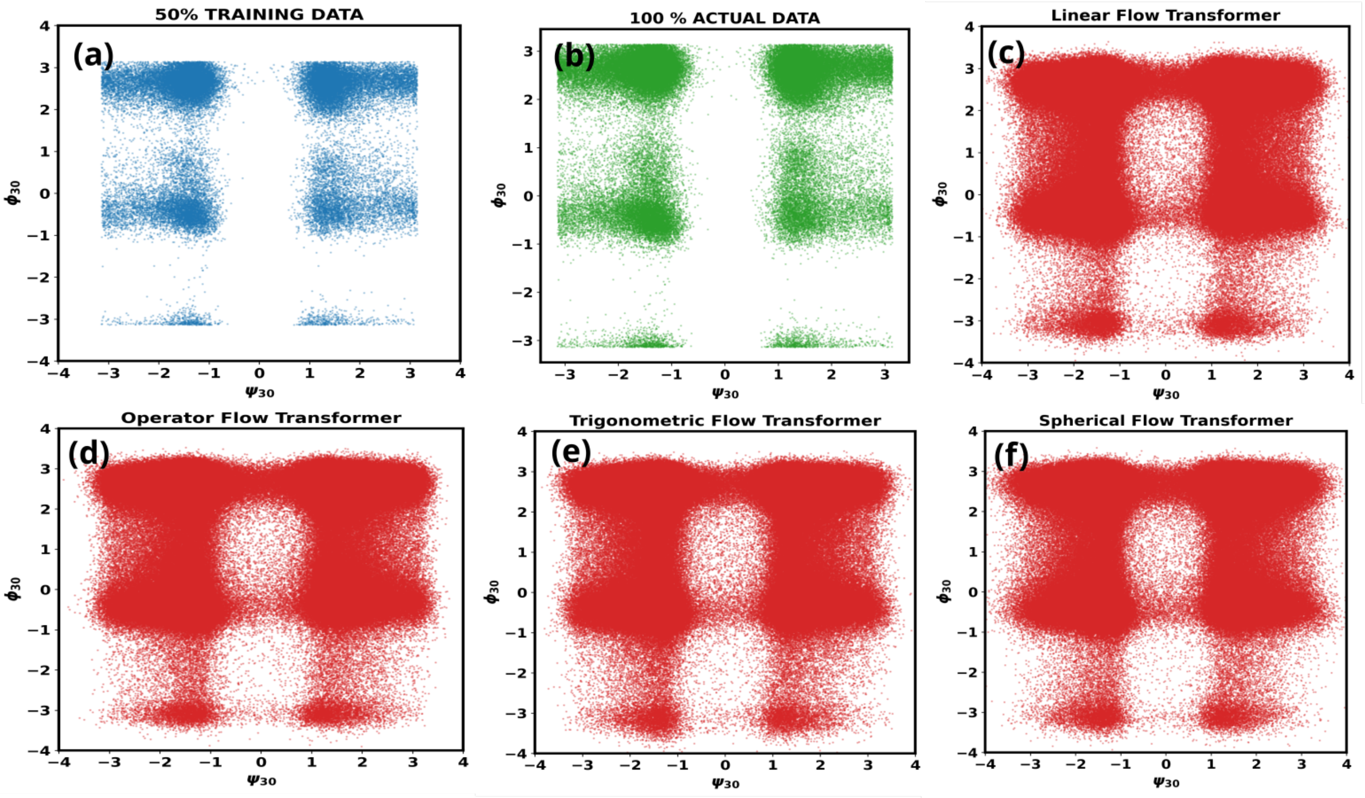


FIG. S27. $\alpha$**-Synuclein** $(\psi_{30}, \phi_{30})$ **marginal distribution.** Panels (a)–(f) as in Fig. S23.

### B. KL divergence and velocity–score interaction

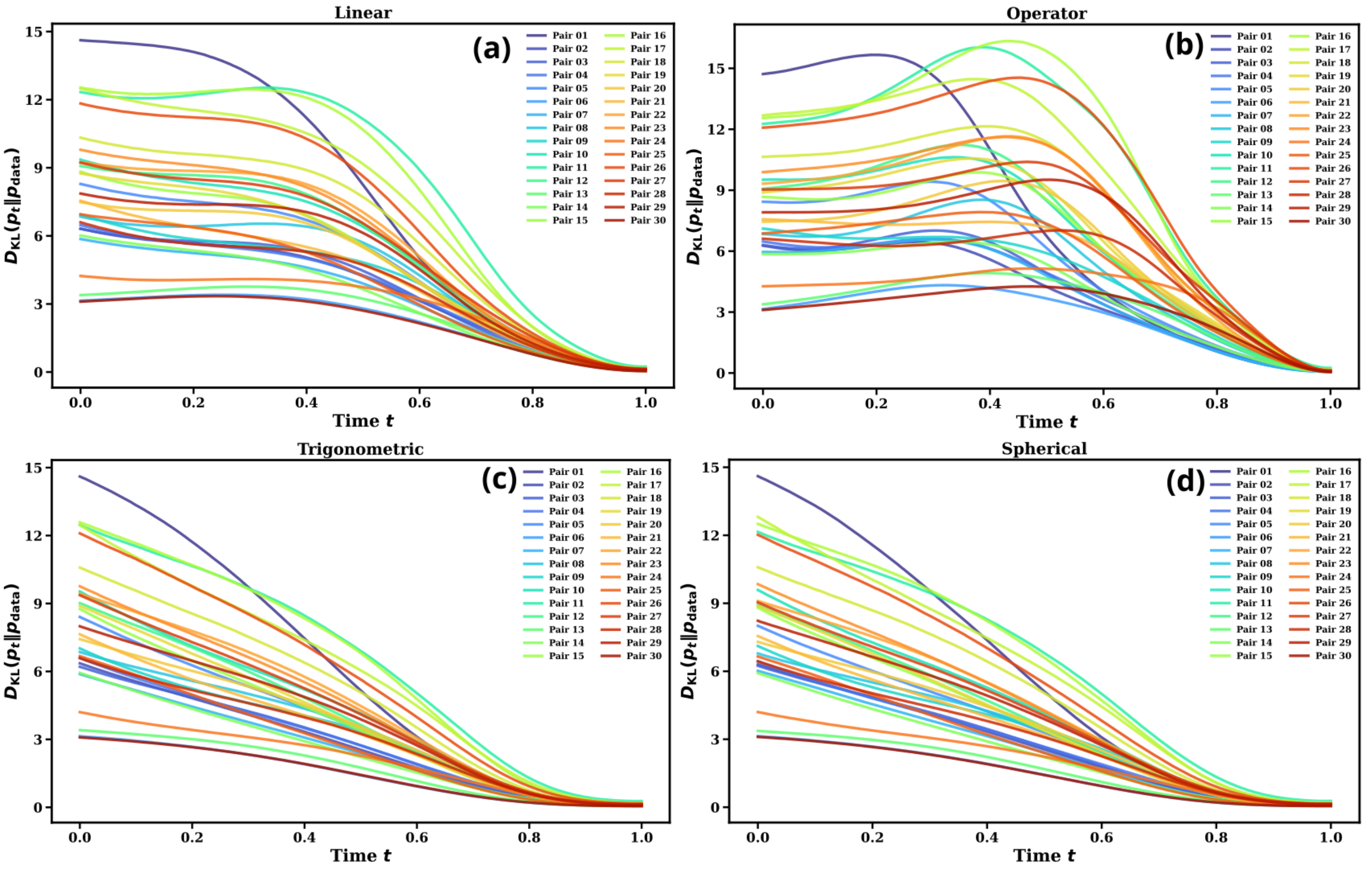


FIG. S28. $\alpha$-**Synuclein: marginal KL divergence versus flow time for the 30** $(\psi, \phi)$ **pairs.** (a)–(d) Linear, operator-based, trigonometric and spherical paths.

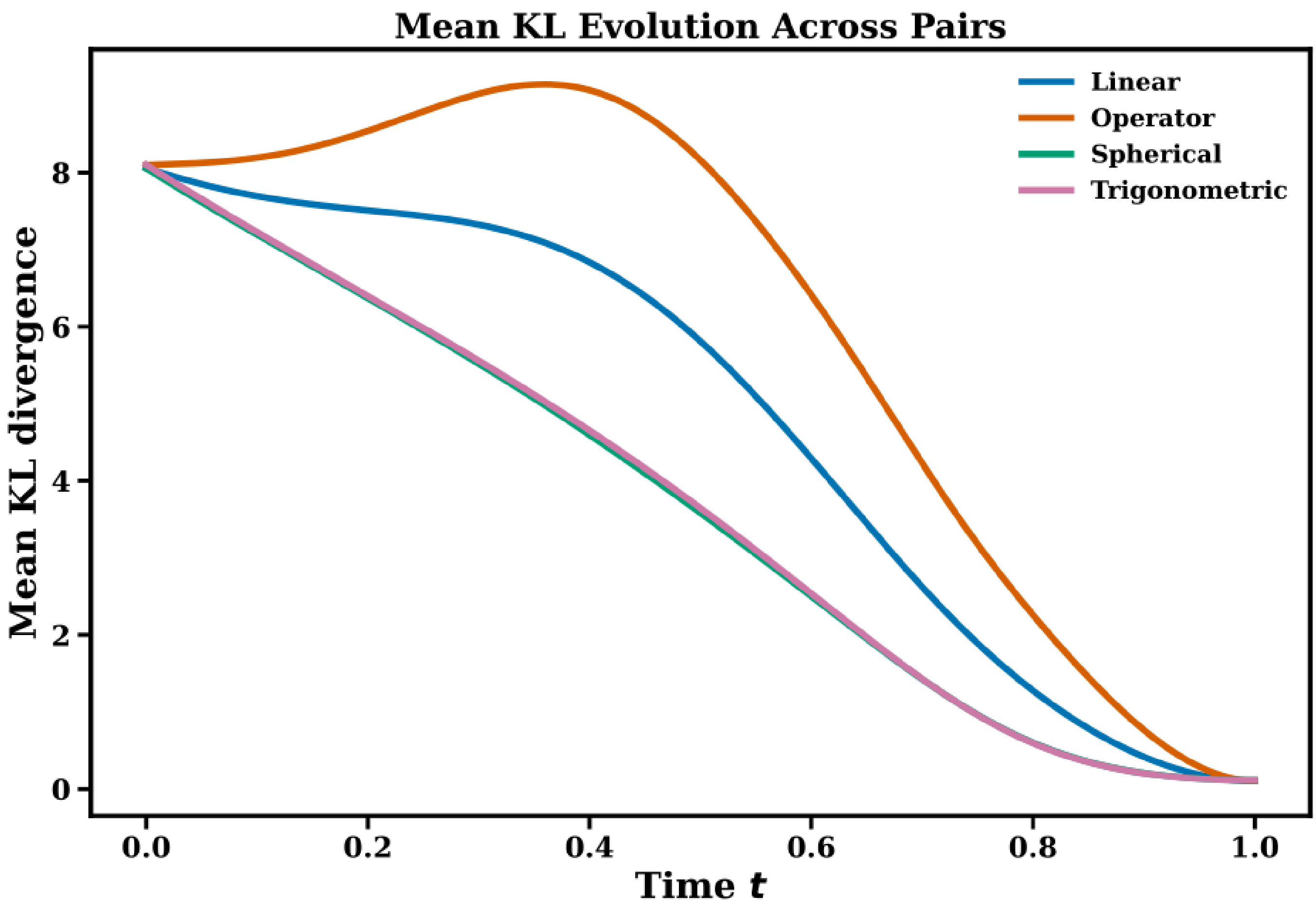


FIG. S29. **$\alpha$-Synuclein: marginal KL divergence averaged over the 30 $(\psi, \phi)$ pairs as a function of flow time.**

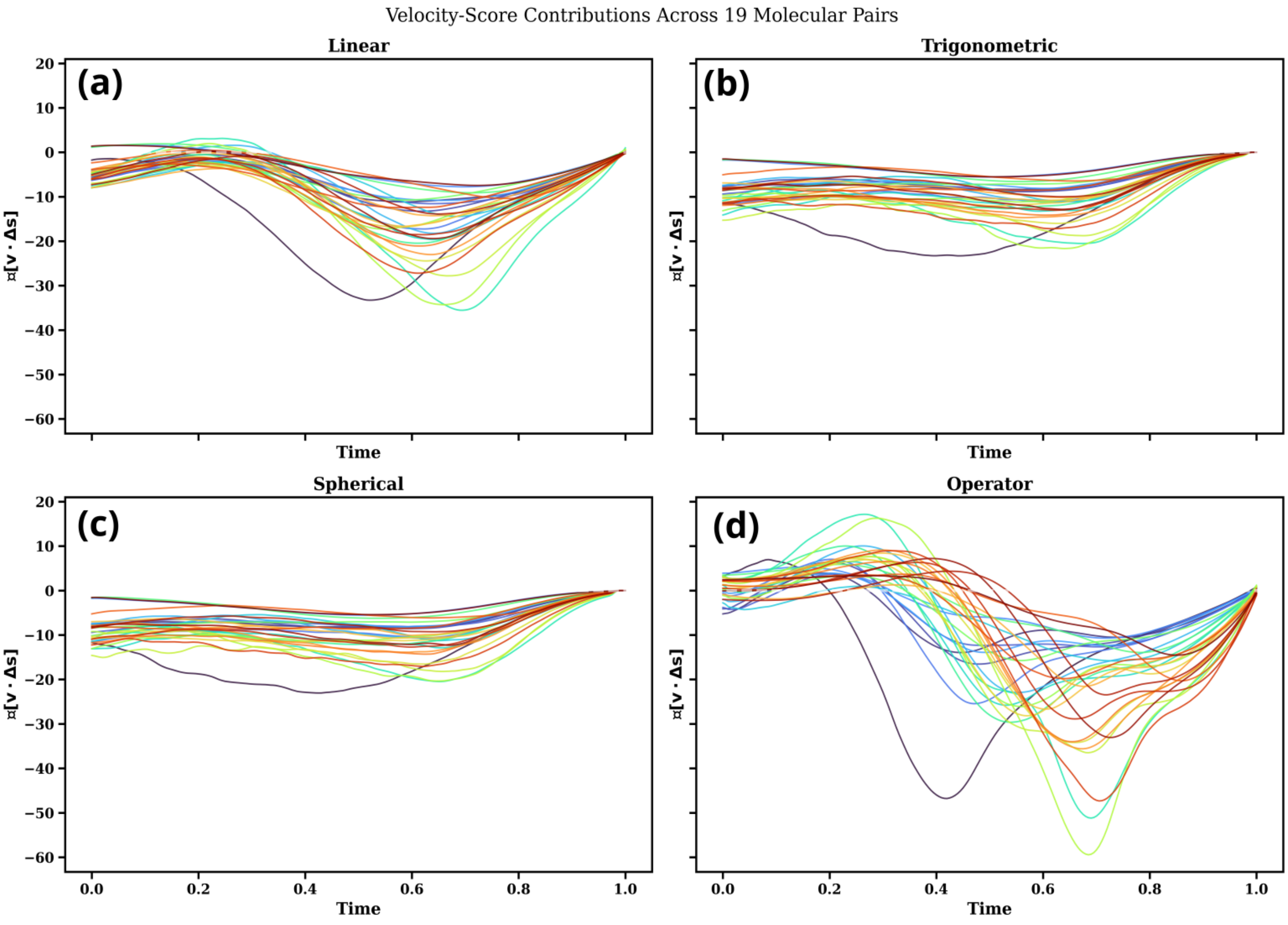


FIG. S30. $\alpha$-**Synuclein: velocity–score interaction $\mathbb{E}_{p_t}[\mathbf{v}_\theta \cdot \Delta \mathbf{s}_t]$ for the 30 $(\psi, \phi)$ pairs.**

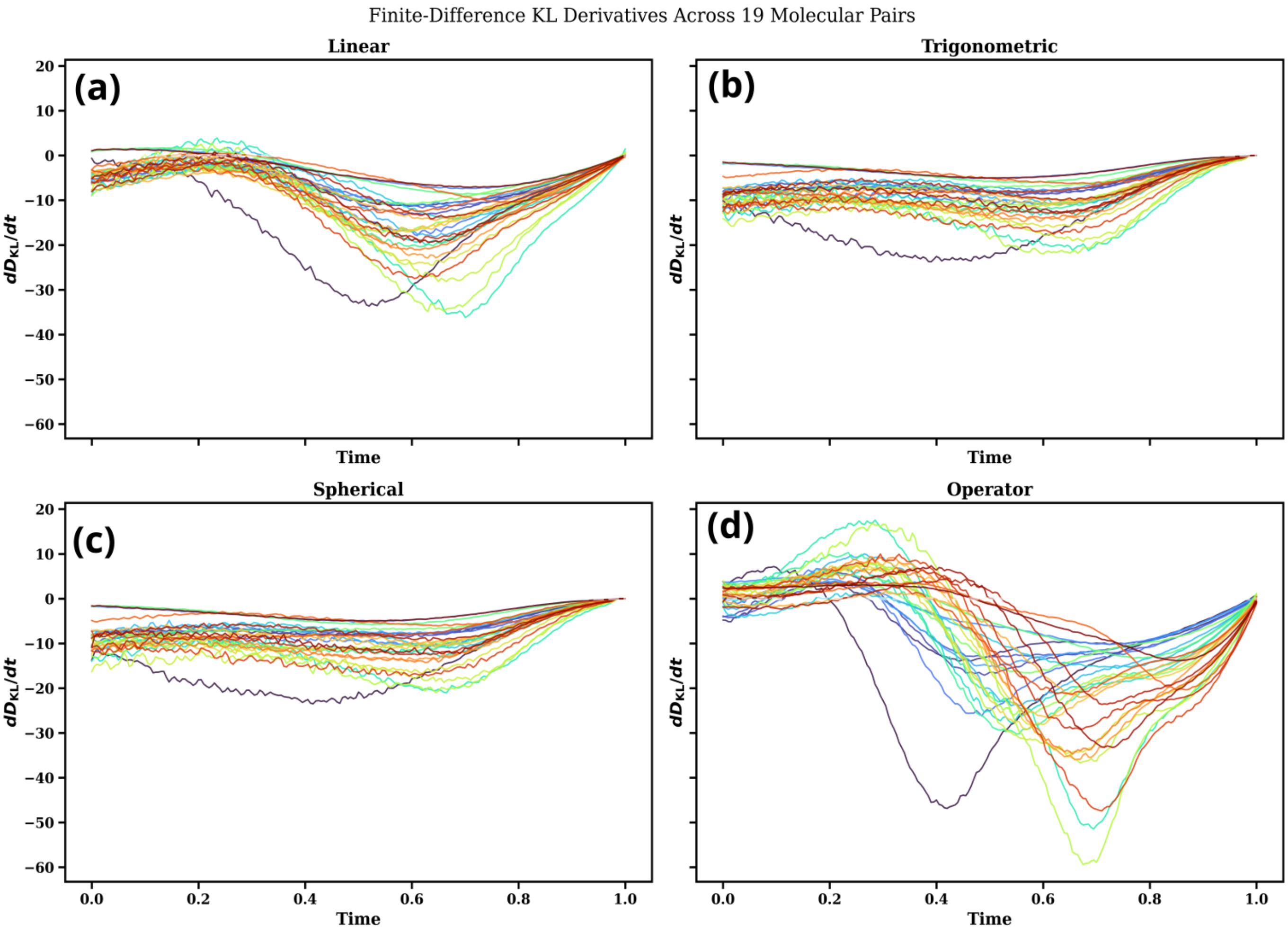


FIG. S31. **$\alpha$-Synuclein: finite-difference estimate of $dD/dt$ for the 30 $(\psi, \phi)$ pairs.**

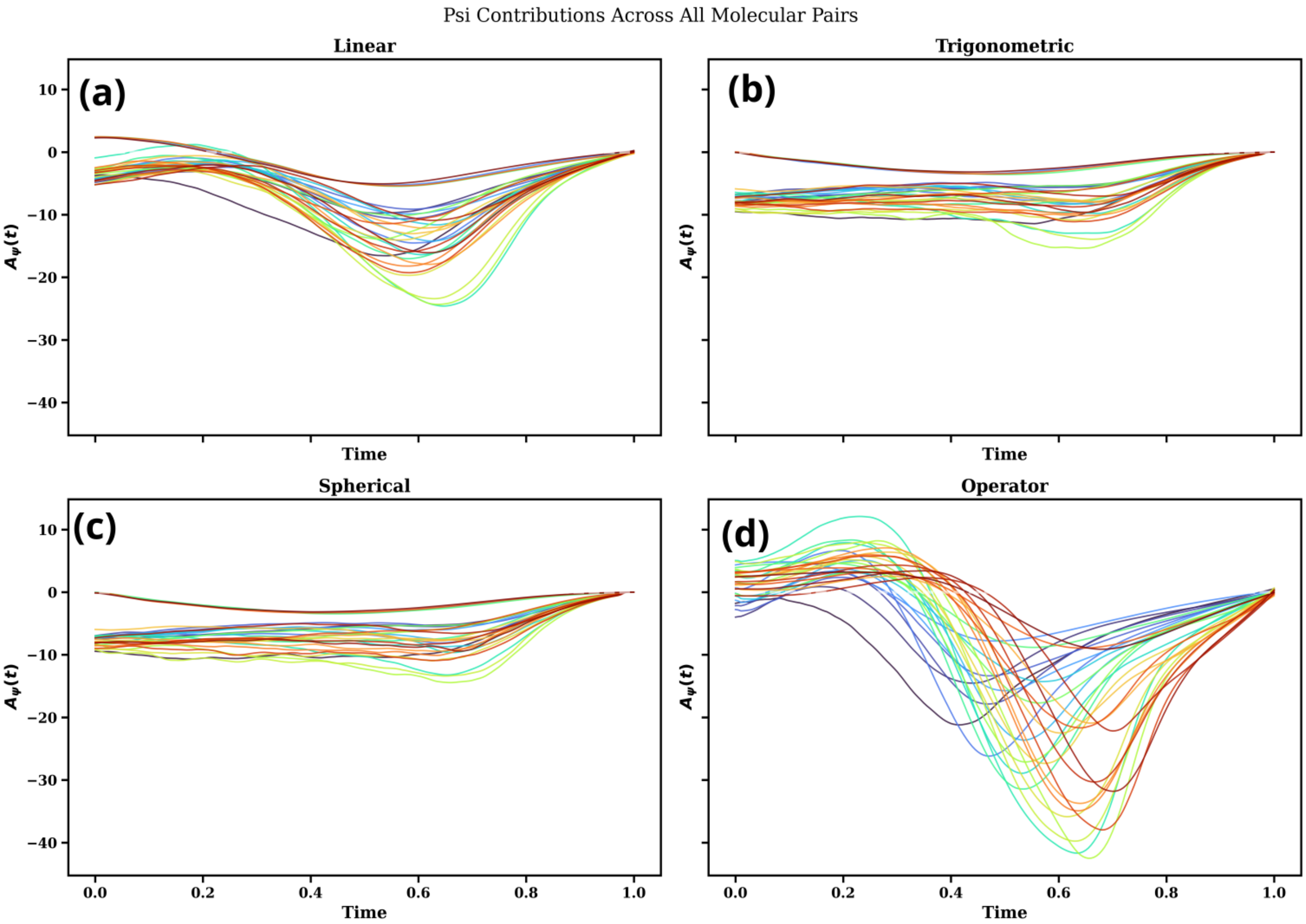


FIG. S32. $\alpha$**-Synuclein:** $\psi$ **contribution to the velocity–score interaction for the 30** $(\psi, \phi)$ **pairs.**

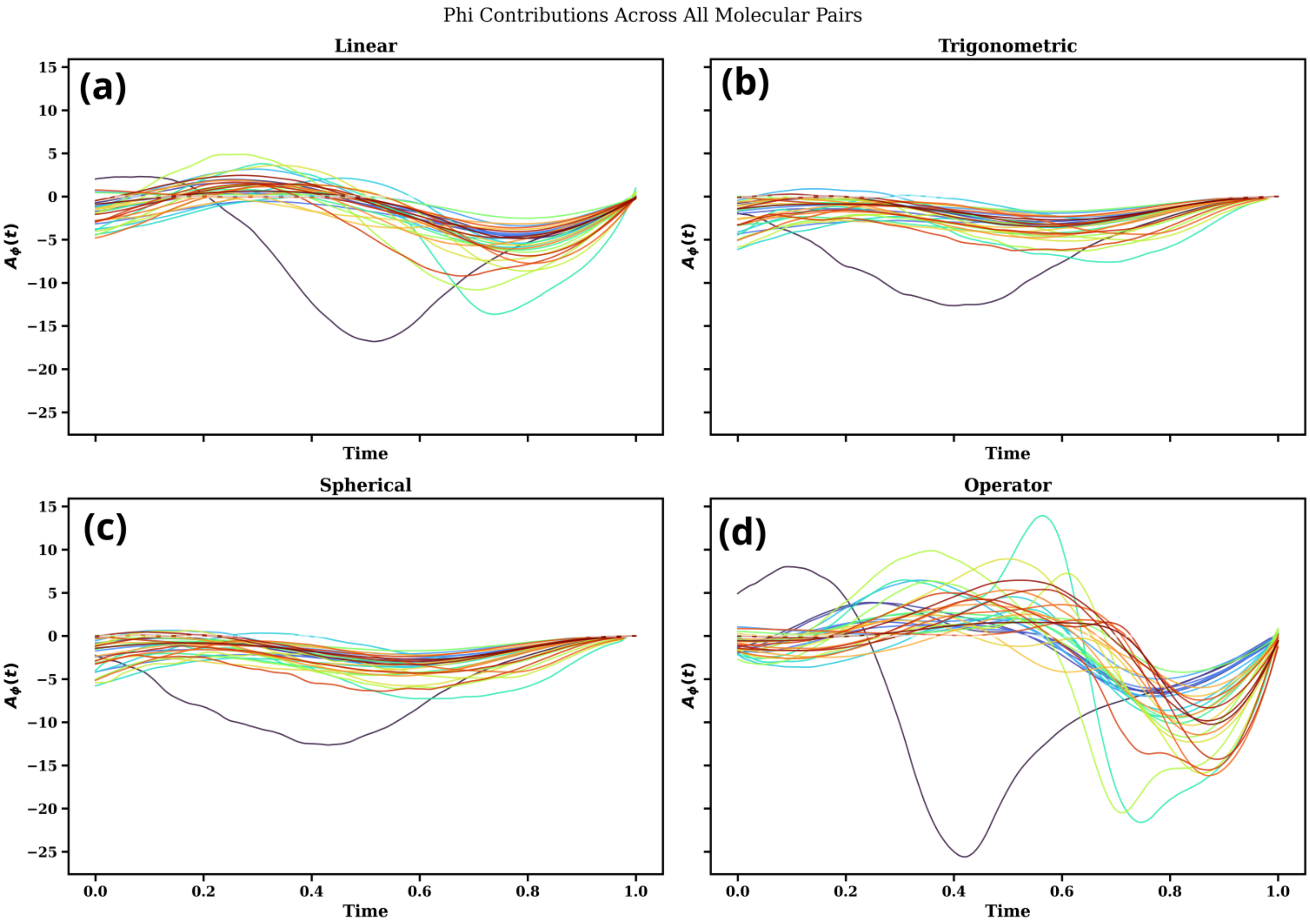


FIG. S33. $\alpha$**-Synuclein:** $\phi$ **contribution to the velocity–score interaction for the 30** $(\psi, \phi)$ **pairs.**

### C. Moment evolution: linear and operator-based paths

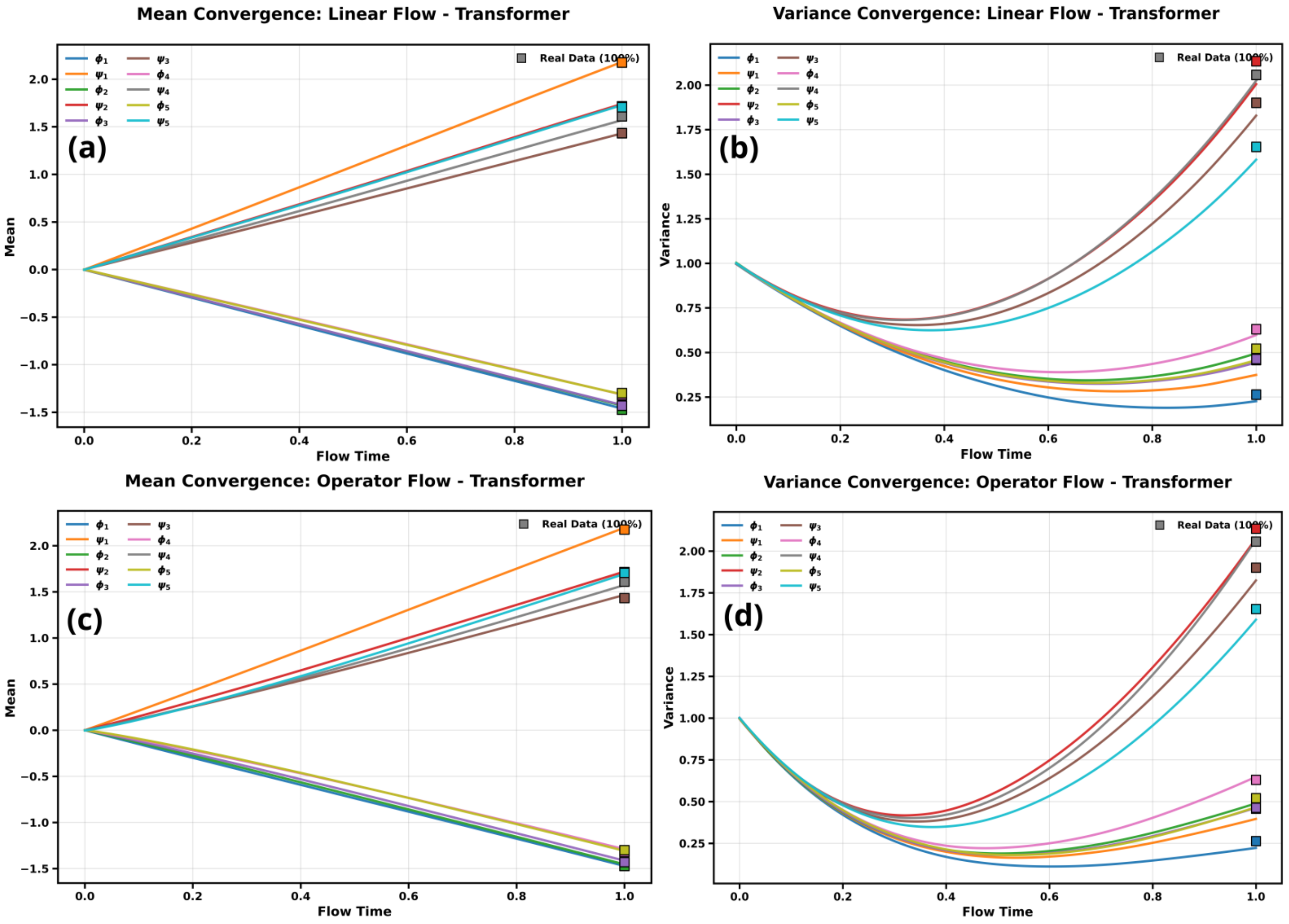


FIG. S34. **$\alpha$-Synuclein, pairs 1–5: mean and variance versus flow time for the linear (a,b) and operator-based (c,d) paths.** Squares mark the reference values.

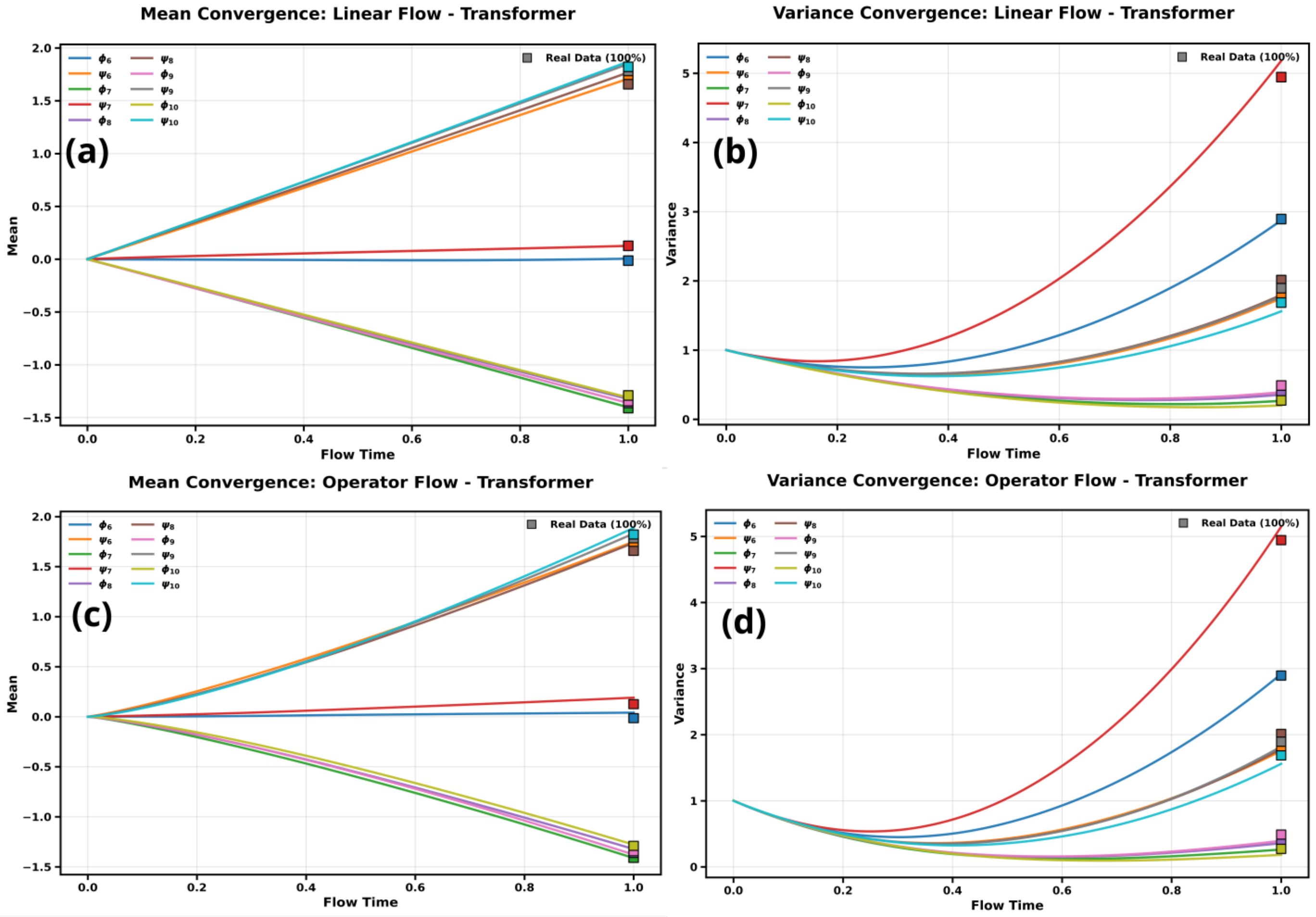


FIG. S35. $\alpha$**-Synuclein, pairs 6–10: linear and operator-based paths.** Panels as in Fig. S34.

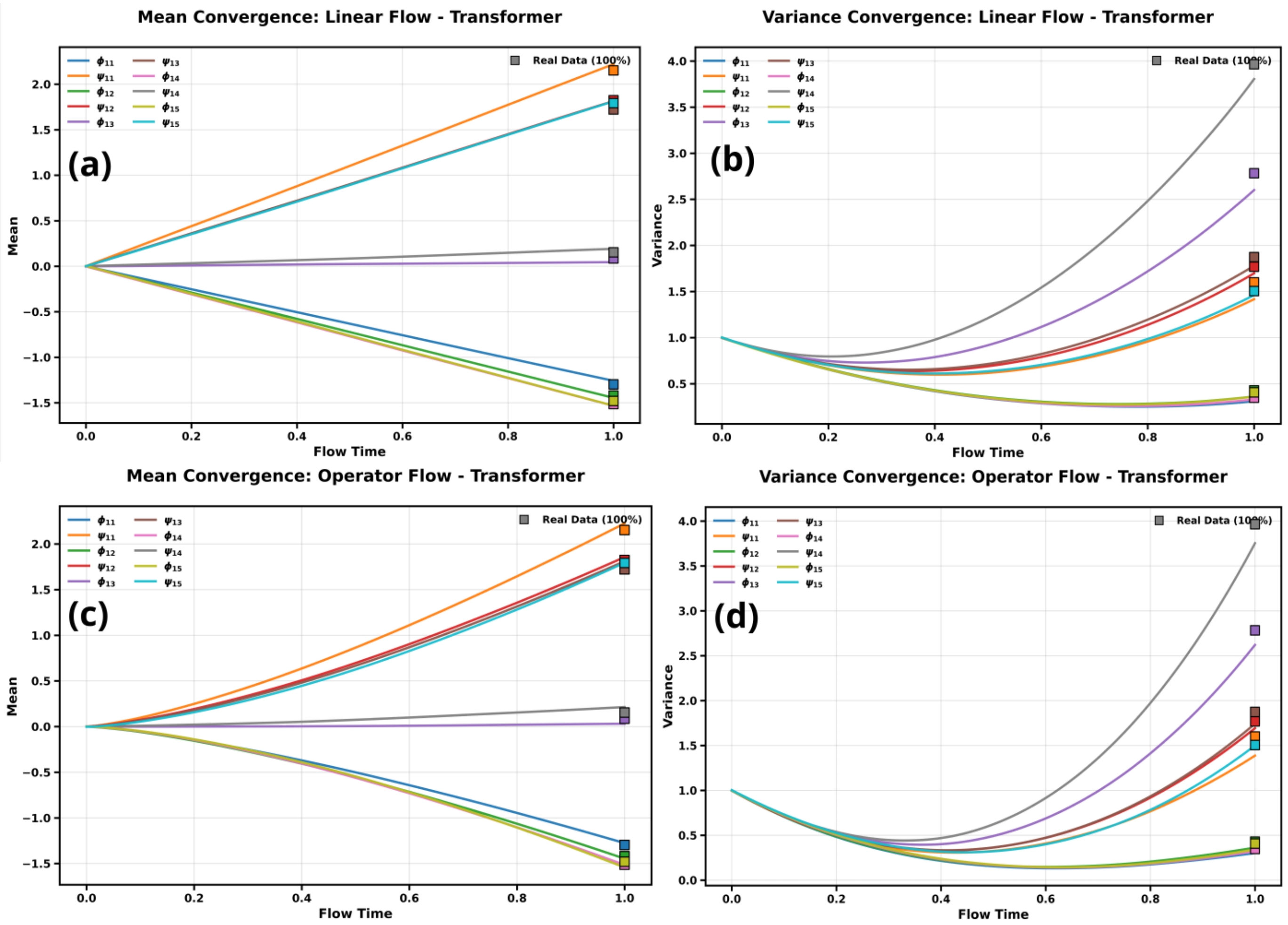


FIG. S36. **$\alpha$-Synuclein, pairs 11–15: linear and operator-based paths.** Panels as in Fig. S34.

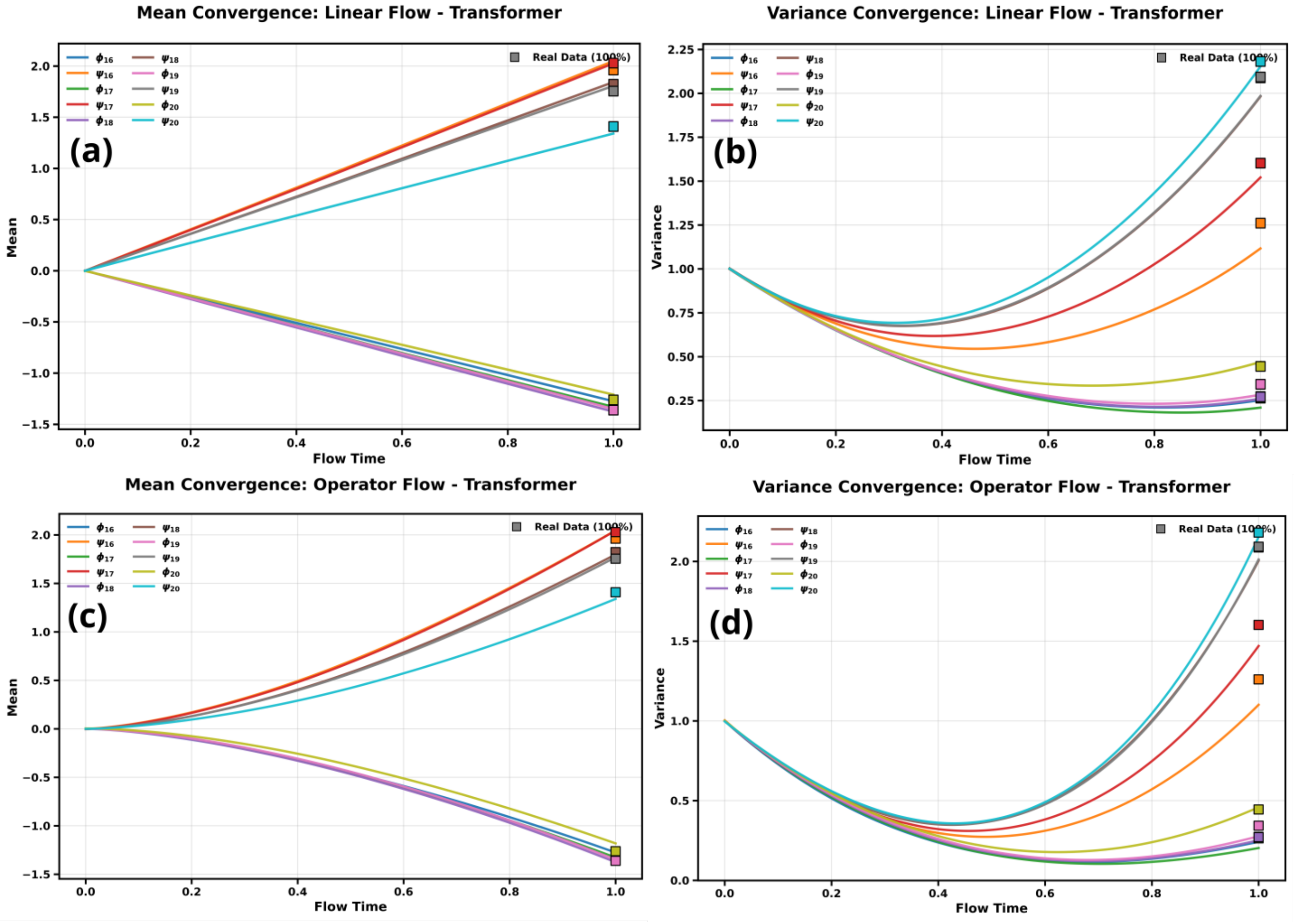


FIG. S37. **$\alpha$-Synuclein, pairs 16–20: linear and operator-based paths.** Panels as in Fig. S34.

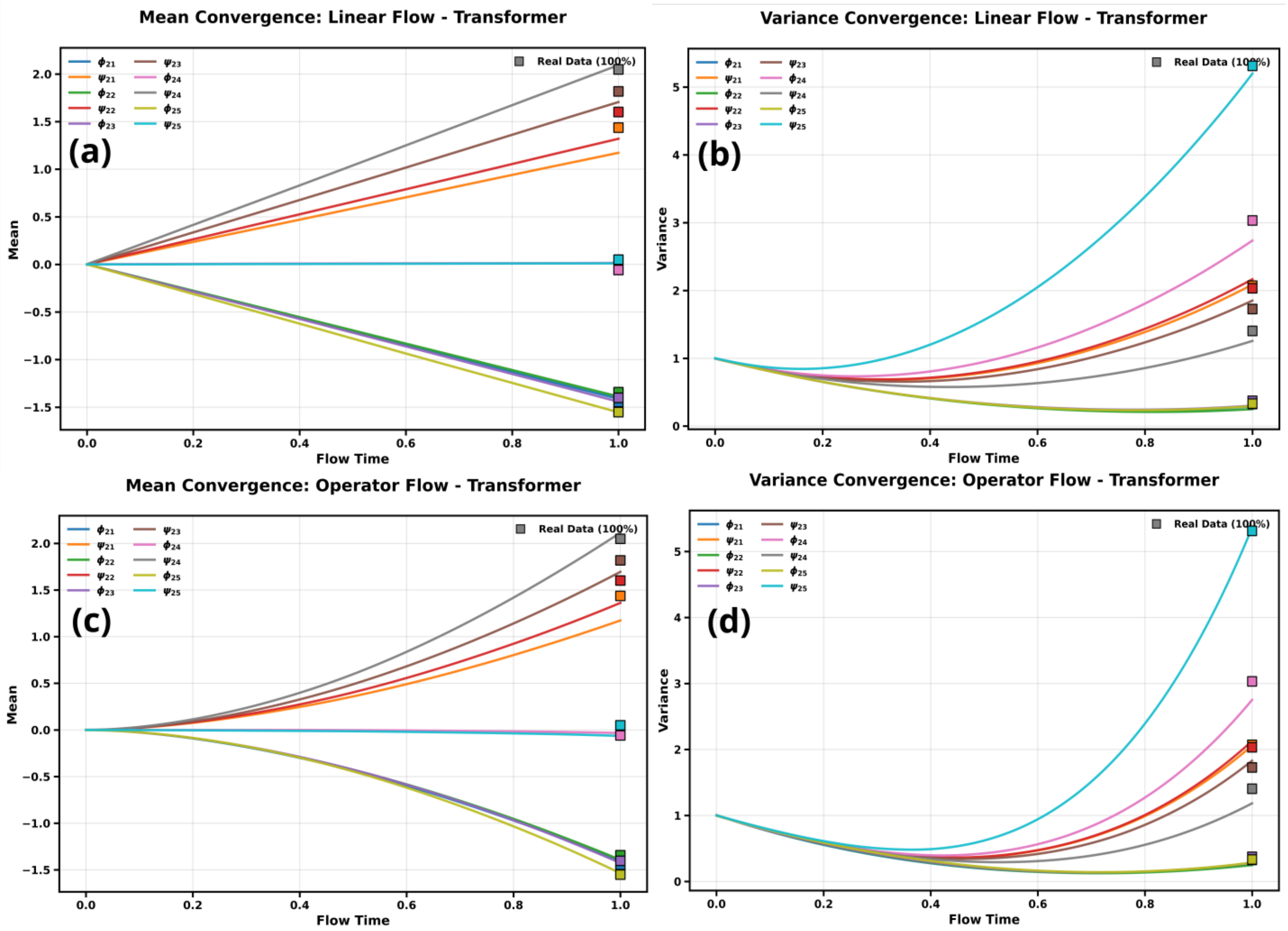


FIG. S38. **$\alpha$-Synuclein, pairs 21–25: linear and operator-based paths.** Panels as in Fig. S34.

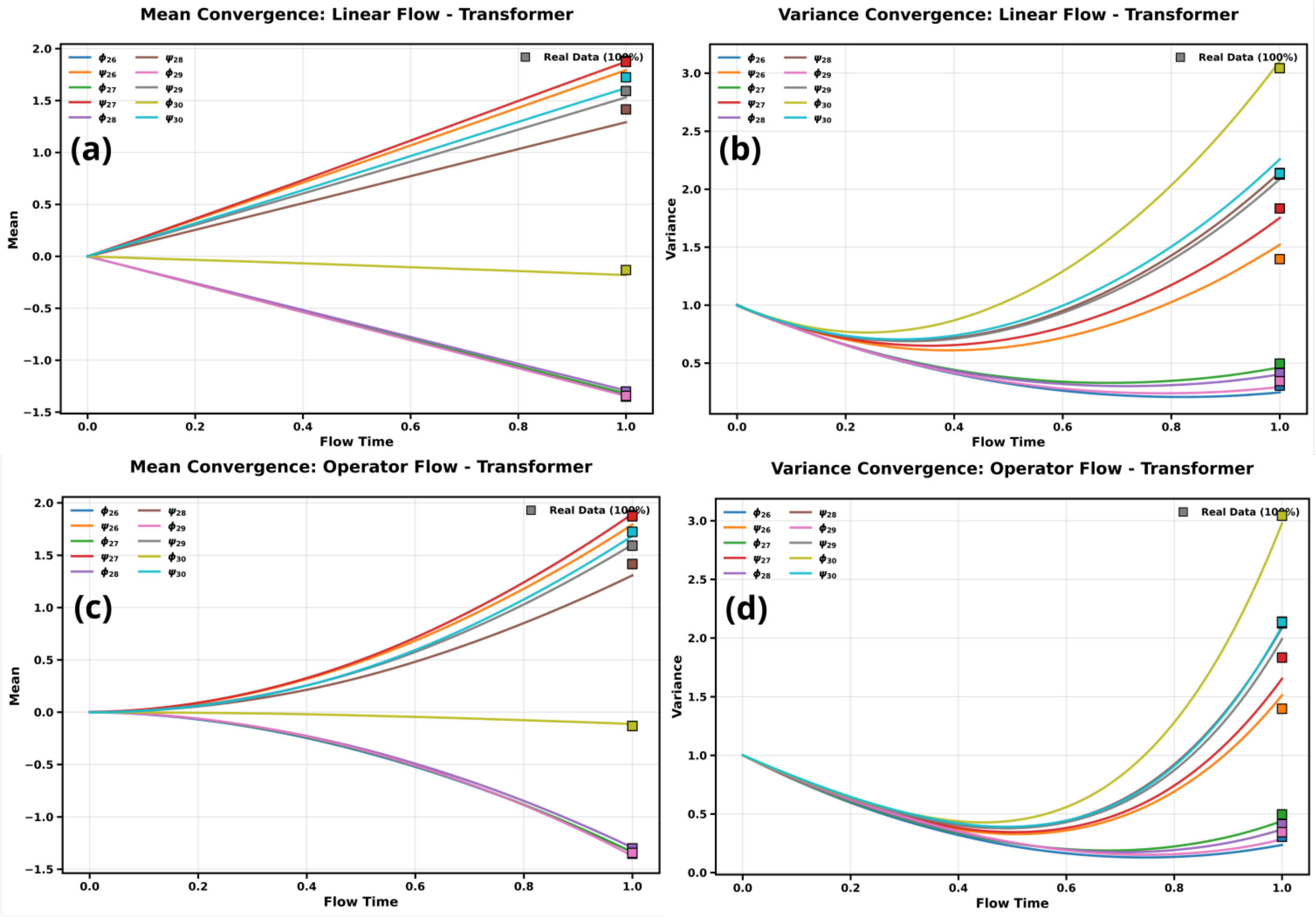


FIG. S39. **$\alpha$-Synuclein, pairs 26–30: linear and operator-based paths.** Panels as in Fig. S34.

### D. Moment evolution: trigonometric and spherical paths

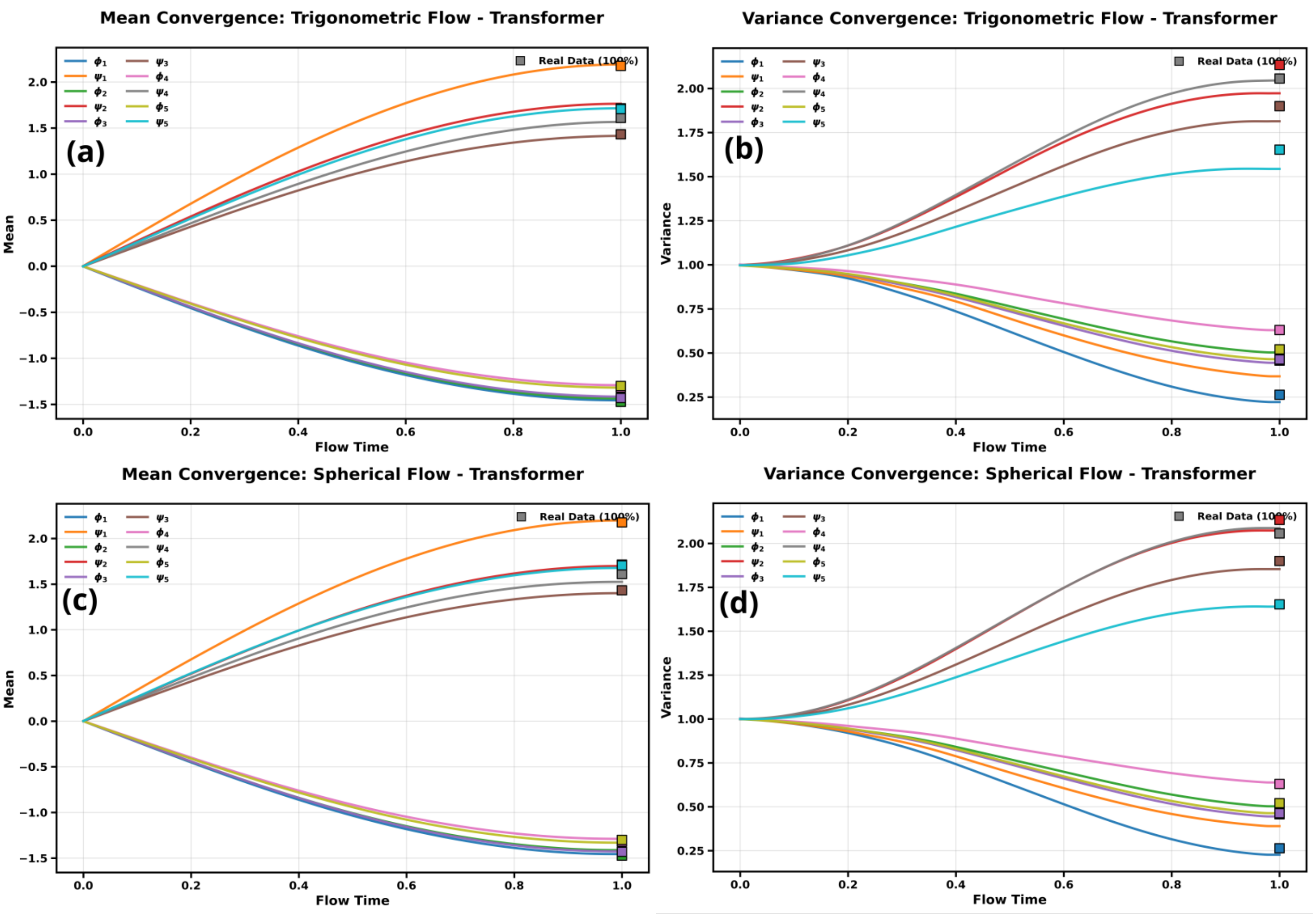


FIG. S40. $\alpha$**-Synuclein, pairs 1–5: mean and variance versus flow time for the trigonometric (a,b) and spherical (c,d) paths.** Squares mark the reference values.

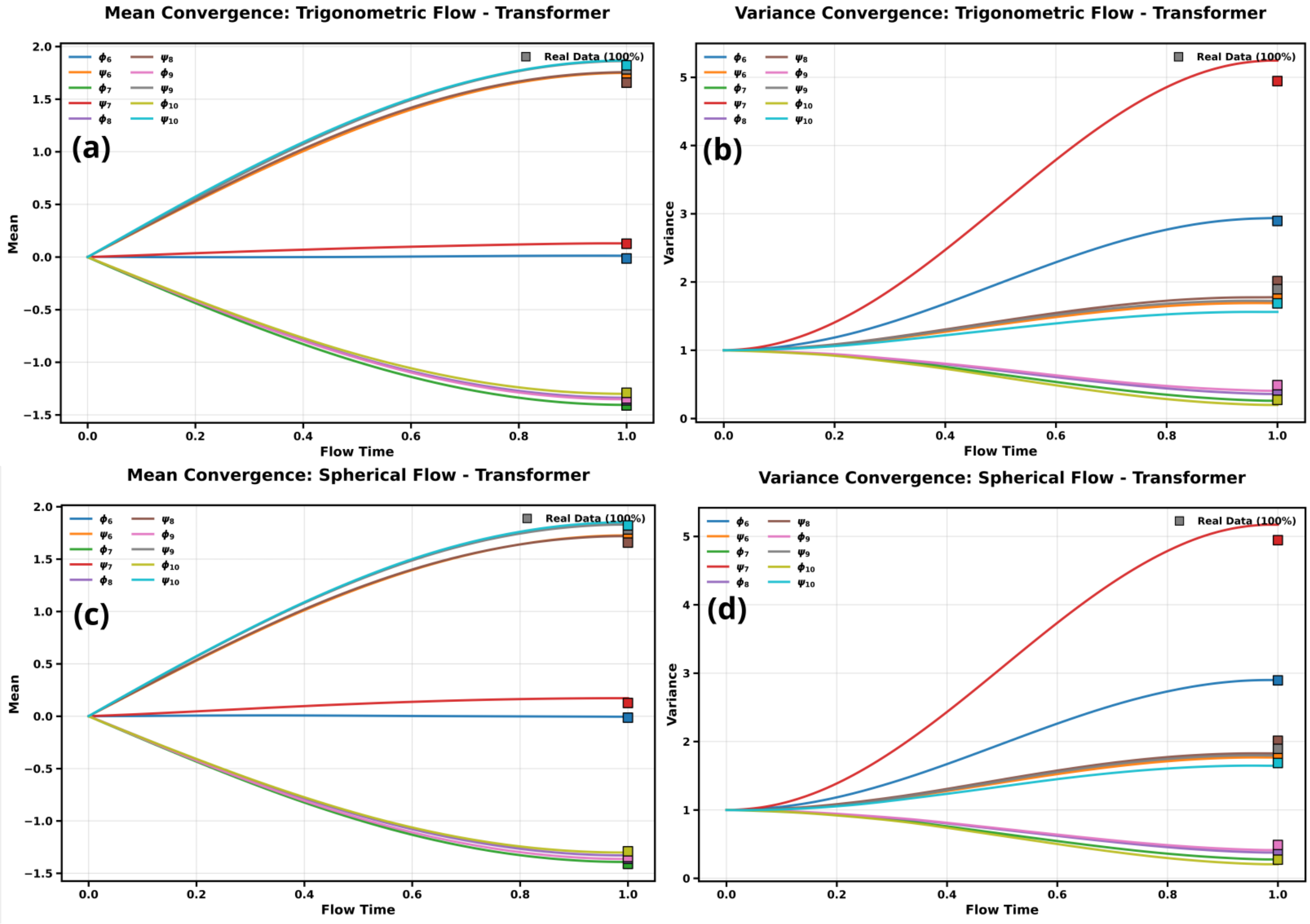


FIG. S41. **$\alpha$-Synuclein, pairs 6–10: trigonometric and spherical paths.** Panels as in Fig. S40.

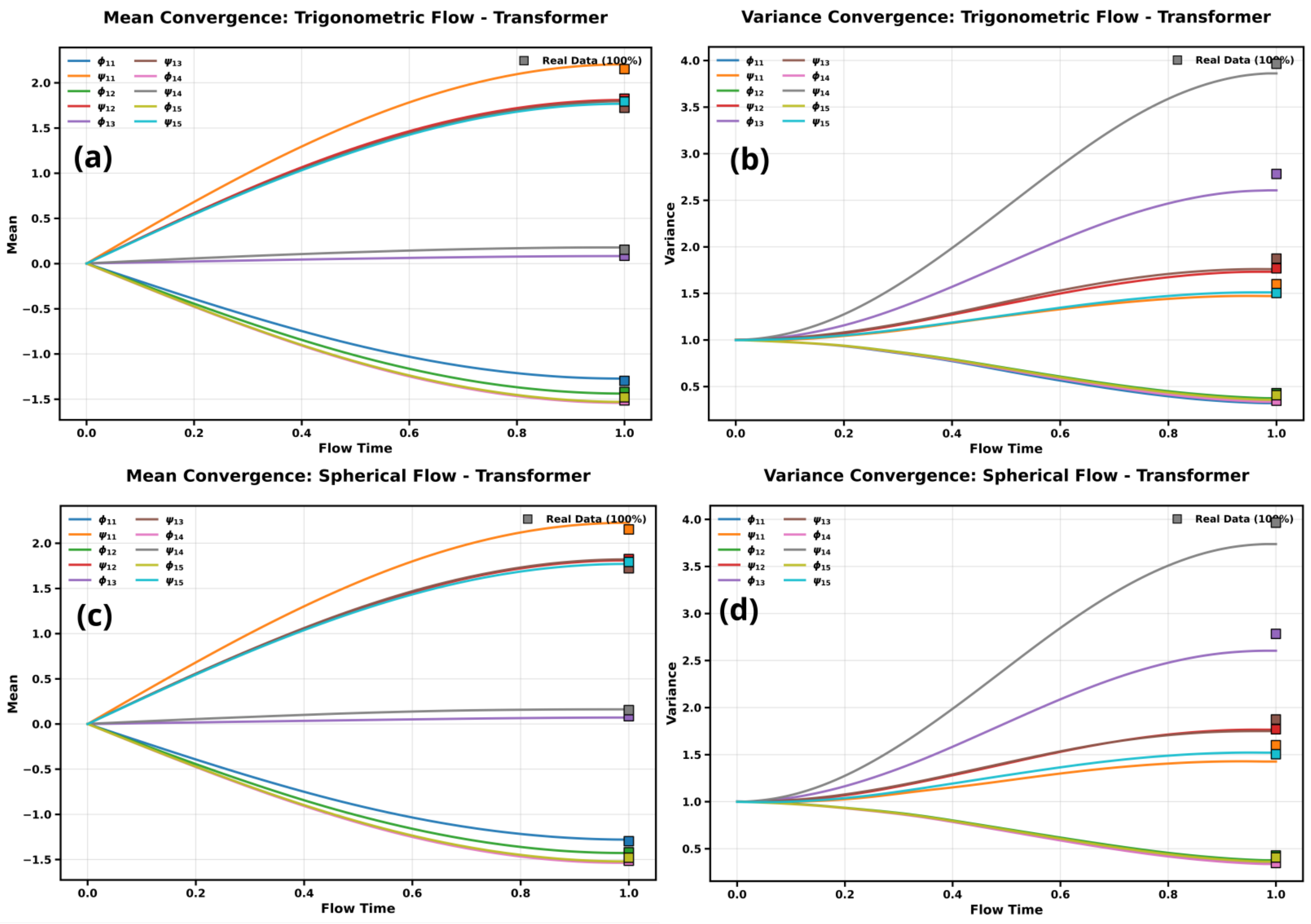


FIG. S42. **$\alpha$-Synuclein, pairs 11–15: trigonometric and spherical paths.** Panels as in Fig. S40.

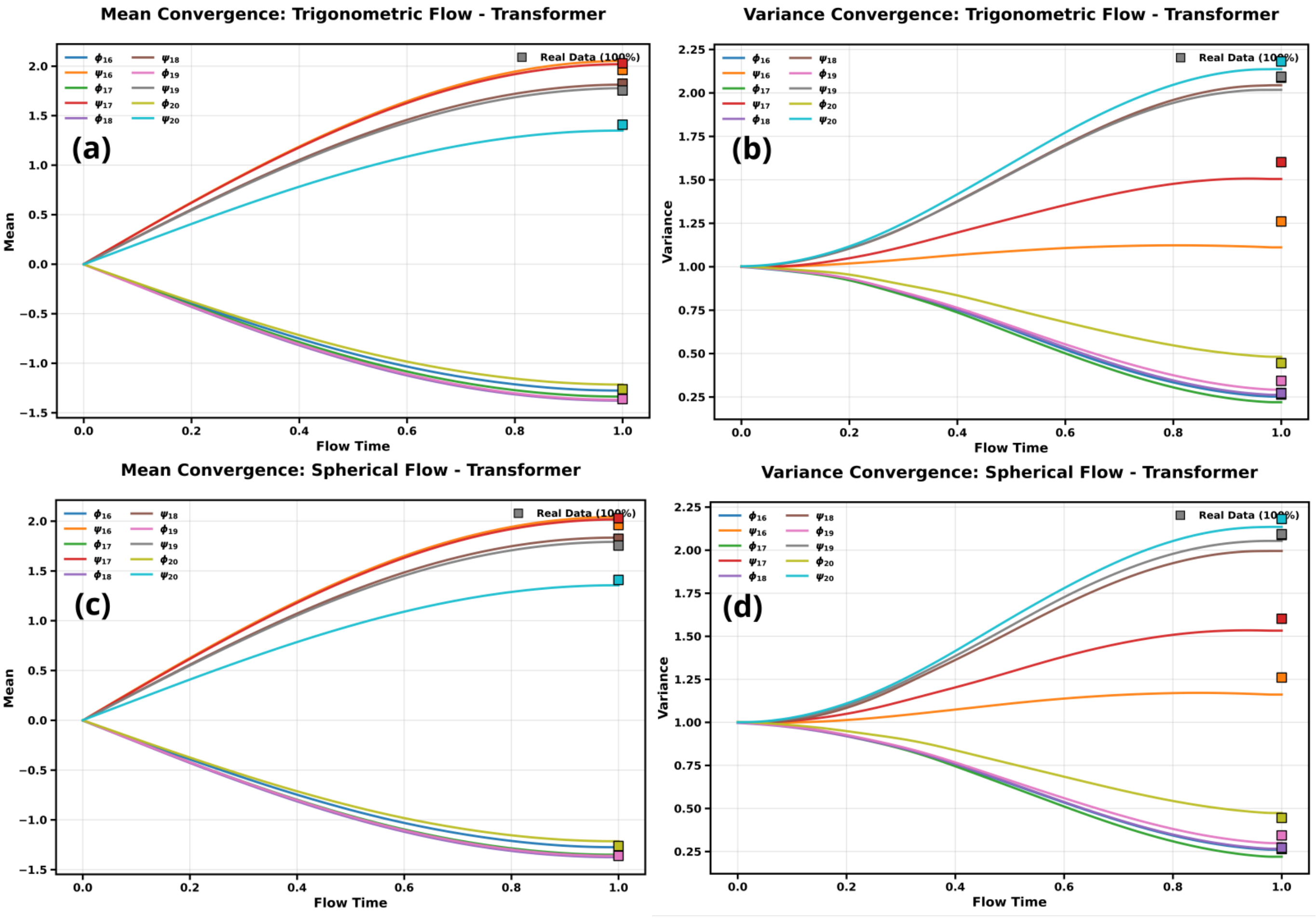


FIG. S43. **$\alpha$-Synuclein, pairs 16–20: trigonometric and spherical paths.** Panels as in Fig. S40.

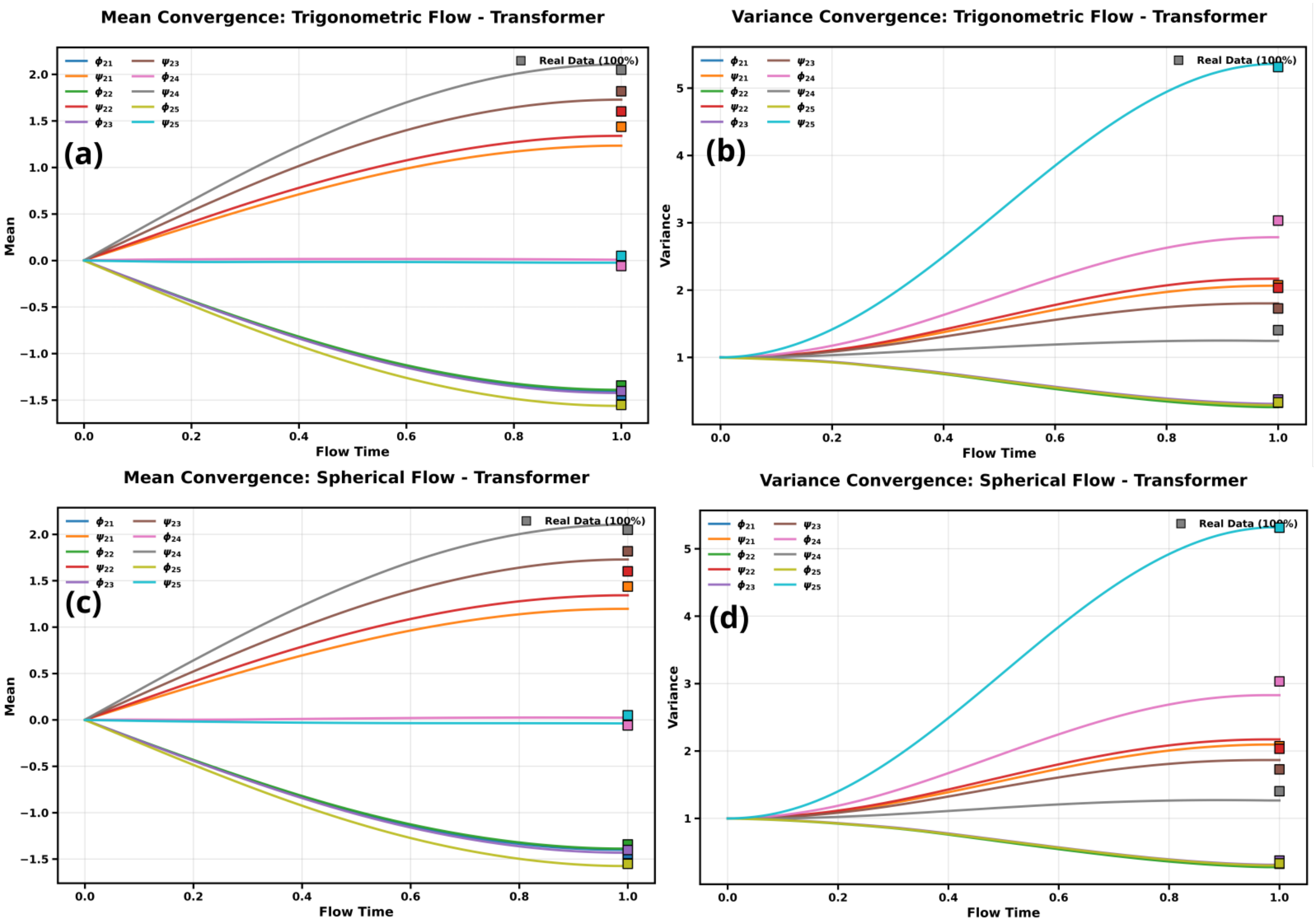


FIG. S44. **$\alpha$-Synuclein, pairs 21–25: trigonometric and spherical paths.** Panels as in Fig. S40.

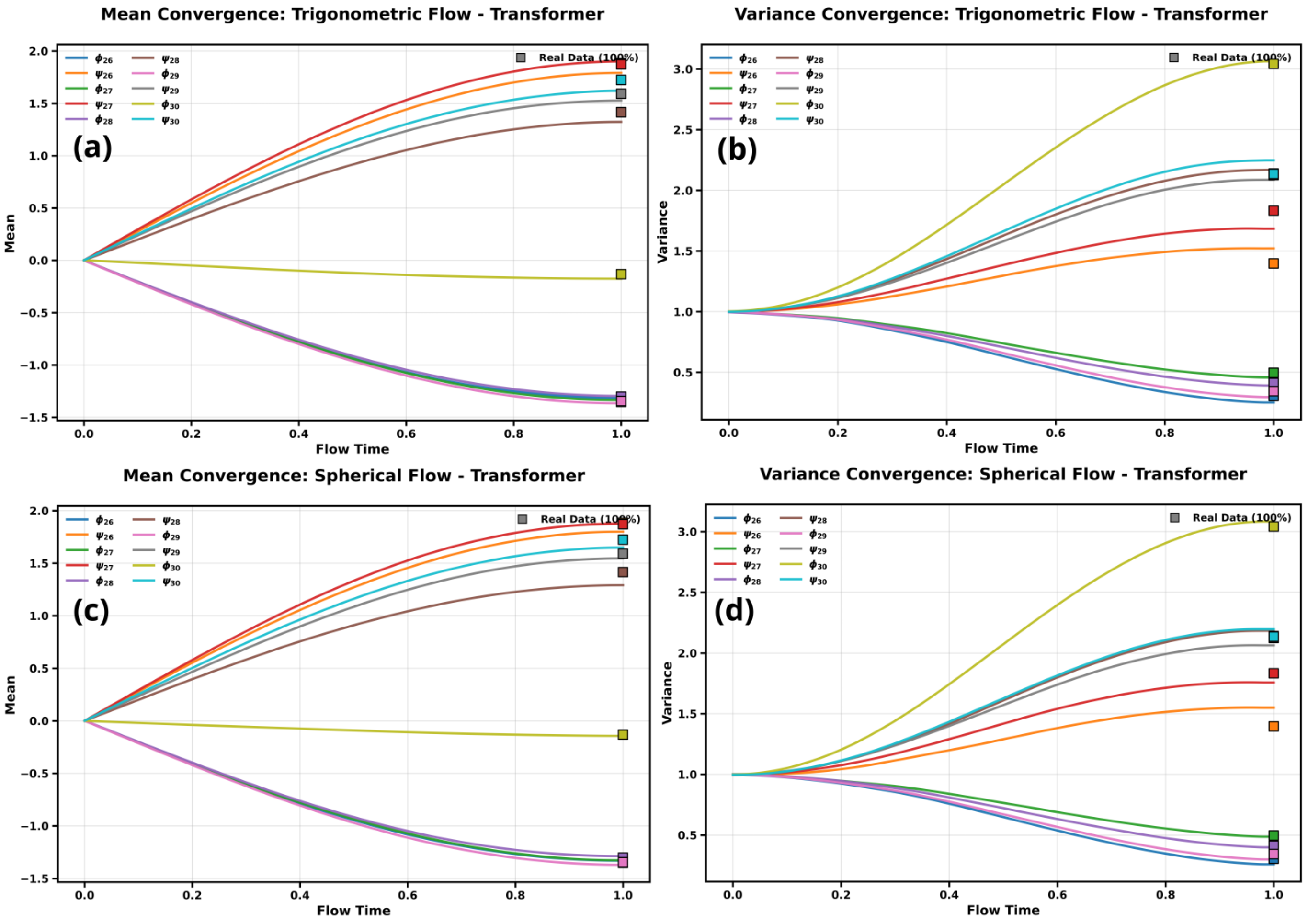


FIG. S45. **$\alpha$-Synuclein, pairs 26–30: trigonometric and spherical paths.** Panels as in Fig. S40.
